# iLoci: Robust evaluation of genome content and organization for provisional and mature genome assemblies

**DOI:** 10.1101/2021.10.03.462917

**Authors:** Daniel S Standage, Tim Lai, Volker P Brendel

## Abstract

**Background:** The rate at which new draft genome assemblies and corresponding annotation versions are being produced has long outpaced the scientific community’s capacity to refine these drafts into “finished,” reference-quality data resources to a standard typically expected from dedicated efforts of model organism research communities. Nonetheless, scientists must be able to evaluate newly sequenced genomes in the context of previously published data, requiring summaries of genome content and organization that can be quickly computed, updated, and meaningfully compared. As annotation quality will necessarily vary within and across data sets, the ability to select subsets of only those data that are well supported is critical for distinguishing technical artifacts from biological effects in genome-wide analyses.

**Results:** We introduce a new framework for genome analyses based on parsing an annotated genome assembly into distinct *interval loci* (*iLoci*), available as open source software as part of the AEGeAn Toolkit (https://github.com/BrendelGroup/AEGeAn). We demonstrate that iLoci provide an alternative coordinate system that is robust to changes in assembly and annotation versions and facilitates granular quality control of genome data. We discuss how statistics computed on iLoci reflect various characteristics of genome content and organization and illustrate how these statistics can be used to establish a baseline for assessment of the completeness and accuracy of the data. We also introduce a well-defined measure of relative genome compactness and compute other iLocus statistics that reveal genome-wide characteristics of gene arrangements in the whole genome context.

**Conclusions:** We present a coherent computational framework that calculates informative statistics from genome assembly/annotation data input. Given the fast pace of assembly/annotation updates, our AEGeAn Toolkit fills a niche in computational genomics based on deriving persistent and species-specific genome statistics. Gene structure model centric iLoci provide a precisely defined coordinate system that can be used to store assembly/annotation updates that reflect either stable or changed assessments. Large-scale application of the approach revealed species and clade specific genome organization in precisely defined computational terms, promising intriguing forays into the forces of shaping genome structure as more and more genome assemblies are being deposited.

## Background

The ready availability of Next-Generation Sequencing (NGS) technologies has resulted in genome data for thousands of species, with no slowing down of data accumulation in sight. Given this volume of data, fast and accurate computational approaches are needed now more than ever to process the initial sequence data into meaningful units of knowledge about the sequenced genomes. The conventional paradigm for such tasks from the early days of genome sequencing is outdated. At that time, one could expect community groups to carefully assemble and annotate the genomes of their expertise, resulting over a period of time in gap-filled assemblies and refined documentation of genome content in terms of protein-coding genes, products of alternative splicing, non-coding RNA (ncRNA) genes, transposable elements, repetitive sequences, and so forth. These genomes typically attained the status of “reference model genomes.” However, the time-consuming and expensive efforts required are impractical for the vast majority of organisms currently being sequenced with NGS technologies.

Out of necessity, the old paradigm has for the most part been replaced by an implicit new standard: genome data are presented as massive short read collections available from databases like the NCBI Sequence Read Archive [1] and in processed form as sets of assembled and computationally annotated scaffolds. Concomitantly, downstream analyses of these data have to be adjusted to scope- and quality-limitations intrinsic to the new data production process. First, assembly completeness will vary depending on the degree of read coverage and genome complexity (size and repetitiveness). Typically, assemblies will consist of tens to hundreds of large scaffolds, which in the best case can be ordered into linkage groups that approach pseudo-chromosomes, and additionally of manifold more short scaffolds, typically unplaced relative to any linkage groups. Second, annotation will commonly not have been expertly curated, but rather have resulted from first-pass outputs of annotation workflows such as AUGUSTUS [2], MAKER-P [3], BRAKER1 [4], or NCBI Gnomon [5].

The temporary nature of the data is also challenging. As additional sequences can often be acquired cheaply and easily for a species (for example, genomic DNA reads for libraries of different insert sizes; RNA-seq reads from transcriptome studies under various conditions; or spliced alignments of protein sequences from a newly annotated, closely related species), both the species’ genome assembly and its genome annotation may change. However, in the common scenario laid out above, the additional analyses will typically come without the community support to carefully sort out and document all the changes. Thus, over a short span of several years, there may be several annotation versions even for a single stable genome assembly, and it becomes difficult to track references to particular genes and genome features. A pertinent example from our experience is provided by the number of concurrent annotations in use for the honey bee (*Apis mellifera*) genome [6, 7, 8], including the current much more complete assembly based on long-read sequencing technologies [9].

How then should one compare results of a study on a current genome assembly and annotation version with previous results in the literature that used a prior assembly/annotation pair? How could one derive subsets of just those gene models that are solidly supported by evidence, to the extent that future genome-wide assembly/annotation improvements in all likelihood will not invalidate these current models? How does one disentangle artifacts of incomplete or inaccurate assembly/annotation from genuine species-specific genome features? What statistics should be calculated that capture a (newly sequenced) genome’s content and organization and allow meaningful comparison with other genomes?

A solution to the problem must address the dual issues of reproducibility and scalability to accommodate thousands of genomes, each potentially with multiple assemblies and annotations. At the core of a solution must be the ability to distinguish what has changed from what has remained invariant when comparing one assembly/annotation pair to another. Discriminating between solid, reliable annotations and annotations of uncertain quality is also crucial in order to enable separation of technical artifacts from effects of interest rooted in the underlying genome biology. Typical examples of this challenge include annotation of untranslated regions (UTRs), ncRNA genes, or identification of transposable elements: comparing two genome annotations, one would like to know whether differences in UTR lengths or ncRNA gene and transposon content are due to insufficient data for annotation, annotation workflow settings, or genome evolution.

Here we present our **AEGeAn** [10] (**A**nalysis and **E**valuation of **Ge**nome **An**notations) framework and toolset as a practical approach to facilitate comparisons across assemblies, annotations, and genomes in view of the described challenges. **AEGeAn** generalizes our previously published ParsEval software [11] for comparing two sets of annotations for the same genome assembly. The basic idea is to represent a given assembly/annotation pair as a set of distinct units that can be largely independently characterized and updated. We show how the parsing of a genome into such distinct *iLoci* provides a suitable “coordinate system” for working with rapidly changing genome assembly/annotation data. Applications to genome project data for various animal and plant species demonstrate how *iLoci* analyses can give insights into genome organization and features, as well as assembly and annotation status.

## Methods

### Toolkit scope and design

Motivated by the challenges of present day genome data reviewed in the **Background** section, we have developed a computational toolkit for the **A**nalysis and **E**valuation of **Ge**nome **An**notations (**AEGeAn** [10]). AEGeAn includes functions that address questions of genome content, genome organization, and cross-genome comparisons by precisely defined measures. The first range of questions concerning genome content include: How many genes are annotated for a particular assembly/annotation pair? What can be said about their length, number of exons, nucleotide composition, and other characteristics? What proportion of the genome is occupied by these genes? What fraction of genes are protein-coding versus ncRNA genes? How many of the gene models have support from transcript evidence, and how many genes can be identified as likely homologs of genes in other species?

These seemingly simple questions actually require very precise processing of the annotation file to be reproducibly and meaningfully answered. In particular, the handling of alternative transcription as well as overlapping gene models needs to be unambiguously defined.

The second range of questions concerning genome organization include: How densely or sparsely packed are the genes? Is there clustering of genes, and if so, how large are these clusters, and what types of genes occur in clusters (e.g., [12])? More generally, how is the intergenic space organized?

Thirdly, all of the above questions are of interest in a comparative genomics context (e.g., [13]). To what extent are genomes within a clade of species similarly organized? And, maybe even more intriguingly, to what extent is genome organization functionally important?

The design of the toolkit followed bioinformatics software engineering principles that emphasize reproducible, scalable, and extensible open source code that is easy to use and integrates with existing data repositories such as NCBI Genome [14] and other toolkits such as GenomeTools [15, 16]. Minimal required data input consists of a triplet of files (*G, A, P*): *G* is a set of of one or more genome sequences provided in multi-FASTA formats; *A* is the associated genome annotation provided in GFF3 [17] format; and *P* is the set of annotated protein-coding gene products, in multi-FASTA format. For most sequenced genomes, such files are readily accessible at NCBI Genome [14]. For simplicity, genome annotation provided in other formats would have to be converted to GFF3 input using widely available third-party scripts. In most cases, the protein file *P* could be generated from the CDS annotation in the GFF3 file. However, the more general specification of a separate *P* file accounts for non-templated gene products that may be cited in the annotation file. AEGeAn includes format-checking utilities that flag semantic inconsistencies in the input and suggest GenomeTools functions to remedy identified problems.

### Conceptual definition of interval loci

To address the toolkit design prescriptions, we introduce a precise parsing of an assembly/annotation pair into smaller units, termed *interval loci*, that provide a robust, granular, and dynamic strategy for answering the biological questions posed above. Each interval locus (or *iLocus*) is intended to capture the local genomic context of a genic or intergenic space, providing an alternative coordinate system to the conventional scaffold-based system; an alternative that is substantially more robust to changes in assemblies and annotations. Conceptually, an iLocus is a genomic interval, the boundaries of which are computed from annotated gene models, with an extension to include probable adjacent *cis*-regulatory regions. The precise procedure for computing iLoci is described in detail in the next section.

iLoci can be distinguished by various characteristics, as summarized in **Figure 1**. iLoci containing genes are referred to as *giLoci*, with those encoding protein-coding genes labeled as *piLoci* and those containing non-coding genes labeled as *niLoci*. piLoci harboring multiple overlapping gene models are designated complex (*ciLoci*), while those with a single isolated gene model are designated simple (*siLoci*). iLoci containing no gene models are designated as intergenic (*iiLoci*) if they are flanked on both sides by genes, or as incomplete fragments (*fiLoci*) if they are are flanked on at least one side by an end of the corresponding parsed sequence.

**Figure 1:**
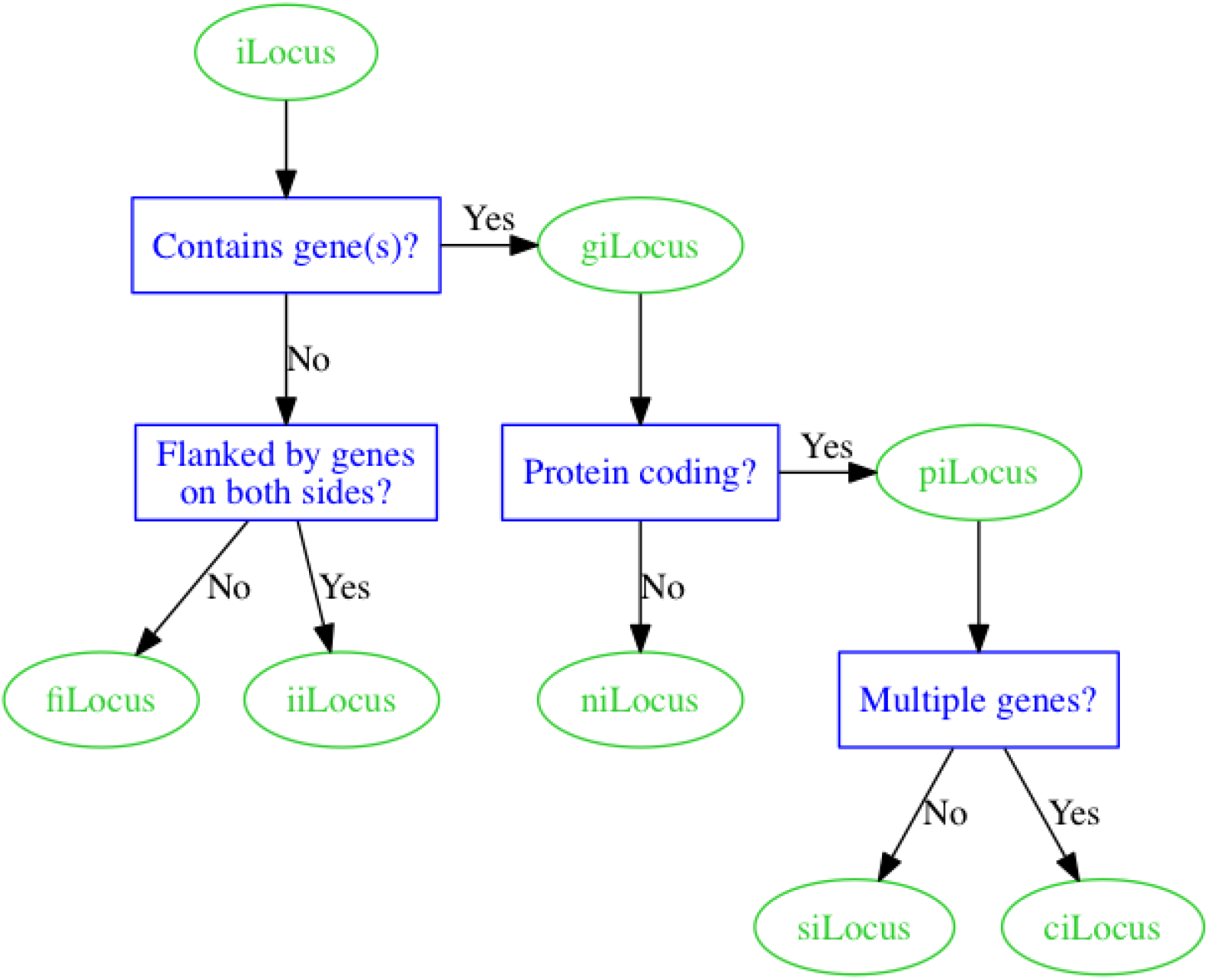
Classification of iLoci. Designation of iLocus types is shown in green, with classification logic described in blue. Abbreviations: fiLocus, fragmented intergenic iLocus; iiLocus, complete intergenic iLocus; giLocus, genic iLocus; niLocus, non-coding gene-containing giLocus; piLocus, protein-coding gene-containing giLocus; siLocus, simple piLocus; ciLocus, complex piLocus.

To illustrate these concepts, **Figure 2** shows the parsing of a hypothetical scaffold into its constituent iLoci. The parsing captures an intuitive and practical decomposition of the genome. The piLoci comprise a non-redundant set of protein-coding genes when reporting gene number or calculating descriptive statistics on gene features. However, more reliable results would be expected from the siLoci, or even better a subset of the siLoci with well supported gene models. The ciLoci will typically require a whole lot more attention in order to establish whether the overlapping gene models reflect observed transcription or are artifacts of unresolved annotation conflicts.

**Figure 2:**
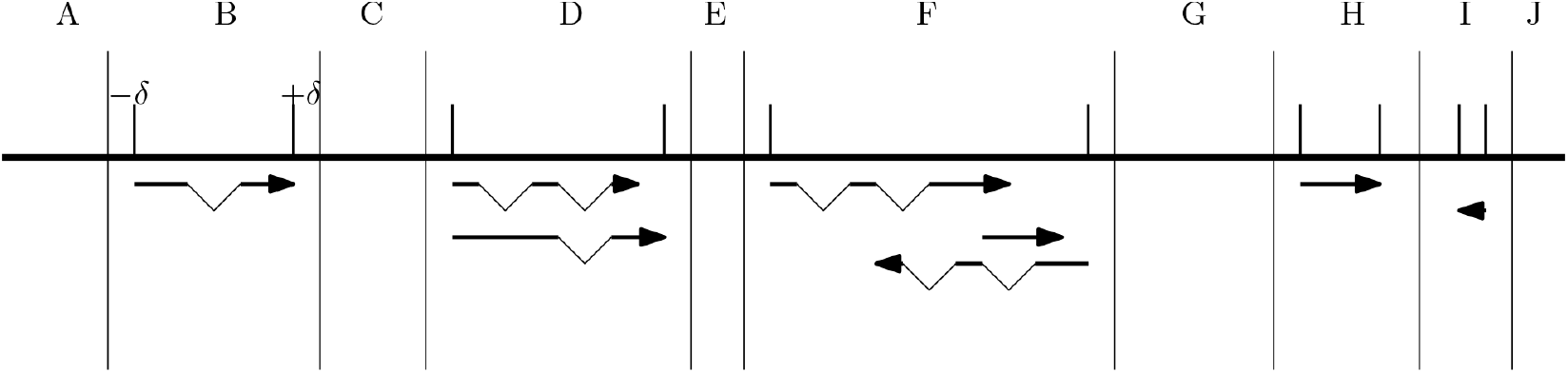
Parsing an annotated genome sequence into iLoci. The letters A to J indicate 10 adjacent iLoci on the genomic sequence (central horizontal line), separated by the long vertical bars. Gene annotations are shown underneath the genome sequence. Exons are schematized by bold horizontal lines and introns by the triangular thin lines connecting exons. Arrows indicate transcriptional direction. iLoci A, C, E, G, and J are without gene annotation, with A and J representing potentially incomplete genomic fragments (fiLoci), and C, E, and G representing complete intergenic regions (iiLoci). Each siLoci contains annotation for a single gene, which may involve a unique transcript (B, H, and I) or multiple alternative transcripts (D). ciLocus F contains three distinct, but overlapping genes. The boundaries of the gene-containing iLoci (giLoci) are derived from the annotation ends, extended in each direction by *δ*. An exception occurs between giLoci H and I, where the extension would result in an iiLocus shorter than *δ*: in this case, the bordering giLoci (H and I) are extended towards each other to fill the entire space.

### Operational definition of iLoci

#### Basic procedure

Computing iLoci for an assembled contig/scaffold/pseudo-chromosome *S* depends on a set of intervals *G* (corresponding to gene models annotated on *S*) and an extension parameter *δ* (default value: 500). The basic procedure is described in **Algorithms 1** and **2**. In brief, the ComputeLoci algorithm computes a set of intervals *L* such that any two overlapping elements *g*_*m*_, *g*_*n*_ *∈ G* are bounded by the same interval *loc ∈ L*. Although the algorithm is general, here *g*_*m*_ and *g*_*n*_ refer to gene bodies, defined as the interval from the start to the end of the respective annotated transcription events. The ExtendIntervals algorithm then assesses each pair of adjacent intervals *loc*_*m*_, *loc*_*n*_ *∈ L* and determines how far the intervals can be extended toward each other and whether any additional space remains between them for the creation of a third interval: if the number of nucleotides separating the two intervals *dist*(*loc*_*m*_, *loc*_*n*_) *>* 3*δ* nucleotides, then *loc*_*m*_ and *loc*_*n*_ will be extended toward each other by *δ* nucleotides, each designated as a giLocus, and the remaining space between them will be designated as an iiLocus; if 2*δ < dist*(*loc*_*m*_, *loc*_*n*_) *≤* 3*δ*, then *loc*_*m*_ and *loc*_*n*_ are extended toward each other equally until they meet, with extensions potentially as long as 1.5*δ*, to prevent recording a short iiLocus of positive length *≤ δ*; if *dist*(*loc*_*m*_, *loc*_*n*_) *≤* 2*δ, loc*_*m*_ and *loc*_*n*_ will each be extended by *δ* resulting in slightly overlapping iLoci. The rationale for allowing iLoci boundary overlaps in these cases is to assure that any giLoci selected for inspection will have *δ* nucleotide flanks around the transcript-based gene annotation. In both cases where *dist*(*loc*_*m*_, *loc*_*n*_) *≤* 3*δ*, the toolkit records a zero-length iLocus (*ziLocus*) between the adjacent giLoci for consistency and calculation of cumulative statistics described below.

##### Algorithm 1 Compute giLocus boundaries

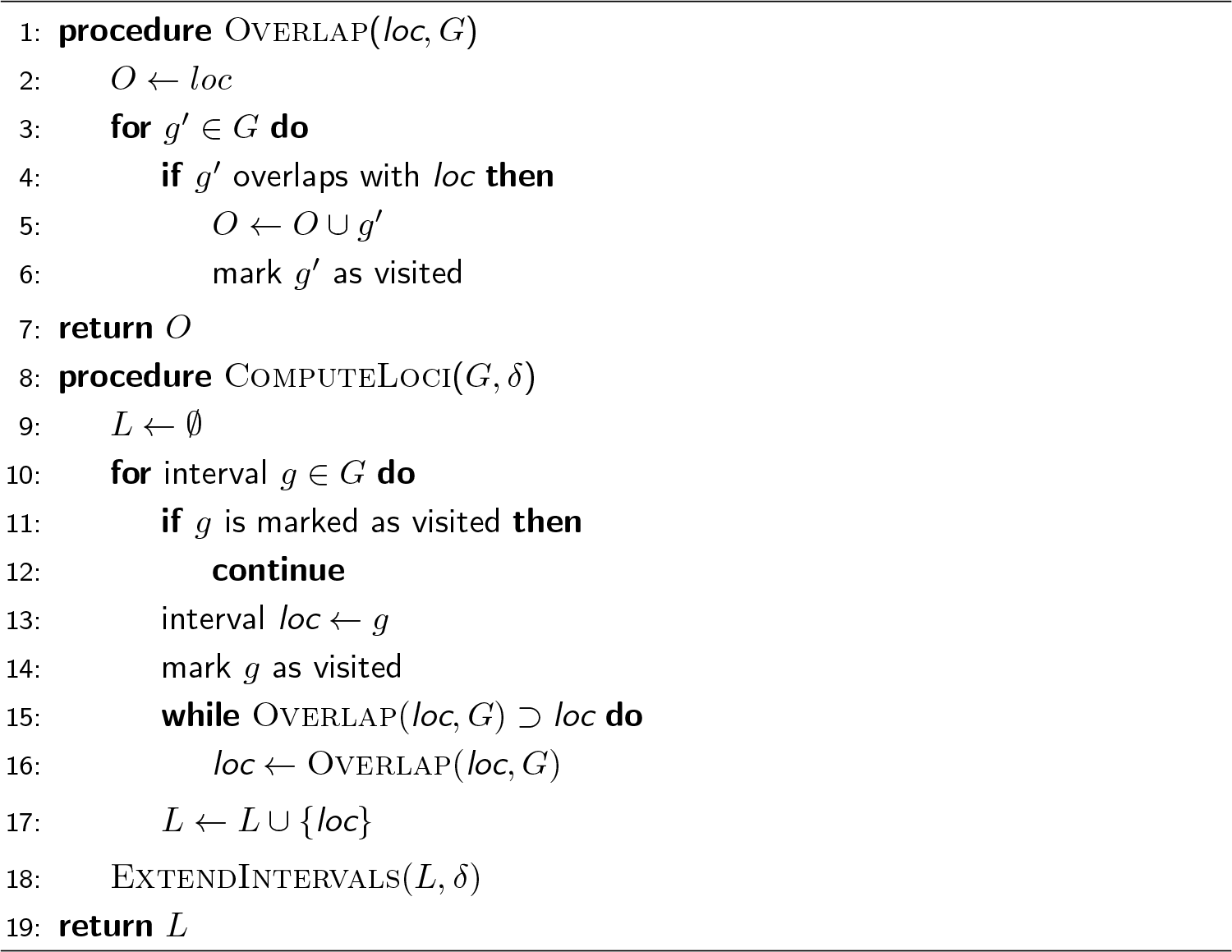

##### Algorithm 2 Extend giLocus boundaries, identify iiLoci

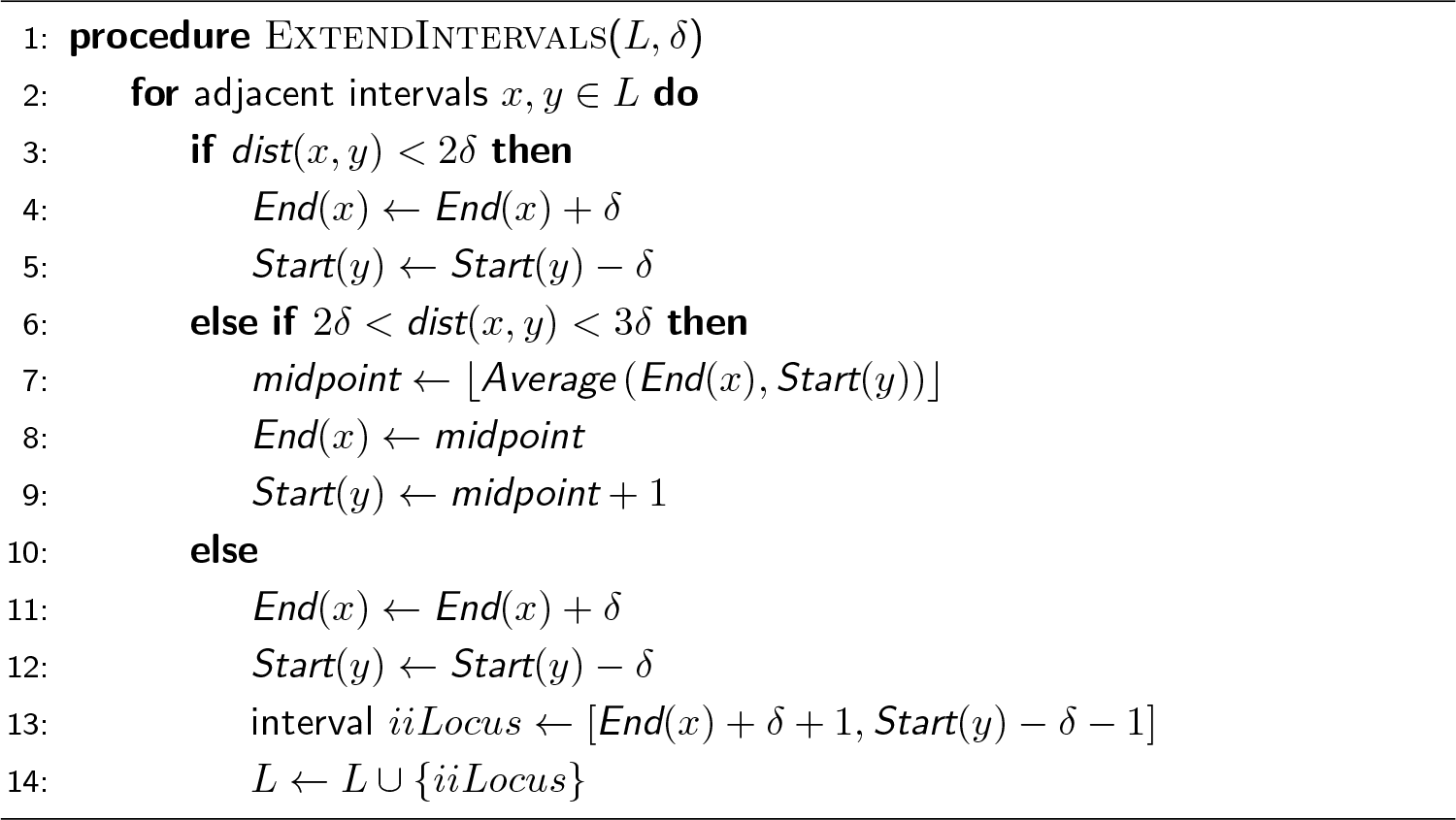

#### Post-processing to refine iLoci

The iLocus parsing procedure is designed with the canonical case of gene organization in mind: a single gene model flanked on both sides by hundreds or thousands of nucleotides of intergenic space. All eukaryotic genomes have exceptions to this case, some to a greater extent than others. The basic parsing procedure can handle some exceptions, such as genes separated by very little intergenic space, but there are additional exceptions that occur frequently enough to merit additional post-processing and refinement.

The basic procedure places two gene models in the same iLocus if their gene bodies have any overlap. While this is intended to capture gene models that may be conflicting or misannotated and in need of additional attention to resolve coordinates, an unintended consequence is the occasional grouping of genes with a trivial amount of incidental overlap. For example, if two genes—each a few kilobases in length—happen to have 10-20 nucleotides of overlap in their UTRs, they should be separated and handled as distinct loci. In post-processing, we enable splitting of such trivially overlapping iLoci by introducing two additional parameters: *ω*, the number of nucleotides that two gene models must overlap to remain in the same iLocus, and *κ* indicating whether that overlap is calculated using entire gene bodies (*κ* = 0) or just the coding sequences (*κ* = 1).

The initial procedure also groups ncRNA genes and protein-coding genes together if they overlap. In post-processing, ncRNA genes and protein-coding genes are treated separately and will not be grouped in the same iLocus regardless of overlap, although overlapping ncRNA genes are grouped in the same niLocus.

An additional exception occurs when a gene resides completely within a single intron of another gene. These genes are placed in the same iLocus during the initial parsing procedure, but are separated into distinct iLoci during post-processing.

### Implementation

In keeping with the conventions implemented by the GenomeTools library [15], most of the core functionality of the AEGeAn Toolkit [10] is implemented by means of *node streams* for sequential processing of genome features that are represented as *feature graphs*. In brief, genome features such as genes, exons, UTRs, and coding sequences are represented as nodes in a directed acyclic graph, and parent/child relationships between features, denoted by *ID* and *Parent* attributes in GFF3, are represented as edges in the graph. Each connected component (CC) in the graph, typically corresponding to a gene and its subfeatures, is then processed sequentially by one or more node streams, each designed for a specific annotation processing task. One advantage of this approach is that it leverages streaming algorithms with a low memory footprint, as at most only a small number of CCs need be loaded into memory at any given moment.

The *AgnLocusStream* module in the AEGeAn Toolkit implements a node stream for computing iLocus boundaries. This node stream expects as input gene annotations (CCs with a gene feature as the root node) sorted by genomic position, but it is designed to work with arbitrary feature types. Initially, the node stream will collect a single gene feature from the input and store it in a buffer. Any subsequent gene features that overlap with genes in the current buffer (that is, the leftmost position of candidate gene is less than or equal to the rightmost position of any gene in the buffer) are accumulated into the buffer. This continues until the node stream encounters a gene that does not overlap with the buffer, initiating two operations: first, the node stream emits a giLocus feature spanning all genes in the buffer; second, the node stream resets the buffer and begins accumulating the next gene or set of genes. A reference to the previously emitted giLocus is also maintained, enabling the refinement of boundaries between giLoci and, when appropriate, the designation of iiLoci, as described in **Algorithm 2**.

The AEGeAn Toolkit’s *AgnLocusRefineStream* module implements a node stream for post-processing the initial iLocus designations, as described in the previous section. Any genes belonging to the same giLocus that do not overlap by at least *ω* nucleotides in their gene bodies (or coding sequences if *κ* = 1), as well as genes contained completely within the intron of another gene, are split into distinct overlapping giLoci.

More generally, the AEGeAn Toolkit includes a variety of components. Node streams and other core components are implemented in the C language and organized into reusable modules. All core modules are compiled into a single shared object file to facilitate integration with other software by dynamic linking. Finally, a variety of executable programs for annotation processing and analysis composed from these core modules are also provided. In particular, the *LocusPocus* program provides the primary user interface to the *AgnLocusStream* and *AgnLocusRefineStream* modules. A detailed description of command-line usage and program inputs and outputs is provided in the AEGeAn Toolkit’s source code distribution.

### Genome content statistics

As discussed in the **Background** section, derivation of genome characteristics for comparison across species requires selection of reliable subsets of data for analysis. The precise selection criteria used will depend on the questions being asked, but commonly involve a small set of descriptive statistics (see e.g. [18]) that can easily be computed from the iLocus sequence and/or associated annotation. These include the length and nucleotide composition of the iLocus itself, as well as the count, length, and composition of corresponding features such as genes, RNAs, exons, introns, and coding sequences. Statistics are computed by invoking the *stats* task of the AEGeAn *fidibus* script (see **Additional file 2: wfscripts/run-fidibus-stats.ipynb**) and are stored in tab-separated plain text (.tsv) files to facilitate import into popular statistics packages.

Additional characteristics for comparison and filtering may not always be directly accessible from the iLocus sequence or annotation but derive from computation using external data sources. Such values can then be attached to an iLocus annotation using key-value pairs in GFF3’s **attribute** column. For example, gene model quality can be measured with statistics such as Maker’s *annotation edit distance* [19] or the GAEVAL *integrity score* [20], and homology status can be determined via reciprocal BLAST searches or clustering of iLocus protein products.

Descriptive statistics are reported only for a single annotated transcript at each iLocus to ensure that aggregate statistics are not biased by redundancy in the data resulting from genes with many annotated isoforms, for example. The reported transcript is selected according to the amino acid length of its translation product: the transcript with the longest product is reported. In cases where multiple transcripts have translation products of identical length, the transcript with the lexicographically smallest **ID** attribute is reported, ensuring reproducible and deterministic reporting.

Cumulative lengths of different iLocus types are calculated after proper accounting of any iLocus overlaps to ensure each nucleotide in the genome is counted only once (see **Additional file 2: wfscripts/make-Tables1-3.sh**). When reported as a fraction of the entire genome, the genomic space occupied for different iLocus categories is calculated as a fraction of *effective genome size*, defined as the total number of nucleotides in the genome that do not reside within fiLoci. This will mitigate potentially confounding inflation of genome size by many short unannotated sequences or sequence fragments.

### Genome organization statistics

Beyond genome content, the iLocus framework also allows systematic study of different aspects of genome organization. Here we focus on gene orientation and spacing: are there species-specific patterns of gene arrangements, and how do natural genomes differ in these respects from statistical expectation (e.g. [21])? Because of the flexible design of the code base described in the **Implementation** section, these questions can easily be generalized and extended, for example with respect to selection of subtypes of genic loci.

To study gene orientation, the *LocusPocus* program reports for each iiLocus the transcriptional orientation of the flanking giLoci as FF, RR, RF, or FR, corresponding to forward, reverse, outward, and inward orientations, respectively. For example, FF indicates that both flanking genes are transcribed on the top strand relative to the given assembly and annotation. In the case that an iiLocus is flanked by one or more ciLoci, the orientation of the gene models directly flanking the intergenic space are reported. Differences in occurrence numbers and lengths of outward and inward iiLoci are determined for possible interpretation in terms of promoter architecture: outward orientation for a short iiLocus might correspond to a bidirectional promoter. One could also identify the longest stretches of genes all on the forward strand, all on the reverse strand, or periodically alternating between strands to probe the extent of co-linear transcription.

Long iiLoci are flagged as regions for annotation review. More generally, for each giLocus, the lengths of the flanking iiLoci are reported. In cases where a giLocus abuts or overlaps with another giLocus, the corresponding iiLocus length is set to zero, and the number of overlapping nucleotides is recorded. The software tracks these cases as *zero-length iiLoci* (*ziLoci*). The iiLocus lengths are used in two different ways to reveal gene spacing characteristics. First, the distribution of aggregate lengths of *n* adjacent iiLoci shows the mode of typical gene spacings as well as outliers. Secondly, overlapping or abutting giLoci are collapsed into *merged iLoci* (*miLoci*) during post-processing and represent gene clusters; the resulting ziLoci are reflected in statistics that measure the characteristics of *N* adjacent iiLoci or of all iiLoci in aggregate.

To evaluate observed gene spacing patterns with statistical expectation, we implemented a procedure to generate randomized gene arrangements relative to a given input genome annotation. First, iLoci are computed with *δ* = 0 to identify the precise boundaries of annotated genic regions. Next, giLoci are removed from the sequence and the remaining iiLoci are concatenated. Then, new positions are randomly selected from a uniform distribution for re-inserting the giLoci in shuffled order into the sequence. As each giLocus is re-inserted, the genomic sequence is expanded, and all downstream re-insertion site positions are adjusted accordingly. Re-running the iLocus parsing procedure and computing neighbor statistics on these random arrangements provides a baseline for comparison, revealing how genome annotations as observed differ (at the genome scale) from what could be expected from a completely random arrangement of genes.

### Comparing assembly/annotation pairs: iLocus stability

Given two assembly/annotation versions *A* and *B* for the same genome, the question arises how the *A* iLoci map onto the iLoci set calculated for *B*. Let us assume that *B* is a later, improved version of *A*. Two cases can be distinguished. In the first case, the genome assembly is the same for *A* and *B*, but the annotation has changed, for example by inclusion of newer experimental data that led to annotation of non-coding genes, novel splice forms, or rejection of previous hypothetical gene structure models. In the second case, both the genome assembly and the annotation have changed, the former presumably due to additional genomic sequencing data that led to a less fragmented assembly. The mapping of iLoci may include several possibilities: 1) an *A* iLocus maps essentially unchanged to a *B* iLocus (although its genomic sequence identifier and coordinates may be different in a new assembly; an old iLocus may not map at all to the *B* set; 3) set *B* may include novel iLoci; and 4), there may be partial mapping of iLoci, for example when a novel non-coding gene annotation breaks up genomic space that had previously been annotated as intergenic space.

Mapping of the iLoci may involve sequence alignments spanning considerable gaps, as would be the case when a newer assembly provides gap-filling compared to the older assembly. Thus, we chose the LASTZ pairwise aligner [22] and evaluate results based on the overall quality and length of the maximal chain of high-scoring segment pairs. Specifically, query iLoci sequences were matched against target iLoci sequence sets with LASTZ parameters *–ambiguous=iupac –filter=identity:95 –chain* (**Additional file 2: comparisons/run*.sh**). The output of LASTZ was processed as follows to provide a classification of a query locus *qlocus* of length *qlength* based on any chained matches (chain length *clength*) against a subject locus *slocus* of length *slength*:

- *qlocus* is **without hits**
- *qlocus* has no qualifying hits and is designated as **unmapped**
- *qlocus* matches *slocus* such that *clength/qlength ≥* 0.9 and *clength/slength ≥* 0.9 and is designated as **conserved**
- *qlocus* matches *slocus* such that *clength/qlength ≥* 0.9 and *clength/slength <* 0.9 and is designated as **contained**
- *qlocus* matches *slocus* such that *clength/qlength <* 0.9 and *clength/slength ≥* 0.9 and is designated as **anchored**
- *qlocus* is designated **redefined** if there are subject loci with respect to which it is **contained** and others with respect to which it is **anchored**

Cases in which a query ilocus is conserved with respect to multiple subject iloci may occur when the assemblies contain duplicated genes and are noted in the LASTZ parsing script output.

### Comparing genome content and organization between related genomes: homologous iLoci

Given a set of annotated genome assemblies for a clade of related species, we compute *homologous iLoci (hiLoci)* via a protein clustering procedure. For each species, a representative protein sequence is selected for each piLocus (as described in the **Genome content statistics** section). The distinct protein complements from all species are then combined, and the aggregate collection of protein sequences is clustered using cd-hit [23].

In brief, cd-hit processes proteins iteratively from longest to shortest. The first protein is assigned to a cluster by itself and is designated the *representative sequence* of the cluster. Each subsequent protein is compared to all previous clusters: If the alignment of the protein to a cluster’s representative sequence satisfies the specified sequence identity, length similarity, and alignment coverage criteria, it is added to that cluster, and the program advances to the next protein; if a protein cannot be added to any cluster by user-specified clustering criteria, it is placed in a new cluster by itself and designated the representative sequence of that cluster.

Following the clustering procedure, a data structure designated as *homologous iLocus* (*hiLocus*) is created for each protein cluster, and the piLoci corresponding to the proteins in that cluster are assigned to that hiLocus. The hiLocus thus provides a link between piLoci from related species and a relative measure of how well conserved the corresponding protein is within the given clade.

This protein clustering procedure is invoked using the *cluster* task of the AEGeAn *fidibus* script. The default parameters are as follows: sequence identity *≥* 50%; length difference *≤* 50%; alignment coverage for longer sequence *≥* 60%; alignment coverage for shorter sequence *≥* 60%. On the command line these parameters are specified as -c 0.50 -s 0.50 -aL 0.60 -aS 0.60. The default values can be overridden, and additional criteria can be set by the user.

### Data sets analyzed

We retrieved RefSeq genome assemblies and corresponding annotations for ten model organisms (as listed in Table 1) to illustrate the utility of iLoci for providing a descriptive overview of genome composition and organization. Species were selected to provide a broad sampling of eukaryotic diversity, with a preference for robust model organisms with mature chromosome-level genome assemblies and extensive community-supported annotation. For each species, we computed iLoci and associated feature statistics, including length, nucleotide composition, exon count, and *effective length*, using standard fidibus build tasks as described before.

**Table 1:** iLocus content of genomes from ten model organisms and four additional species.

| Species | Mb<sup>1</sup> | #Seq<sup>2</sup> | fiLoci | iiLoci | niLoci | siLoci | ciLoci |
| --- | --- | --- | --- | --- | --- | --- | --- |
| *Saccharomyces cerevisiae* | 12.1 | 16 | 11 | 274 | 393 | 5,777 | 101 |
| *Caenorhabditis elegans* | 100.3 | 6 | 3 | 6,152 | 19,843 | 19,373 | 359 |
| *Chlamydomonas reinhardtii* | 120.2 | 1,556 | 1,487 | 6,245 | 0 | 14,253 | 42 |
| *Medicago truncatula* | 429.6 | 40 | 64 | 28,175 | 5,929 | 31,350 | 142 |
| *Anopheles gambiae* | 265.0 | 8,089 | 8,037 | 7,726 | 644 | 11,986 | 318 |
| *Drosophila melanogaster* | 143.7 | 1,869 | 1,874 | 3,452 | 3,356 | 12,626 | 650 |
| *Xenopus tropicalis* | 1437.5 | 7,727 | 8,004 | 18,580 | 5,181 | 21,454 | 403 |
| *Danio rerio* | 1371.7 | 1,060 | 1,295 | 23,979 | 15,208 | 25,392 | 481 |
| *Mus musculus* | 2725.5 | 21 | 42 | 23,771 | 13,342 | 21,117 | 616 |
| *Homo sapiens* | 3088.3 | 24 | 48 | 22,242 | 16,584 | 18,947 | 927 |
| *Volvox carteri* | 137.7 | 1,251 | 1,198 | 7,790 | 0 | 14,346 | 44 |
| *Polistes dominula* | 208.0 | 1,483 | 1,665 | 3,969 | 1,049 | 9,715 | 282 |
| *Daphnia pulex* | 197.3 | 5,191 | 4,759 | 13,052 | 0 | 30,454 | 160 |
| *Manacus vitellinus* | 1072.3 | 3,619 | 3,760 | 11,319 | 1,999 | 13,096 | 193 |

<sup>1</sup>Total number of nucleotides in the genome assembly.<sup>2</sup>Total number of assembled (pseudo-)chromosomes plus any unplaced genomic scaffolds.

Using iLocus summaries of these ten model organisms as a baseline for comparison, we characterized the genome content and organization of four additional species of interest that serve as important experimental models for evolutionary and ecological studies: the microcrustacean *Daphnia pulex*, the primitively eusocial paper wasp *Polistes dominula*, the green alga *Volvox carteri*, and the suboscine passerine bird *Manacus vitellinus*. These four genomes were processed using the same procedure as the ten model organisms. Precise configurations and commands run for all analyses are available in **Additional file 1** and at https://github.com/BrendelGroup/iLoci_SLB21.

Finally, we retrieved and processed, in the same manner as above, large collections of genomes from NCBI RefSeq branches and computed branch averages of all statistics of interests. We report on these statistics as another baseline for genome evaluation in taxonomic evolutionary context.

#### Classifying hiLoci from a clade of 9 chlorophyte species

To investigate the extent of gene conservation in the green algae (phylum: Chlorophyta), we collected and processed data for nine chlorophyte species (*Auxenochlorella protothecoides, Chlamydomonas reinhardtii, Chlorella variabilis, Coccomyxa subellipsoidea, Micromonas commoda, Micromonas pusilla, Ostreococcus lucimarinus, Ostreococcus tauri*, and *Volvox carteri*), as well as four land plants (*Arabidopsis thaliana, Brachypodium distachyon, Medicago truncatula*, and *Oryza sativa*) as an outgroup. Retrieval of annotations and sequences and calculation of hiLoci was invoked using standard procedures as described in previous **Methods** sections (and **Additional file 2: README explore-Chlorophyta.md**). Following the protein clustering procedure, each hiLocus was assigned a preliminary classification: *highly conserved* if it had a representative from each of the nine chlorophyte genomes; *conserved* if it had a representative from at least four chlorophyte genomes; *matched* if it had a representative from at least two genomes (including the outgroups); and *unmatched* if it had a representative from only a single genome. hiLoci initially classified as *unmatched* were subjected to additional screening to distinguish conserved proteins lacking a nearly-full-length match (due to incomplete or incorrect annotation, or true evolutionary divergence) from orphan proteins without any reliable match. hiLoci with a BLASTP match against another species (`-evalue 1e-10`) were reclassified as *matched*, while those lacking a match were reclassified as *orphan*.

## Results

### iLoci provide an informative decomposition of genome content

We computed iLoci for ten model organisms representing a wide range of eukaryotic diversity and provide a summary of each genome and its iLocus complement in **Table 1** (for workflow commands, see **Additional file 2: README refr-genome-summary.md**). The genome assembly sizes in this sampling of eukaryotes span two orders of magnitude, ranging from 12.1 Mb in *Saccharomyces cerevisiae* to over 3 Gb in *Homo sapiens*. Several genomes are represented exclusively by chromosome sequences, some exclusively by unplaced genomic scaffolds, and some by a combination of both. The number of fiLoci, with a strict upper bound of twice the number of assembled sequences, is informative primarily with respect to assembly status. For most of these genomes, the observed number of fiLoci is close to half of the upper limit. There are two reasons for why the observed number of fiLoci can be lower than the upper limit: (1) the presence of gene annotations near the end of a genomic sequence (within 2 *× δ*, in which case no fiLocus is recorded); and (2) the inclusion of unannotated (short) scaffolds in the genome sequence set (which results in one fiLocus per unannotated scaffold spanning the entire sequence). Here, for example, the numbers for *S. cerevisiae* are consistent with a compact genome, the numbers for *C. reinhardtii* are consistent with a fragmented genome assembly including many unannotated scaffolds, and the numbers for mouse and human are consistent with complete genomes.

iiLoci correspond to intergenic DNA and are reflective of genome organization. There can be at most *n − m* iiLoci in a genome with *n* genes and *m* annotated sequences, but closely-spaced genes will reduce the number of observed iiLoci, as will the presence of unannotated scaffolds.

The abundance of piLoci in each genome (representing distinct protein-coding regions) spans just a single order of magnitude, from 5,878 piLoci in *Saccharomyces cerevisiae* to 31,492 in *Medicago truncatula* (**Table 2**). The total space occupied by piLoci, however, spans two orders of magnitude, similar to genome size. This is explained by a distinct contrast in siLocus length between vertebrates and the other species (**Additional file 1: Figure S1**; (for workflow commands, see **Additional file 2: notebooks/make-SF1.ipynb**)), the compound result of increases in both intron abundance and length (**Additional file 2: notebooks/make-SF5c-SF8.ipynb**). We note that while the protein-coding gene portion of the human genome is commonly reported as 2-4%, this refers only to protein-coding exons. The inclusion of introns and UTRs places the protein-coding gene fraction of the genome at approximately 40% for both human and mouse. ciLoci are present in dozens to hundreds in most genomes, accounting for only a small proportion of genes.

**Table 2:** Summary of piLoci from genomes of ten model organisms and four additional species.

| Species | piLoci | Occupancy<sup>1</sup> | Single exon piLoci |
| --- | --- | --- | --- |
| *Saccharomyces cerevisiae* | 5,878 | 11.4 Mb (94.6%) | 5,613 (95.5%) |
| *Caenorhabditis elegans* | 19,732 | 75.7 Mb (75.5%) | 685 ( 3.5%) |
| *Chlamydomonas reinhardtii* | 14,295 | 74.1 Mb (68.2%) | 1,127 ( 7.9%) |
| *Medicago truncatula* | 31,492 | 158.6 Mb (37.0%) | 5,701 (18.1%) |
| *Anopheles gambiae* | 12,304 | 83.8 Mb (35.9%) | 1,154 ( 9.4%) |
| *Drosophila melanogaster* | 13,276 | 95.4 Mb (70.0%) | 2,100 (15.8%) |
| *Xenopus tropicalis* | 21,857 | 687.2 Mb (50.4%) | 1,344 ( 6.2%) |
| *Danio rerio* | 25,873 | 828.8 Mb (61.2%) | 1,095 ( 4.3%) |
| *Mus musculus* | 21,733 | 1,034.4 Mb (38.9%) | 2,370 (11.0%) |
| *Homo sapiens* | 19,874 | 1,240.1 Mb (41.2%) | 1,270 ( 6.5%) |
| *Volvox carteri* | 14,390 | 89.2 Mb (69.1%) | 1,086 ( 7.5%) |
| *Polistes dominula* | 9,997 | 137.7 Mb (72.4%) | 405 ( 4.1%) |
| *Daphnia pulex* | 30,614 | 89.2 Mb (54.3%) | 5,053 (16.5%) |
| *Manacus vitellinus* | 13,289 | 461.4 Mb (44.6%) | 529 ( 4.0%) |

<sup>1</sup>Total number of nucleotides occupied by piLoci and the corresponding fraction of effective genome size.

### iLoci reflect patterns of genome organization

#### Gene clustering is abundant in eukaryotic genomes

There are well-described examples of gene clusters in eukaryotic genomes, such as those associated with *Hox* genes [24]. *Hox* clusters are composed of functionally related developmental genes with a conserved colinear arrangement, a common direction of transcription, occurring in close proximity in the genome. More generally, gene clusters described in the literature need not be comprised only of genes that are directly adjacent, but are loosely defined as sets of genes of a common function situated much closer to each other than would be expected by chance [25]. However, the spatial distribution of genes in general, the extent to which genes are tightly packed throughout the entire genome, and the characteristics of these gene-dense regions have not been extensively studied in eukaryotes. *Merged iLoci (miLoci)* provide a well-defined unit of analysis for investigating the spatial distribution of genes genome-wide. Using miLoci, we surveyed genome organization in the selected ten model organisms.

Genes cluster together frequently in eukaryotic genomes. The most frequent groupings involve a small number (2-4) of genes (see **Table 3**), but all genomes include larger clusters involving dozens or even hundreds of tightly packed genes. The budding yeast *Saccharomyces cerevisiae* is an extreme example, populated almost entirely by just 294 miLoci encompassing all but 176 genes in the entire genome. *Caenorhabditis elegans* and *Drosophila melanogaster* also bear signatures of a higher overall level of genome compactness, with larger numbers (and overall proportion) of genes merged into miLoci and a larger proportion of genomic sequence occupied by miLoci.

**Table 3:** Summary of miLoci from genomes of ten model organisms and four additional species.

| Species | miLoci | Occupancy<sup>1</sup> | Median # genes<sup>2</sup> | Singletons<sup>3</sup> |
| --- | --- | --- | --- | --- |
| *Saccharomyces cerevisiae* | 294 | 11.1 Mb (92.0%) | 12 | 176 ( 2.8%) |
| *Caenorhabditis elegans* | 4,496 | 74.1 Mb (73.9%) | 5 | 2,425 ( 6.1%) |
| *Chlamydomonas reinhardtii* | 3,029 | 54.5 Mb (50.2%) | 3 | 3,796 (26.6%) |
| *Medicago truncatula* | 5,715 | 61.1 Mb (14.3%) | 2 | 22,657 (60.5%) |
| *Anopheles gambiae* | 2,036 | 26.2 Mb (11.2%) | 2 | 6,521 (50.4%) |
| *Drosophila melanogaster* | 2,155 | 75.2 Mb (55.2%) | 4 | 2,626 (15.8%) |
| *Xenopus tropicalis* | 3,224 | 174.0 Mb (12.8%) | 2 | 17,698 (65.5%) |
| *Danio rerio* | 5,843 | 301.6 Mb (22.3%) | 2 | 19,348 (47.1%) |
| *Mus musculus* | 6,039 | 661.9 Mb (24.9%) | 2 | 18,558 (52.9%) |
| *Homo sapiens* | 6,790 | 932.8 Mb (31.0%) | 2 | 16,668 (45.7%) |
| *Volvox carteri* | 3,229 | 57.0 Mb (44.1%) | 2 | 5,256 (36.5%) |
| *Polistes dominula* | 2,085 | 74.1 Mb (38.9%) | 3 | 2,870 (26.0%) |
| *Daphnia pulex* | 6,252 | 58.4 Mb (35.6%) | 3 | 9,934 (32.4%) |
| *Manacus vitellinus* | 2,156 | 118.5 Mb (11.5%) | 2 | 10,061 (65.8%) |

<sup>1</sup>Total number of nucleotides occupied by miLoci and, in parentheses, the corresponding fraction of effective genome size.
<sup>2</sup>Median gene count per miLocus.
<sup>3</sup>Total number of giLoci not contained in miLoci and, in parentheses, corresponding fraction of all giLoci.

In general, clustered genes do not differ substantially in length or nucleotide composition from spaced out genes. However, especially among large miLoci, clustered genes are often functionally related. The longest miLoci in the human genome include a cluster of 22 snoRNA genes on chromosome 14, a cluster of 19 genes from AP2A1 to NUP62CL on chromosome 19, and a cluster of keratin associated proteins on chromosome 21, while in mouse the longest miLocus is comprised of 76 microRNA genes, on chromosome 2. In the non-mammal vertebrates, the longest miLoci consist exclusively of long stretches of hundreds tRNA gene annotations. As tRNA-derived SINE transposons are known to be abundant in at least one of these species [26], and no annotations for such transposons appear to be included in the RefSeq annotation, it is likely these miLoci capture large clusters of misannotated repetitive elements. The latest annotation of *Medicago truncatula* includes several rRNA gene clusters, identified in the miLoci list by our default parameter *δ* = 500. The distribution of miLoci along the chromosome is mostly uniform for compact genomes such as *Drosophila* and *C. elegans*. For less compact genomes, we observe variation in the uniformity of miLocus distribution. For example, in *Medicago*, miLoci appear to be more frequent at the chromosome ends, while in vetebrate species a depletion of miLoci in pericentromeric regions is most obvious (**Additional file 2: notebooks/explore-miLoci.ipynb**).

The spacing of genes over longer ranges is revealed by distributions of aggregate lengths of *r* adjacent iiLoci (*r-scans* [27]). Long-range spacing of genes varies considerably in eukaryotes, with some species exhibiting homogeneous gene spacing over relatively short spans (spans of 5-10 genes in *Caenorhabditis elegans* and *Medicago truncatula*), and others showing heterogeneous spacing even over long spans (spans of more than 30 genes in *Mus musculus*; see **Additional file 1: Figure S2**; for workflow commands, see **Additional file 2: notebooks/make-SF2.ipynb**).

#### Gene orientation

The iLocus framework provides a convenient approach to analyzing the strand locations of genes. We categorize the iiLoci based on the length and orientation of the flanking giLoci, as described in subsection **Genome organization statistics** in **Methods**. The stacked barplots showing the distribution of iiLocus length, grouped by orientation, are given for the ten model organisms in **Additional file 1: Figure S3** (generated by **Additional file 2: notebooks/make-SF3.ipynb**) and for the randomized gene positioning control in **Figure S4** (generated by **Additional file 2: notebooks/make-SF4.ipynb**). Note that for this study, iLoci were determined with *δ* = 0 to allow investigation of short intergenic regions (for workflow commands, see **Additional file 2: wfscripts/run-explore-gene-orientation.sh**).

Comparing the two sets of figures, it is clear that the iiLocus orientation types do not occur in random proportions in the natural genomes. However, the patterns of deviation depending on iiLocus length are different between species. *Anopheles* and *Drosophila* show the most even pattern across all length bins. Mouse and human show an intriguing preponderance of the outward (RF) orientation type for short iiLoci but relative avoidance of the type for longer iiLoci. Zebrafish (*Danio rerio*) seems to favor the colinear types FF and RR until the longest iiLocus length bins. *C. elegans* shows a preference for FF in the same length ranges. Lastly, *M. truncatula* has high numbers of inward (FR) orientation types for iiLocus lengths up to around 1 kb. Detailed interpretations of these differences would involve exploration of gene types and chromosomal location, but for here we simply emphasize the readily availability of these genome organization data within the iLocus framework.

#### Compactness of eukaryotic genomes varies widely

We further explore the notion of compactness of a genome by two complementary measures calculated on the constituent chromosome or long scaffold sequences: *ϕ*, defined as the fraction of giLoci in the sequence merged into miLoci; and *σ*, defined as the proportion of the sequence occupied by miLoci. Distinct quadrants in the plot reflect characteristic overall genome organization. Low values of *ϕ* associated with low values of *σ* (lower left) correspond to genes as “islands” in an “ocean” of intergenic (presumably repetitive) DNA. High values of *ϕ* associated with low values of *σ* (lower right) correspond to “archipelagos” of genes. And high values of *ϕ* associated with high values of *σ* (upper right) correspond to “compact” (or “continental”) genome organization.

Let the average iiLocus length be *ρ*-times the average giLocus length (*g*), and let *m* and *n* be the number of giLoci and iiLoci, respectively. Then *σ* = *ϕmg/*(*mg* + *nρg*), and if *ϕ* is small, then *n ≈* (1 *− ϕ*)*m*, and the following approximation holds:

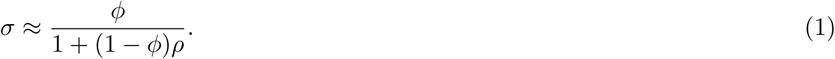

When *ϕ* is close to one, then also *σ ≈ ϕ*, unless the genome had very distinct densely packed multi-genic regions separate from substantial non-genic regions. Thus, major deviations from the expected curve are revealing of extreme genome organization, as discussed above. **Figure 3A** gives the curves for *ρ* equal to 0.1, 1, 2, 4, and 8 (produced by **Additional file 2: notebooks/make-F3a-SF6-SF7.ipynb**; for workflow commands, see **Additional file 2: README refr-genome-compactness.md** and **Additional file 2: wfscripts/run-explore-compactness-refr.ipynb**).

**Figure 3:**
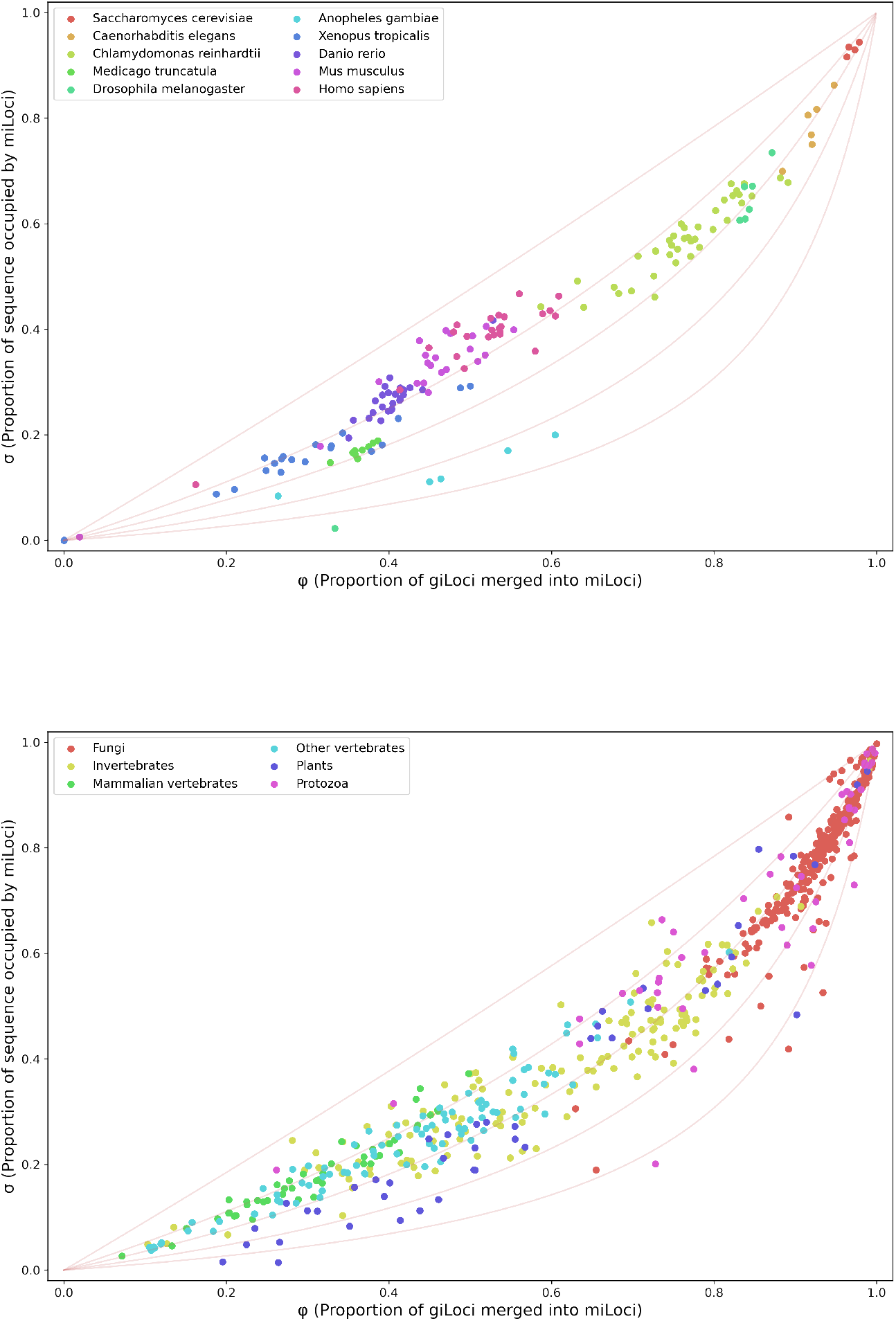
Genome compactness. **A. Ten reference genomes.** The curves correspond from top to bottom to the theoretical *ϕ, σ* functions (Equation 1) for *ρ* equal to 0.1, 1, 2, 4, and 8, respectively. Each data point corresponds to a sequence of length at least 1Mb. Short giLoci (lower 5%-tile) and long iiLoci (top 5%-tile) were removed for each genome prior to calculation. **B. Compactness as a persistent genome characteristic**. Centroids of (*ϕ, σ*) values calculated as in A. for different genomes from the indicated taxonomic groups.

Empirical (*ϕ, σ*) values calculated for continuous genome sequences of at least 1 Mb for the 10 model species reveal a wide range of genome compactness across eukaryotes, yet show remarkable consistency within species (**Figure 3A**) and even within clades and branches (as confirmed by sampling of additional species; see **Additional file 1: Figures S5a-S5d** and **Figure 3B**; for workflow commands, see **Additional file 2: notebooks/make-F3b.ipynb** and **Additional file 2: wfscripts/run-explore-compactness-othg.sh** and **taxa/README.md**). Genome compactness scales roughly with genome size, at least across major clade divisions and levels of organismal complexity. Within Chlorophyta, compactness scales almost perfectly with genome size, although this trend is not maintained in clades characterized by larger genome sizes. Consistent with previous described observations, sequences from *Saccharomyces cerevisiae* are the most compact of all 10 model organisms analyzed. Alternatively, very few sequences show extremely low levels of overall compactness: only six sequences of the ten model organisms have *ϕ <* 0.2 and *σ <* 0.2, two of which correspond to mammalian sex chromosomes, with the other four corresponding to unplaced scaffolds from *Xenopus tropicalis*. This trend continues with most other genomes and annotations from RefSeq, with only a few genome averages below both thresholds (**Figure 3B**). Likewise, very few sequences are dominated by an “archipelago”-type organization (high *ϕ* and low *σ*). Those with *ϕ >* 0.7 and *σ <* 0.3 are annotated almost exclusively with long stretches of dozens or hundreds of tRNA gene annotations in *Xenopus tropicalis* and *Danio rerio*.

Adjusting the value of the *δ* parameter used in the initial iLocus parsing procedure can have a moderate effect on (*ϕ, σ*) measures of genome compactness. As expected, reducing the length of *δ* extensions results in a decrease in reported genome compactness, while increased values of *δ* result in reports of higher genome compactness (**Additional file 1: Figure S6**). However, relative compactness between different genomes appears robust to changes in the *δ* parameter.

#### Gene clustering occurs more frequently than expected by chance

To investigate whether gene clustering occurs more frequently than expected by chance, we computed random arrangements of genes on each long (*≥* 1 Mb) chromosome or scaffold sequence and re-computed iLoci and associated summary statistics for comparison with the observed annotation.

Random positioning of genes results in decreased levels of gene clustering across all species as reflected by several measures: a decrease in the number of miLoci; a decrease in the space occupied by miLoci; a decrease in the number of genes per miLocus; and an increase in the number of singleton genes not associated with miLoci (**Additional file 1: Table S1**). Signatures of genome compactness are also influenced by random arrangement of genes, reflecting less compactness relative to the actual annotated positioning of genes. The (*ϕ, σ*) statistics calculated on long genomic sequences are consistently lower for random arrangements than actual arrangements for all model species (**Additional file 1: Figure S7**), with the exception of the extremely compact *Saccharomyces cerevisiae* genome.

### “LocusPocus Fidibus”: an incantation for any genome

Evaluating new genome assemblies and annotations is a common and critical challenge in contemporary biology, but is hampered by limited community bioinformatics support and the lack of precise standards for systematic comparisons of genome content and arrangement. Having explored the range of genome composition and organization in eukaryotic model organisms, we turn now to the question of newly sequenced genomes: How does the new genome fit into the broader universe of eukaryotic organisms, and more interestingly, how does the new genome compare to genomes from closely related species?

iLoci address these challenges both by offering a well-defined “common currency” for comparisons of genome content and organization and by providing associated software tools to facilitate analysis and re-analysis of old and new data alike. The *LocusPocus* and *Fidibus* programs are designed for painless adoption by researchers with minimal bioinformatics expertise and require only a small number of standard input files. In return, they produce a wealth of descriptive statistics not only on iLoci but also on their constituent genes, transcripts, and associated features.

With baseline expectations about eukaryotic genome content and organization established by iLocus analysis of large numbers of genomes from RefSeq, including ten model organism genomes, we now demonstrate how these tools can be applied to evaluate genomes of particular species of interest.

#### Volvox carteri

The green algae (phylum Chlorophyta) diverged from land plants an estimated 1 billion years ago [28] and encompass a diverse set of organisms ubiquitous in marine and soil environments. Chlorophytes exhibit substantial variation in physical stature, genome size, and cellular complexity, and include many important systems for study of the evolution of multicellularity and photosynthesis. The publication of the *Volvox carteri* genome [29] reported over 5,000 protein families conserved between *Volvox* (a multicellular alga) and *Chlamydomonos reinhardtii* (a unicellular relative), accounting for over a third of both species’ respective proteomes.

The genome content of *Volvox* is very similar to that of *Chlamydomonas* across a variety of iLocus measures. Characteristics of protein-coding regions in particular (summarized in **Tables 1-2**) show striking similarity: piLoci account for 89.2 Mb (69.1%) of the *Volvox* genome (compared to 74.1 Mb (68.2%) of the *Chlamydomonas* genome), and both genomes harbor a similar number of single-exon piLoci (1086 vs 1127, respectively) and very few ciLoci (44 and 42, respectively). With respect to genome organization, *Volvox* and *Chlamydomonas* contain comparable numbers of miLoci (3229 and 3029, respectively; **Table 3**) and exhibit a remarkably similar level of gene density. The (*ϕ, σ*) values measuring genome compactness of the two species fall within a nearly identical range, with *Volvox* shifted to slightly lower values (**Additional file 1: Figures S5a**, produced by **Additional file 2: notebooks/make-F4a-F4b-SF5a.ipynb**). These observations are consistent with the claims that, despite an estimated 50-200 million years of divergence and major differences in cellular complexity, the genomes of *Volvox* and *Chlamydomonas* are impressively similar [29].

With several representative chlorophyte genomes now available from RefSeq [30], we leveraged iLoci to characterize the extent of gene conservation in *Volvox* relative to the entire phylum. piLoci from all nine species were grouped together as *homologous iLoci (hiLoci)* based on a clustering of their protein products, and the relative conservation status of each hiLocus was determined (see **Methods**). **Figure 4** presents a breakdown of all nine genomes according to iLocus type and conservation status, showing both the number of iLoci in each category as well as the proportion of the genome occupied by iLoci from each category (figure produce by **Additional file 2: notebooks/make-F4a-F4b-SF5a.ipynb**). Counts and aggregate space occupied by intergenic regions and assembly fragments (iiLoci and fiLoci, respectively) reflect the diversity of genome size and gene density across Chlorophyta, ranging from 10-25 Mb genomes almost completely devoted to proteincoding genes (in *Micromonas* and *Ostreococcus*) to genomes well over 100 Mb in size with abundant intergenic space (in *Volvox* and *Chlamydomonas*).

**Figure 4:**
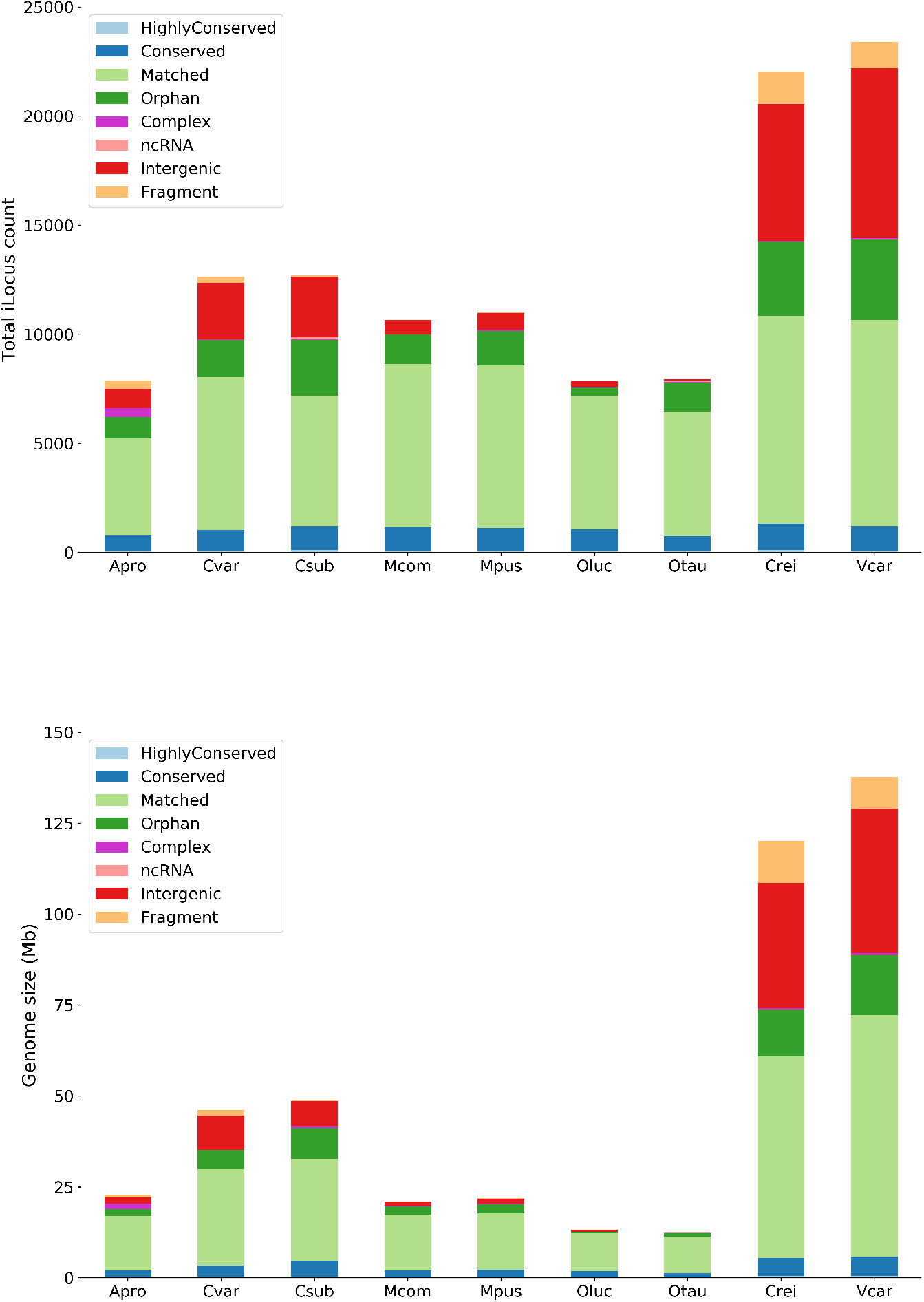
Breakdown of piLocus conservation status across chlorophyte species.

A small number of piLoci from each genome are designated as *orphans*, indicating no reliable protein match in any other species, while the majority are designated as *matched*, having at least one match in another species. The designations *conserved* and *highly conserved* were applied only to hiLoci whose protein products are well conserved throughout the phylum (*conserved*: conserved in at least 4 species; *highly conserved*: conserved in all 9 species) and differ in amino acid length by no more than a factor of 2 within a hiLocus. Given these stricter criteria, we observe on the order of 100 *highly conserved* piLoci and 1000 *conserved* piLoci in each species. A total of 3130 *Volvox* piLoci and 3261 *Chlamydomonas* piLoci were grouped into 2928 common hiLoci, 2803 of which contain a single ortholog from both species.

Highly conserved piLoci are associated with a variety of cellular components and processes, most prominently proteins related to ribosomes and kinase/phosphatase activity. The vast majority of orphan piLoci are annotated as “predicted” or “hypothetical proteins.” Among the handful with functional annotations, flagellar-associated proteins are prominent in *Chlamydomonas reinhardtii* orphans, while Jordan transposition proteins are prominent in *Volvox carteri* orphans.

#### Polistes dominula

The paper wasp *Polistes dominula* is an important model for the study of social behavior and evolution and was one of the first species of the family Vespidae to have its genome sequenced [31]. The *Polistes* genome is intermediate across many measures relative to the survey of ten reference genomes, in particular the two insect genomes (the fruit fly *Drosophila melanogaster* and the mosquito *Anopheles gambiae*). *Polistes* contains 3,969 iiLoci occupying 48.8 Mb (23.4%) of the genome, compared to 3,452 iiLoci occupying 35.5 Mb (24.7%) of the *Drosophila* genome and 7,726 iiLoci occupying 149.2 Mb (56.3%) of the *Anopheles* genome (**Table 1** and **Additional file 2: notebooks/make-SF5c-SF8.ipynb**). *Polistes* is distinct from the other insects, however, in that both siLoci and ciLoci are less abundant in its genome, and yet collectively they account for a larger proportion of the genome and a larger amount of absolute space (**Tables 1-2**). Similar results are observed when compared against invertebrates: raw counts of iLoci are comparable across each category, with a decreased number of piLoci, yet the relative occupancy of piLoci is greater (**Additional file 1: Table S2**).

In terms of gene organization, the *Polistes* genome harbors 2,085 miLoci, compared to 2,155 in *Drosophila*, 2,036 in *Anopheles* (**Table 3**), and a median of 2,298 miLoci in invertebrates (**Additional file 1: Table S3**). The (*ϕ, σ*) statistics computed on miLoci reveal an intermediate level of genome compactness ((**Additional file 1: Figures S5b**, produced by **Additional file 2: notebooks/make-SF5b.ipynb**). The increased dispersion of (*ϕ, σ*) values per sequence in *Polistes* is reduced in the longest genomic scaffolds, likely reflecting local fluctuations in genome organization that are evened out in the pseudo-chromosome level assemblies for *Drosophila* and *Anopheles*.

#### Daphnia pulex

The water flea *Daphnia pulex* is a species of ecological and evolutionary interest, and was the first crustacean genome to be sequenced [32]. Like *Polistes dominula*, characteristics of genome content and organization in *Daphnia pulex* are intermediate relative to the two arthropods surveyed. The most striking feature of the *Daphnia* genome is the large number of annotated genes and large fraction of single-exon piLoci (**Table 2**). *Daphnia* contains 30,614 piLoci, more than twice the number in *Drosophila, Anopheles*, and the median in all invertebrates (**Additional file 1: Table S2**) and second overall only to *Medicago*. However, the space occupied by these piLoci—89.2 Mb (54.3%) of the genome—is around average with respect to the species surveyed.

The amount of intergenic space in the *Daphnia pulex* genome is also moderate— iiLoci account for 75.1 Mb (38.0%) of the *Daphnia pulex* genome, compared to 35.5 Mb (24.7%) in *Drosophila* and 149.1 Mb (56.3%) in *Anopheles*. However, the abundance of piLoci punctuating the intergenic space results in an elevated number of (shorter) iiLoci (13,052 in contrast to 3,452 in *Drosophila* and 7,726 in *Anopheles*); see **Table 1**.

Claims regarding the relative compactness of the *Daphnia* genome, based primarily on average lengths of gene spans and introns, are not supported by our analysis [32]. iiLocus lengths are on average shorter in *Daphnia* compared to *Drosophila* and *Anopheles* (**Additional file 1: Figure S8a**, produced by **Additional file 2: notebooks/make-SF5c-SF8.ipynb**). We confirm that genes are on average shorter in *Daphnia* than in *Drosophila* (**Additional file 1: Figure S8b**), despite a larger number of exons per gene (**Additional file 1: Figure S8c**). However, this appears to be influenced more by reduced exon length rather than by reduced intron length, as originally claimed. Median exon length is substantially shorter in *Daphnia* (154 bp) versus 248 bp in *Anopheles* and 289 bp in *Drosophila*, respectively; see **Additional file 1: Figure S8d**). In contrast, median intron length of siLoci is almost indistinguishable between *Daphnia* and *Drosophila* (75 bp and 70 bp, respectively, and shorter than for *Anopheles* (91 bp); see **Additional file 1: Figure S8e**).

Further, although we observe consistently higher (*ϕ, σ*) values for *Daphnia* than for *Anopheles*, relative to *Drosophila* the values are consistently lower, reflective of a smaller fraction of tightly-packed genes and a smaller proportion of the genome sequence occupied by such gene clusters (**Additional file 1: Figure S5c**, produced by **Additional file 2: notebooks/make-SF5c-SF8.ipynb**). Thus, across multiple quantitative measures, *Daphnia pulex* is characterized by a moderate level of genome compactness relative to other arthropods and eukaryotes in general.

#### Manacus vitellinus

A widespread effort to collect and sequence avian genomes was undertaken in 2014, spanning most orders of bird species, including 38 new genome assemblies [33]. As a representative species, we chose the golden-collared manakin (*Manacus vitellinus*) with the latest NCBI assembly/annotation available from July, 2019 ([34]).

The current genome assembly is still highly fragmented given the large number of sequences and fiLoci for *M. vitellinus* (**Table 1**). Relatively few piLoci (13,289) occupy 44.6% of the genome space (**Table 2**), a value closer to the mammalian average than the average of other vertebrates (**Additional file 1: Table S2**). Notable is the small number of single exon piLoci (**Additional file 1: Table S2**). The miLocus count (2,156) and genome occupancy (11.5%) is considerably lower compared to human and mouse and also low relative to vertebrate averages (**Additional file 1: Table S3**), while the siLocus proportion of all giLoci is large at 65.8% (**Table 3**). Correspondingly, the (*ϕ, σ*) statistics computed on miLoci confirm a low level of genome compactness (see **Additional file 1: Figure S5d**, produced by **Additional file 2: notebooks/make-SF5d.ipynb**).

More complete sequencing and annotation would seem necessary in order to distinguish avian specific genome organization from effects of scope and approach by the avian genome sequencing effort [33], as the currently available assemblies contain multiple long, isolated gene structure models, sometimes even spanning an entire assembly scaffold (suggestive of incomplete presumed intergenic space sequencing).

### iLoci provide a robust representation of the genome

Improvements in genome assemblies come at the expense of disrupting the sequencebased coordinate system typically used for annotating the location of genome features. Parsing an annotated genome into iLoci provides an alternative representation of the genome that is robust to assembly and annotation updates. We illustrate this use case with two model organism examples: (1) Comparing two annotation versions on the same *Arabidopsis thaliana* assembly; and (2) updated annotation on more complete genome assembly of the honey bee *Apis mellifera* compared to the original assembly and an earlier community annotation. For (1), the 2005 TAIR6 release was the first annotation of the *A. thaliana* genome managed by The Arabidopsis Information Resource [35], while the 2010 TAIR10 release integrates TAIR’s latest improvements to both the reference genome assembly and annotation using EST data from Sanger platforms [36].

For both species, we computed iLoci for each assembly/annotation version and determined *iLocus stability* as described in the **Methods** (**Additional file 1: Table S6**). **Figure 5** and **Additional file 1: Table S7** provide a breakdown of conservation by iLocus type.

**Figure 5:**
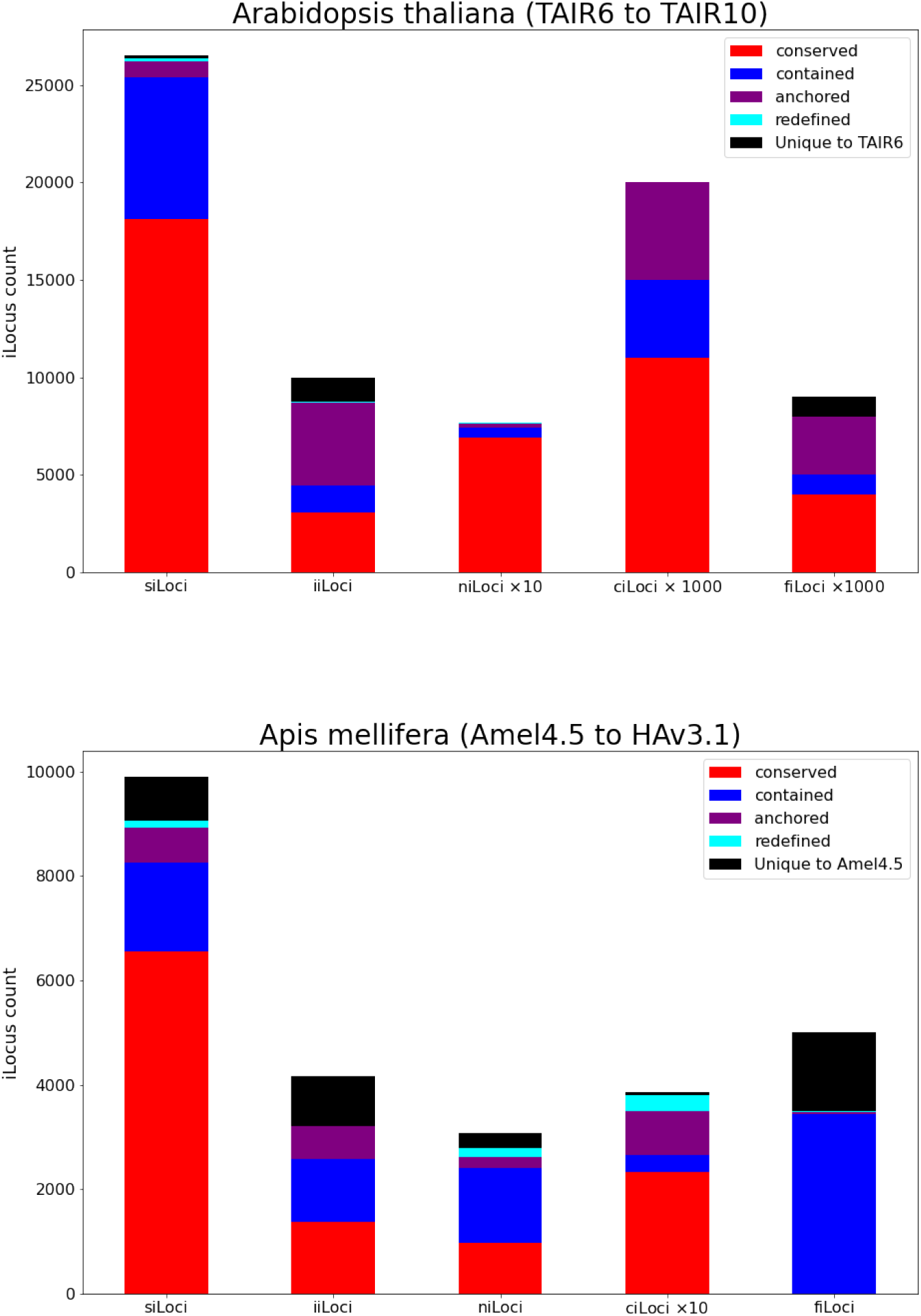
Breakdown of conservation per iLocus type. Note that the numbers of some iLocus types have been multiplied as indicated on the x-axis to allow visualization on the same plot.

The most obvious observation is that in both case studies the numbers of iLoci are fairly stable, except for easily explained changes. Thus, for the TAIR6 to TAIR10 comparison, new developments in publicly available RNA-Seq data and ncRNAs presented an opportunity to update the genome annotation, culminating in Araport11 [37], which has since been incorporated into the TAIR10 labeled annotation used here. As a result, we see a significant increase in ncRNA annotations and a modest increase in protein-coding genes (see **Additional file 1: Table S5**). Improvements in protein-coding gene annotations can be credited to incorporation of augmented depth of RNA-Seq data identifying novel transcript and splicing isoforms [37].

Figure 5 shows that very few siLoci are unique to TAIR6, indicating stability over many years of annotation updates (figure produced by **Additional file 2: notebooks/make-F5-F6.ipynb**). Non-conserved siLoci are mostly contained, i.e. embedded in longer iLoci in the current annotation. By contrast, non-conserved iiLoci are mostly anchored, i.e. the original iiLocus annotation was mapped to a shorter new iiLocus. TAIR6-unique gene models not transferred to TAIR10 tend to be short (**Figure 6**).

For *A. mellifera*, the Honey Bee Genome Sequencing Consortium’s assembly version Amel 2.0 and Official Gene Set 1 (OGSv1.0) were preliminary data resources in use prior to the initial published description of the honey bee genome in 2006 [6], while assembly Amel 4.5 (corresponding to NCBI release 102) and OGSv3.2 represent the consortium’s latest improvements to the genome and corresponding annotation as of 2014 [7, 8]. Release 103, still labeled Amel 4.5 [38], features some small differences from 102, such as a slight increase in the number of protein-coding genes, likely a result of newer gene annotation software. Release 104 (HAv3.1) is NCBI’s latest genome entry for *A. mellifera*, describing a new assembly derived from novel DNA sequencing technologies and, consequently, updated and revised annotations compared to Amel4.5. Unlike in the previous case, both annotations for *A. mellifera* were performed by the NCBI Eukaryotic Genome Annotation Pipeline, an automated pipeline for gene annotation, as part of [30].

**Figure 6:**
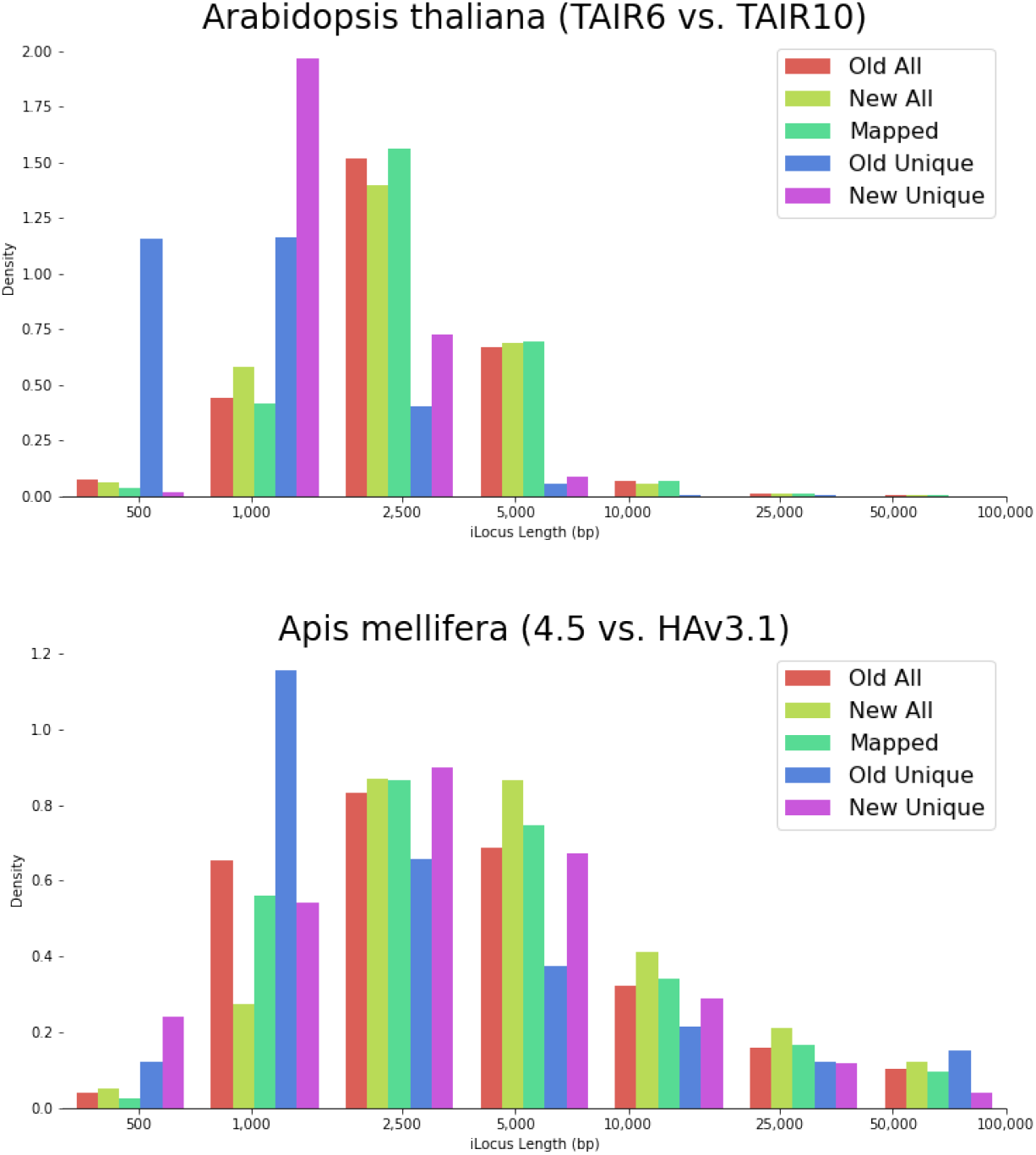
Breakdown of conservation per iLocus length bin.

A large number of annotation 4.5 siLoci are unmapped to assembly/annotation HAv3.1 (**Figure 5**). **Figure 6** shows that these are largely shorter gene structure models (and thus probably explained by gene model prediction algorithm parameter choices).

The main insight from these case studies is that the majority of iLoci can be faithfully mapped from one assembly/annotation pair to another. Practically, this suggests that iLoci identifiers can be used as database keys that point to entries containing both gene information and genome context information that will remain largely stable as assembly and annotation gaps are being filled.

## Discussion

Within the context of annotating a new genome, iLoci provide a quick and convenient solution for leveraging genomes of related model organisms to establish baseline expectations about genome composition and organization for the organism of interest. Similarities to genomes of related species across a broad range of measures gives one confidence in the quality of the genome assembly and annotation. By contrast, any stark differences should point to specific genomic features that warrant additional investigation to distinguish the effects of annotation from real differences in genome biology. Considerable effort has been devoted to making such comparisons as easy as possible: relevant software is freely available as open source code, is engineered with a focus on resource efficiency (enabling it to run easily on laptop or desktop computers), and works with a small number of standard input files. In short, iLoci provide a “common currency” for evaluating new data sets and re-evaluating previously published data sets alike.

Additional applications of iLoci in the annotation and analysis of novel genomes are numerous. Leveraging iLoci with strong support from expression and homology evidence to train species-specific gene prediction models can yield improvements in subsequent annotation efforts. The longest regions of the genome annotated as intergenic can yield insight into the proliferation of transposable and other repetitive elements and ncRNA genes, or alternatively characteristics of regions where annotation workflows fail to predict genes. The largest regions of high gene density, as represented by miLoci, provide an excellent starting place for investigating the clustering of functionally related genes, whereas miLoci containing two genes are candidates for genome-wide analysis of tandem gene duplication.

iLoci also facilitate analysis of genome organization at multiple scales. At the scale of whole chromosomes (or large fractions thereof), iLoci provide a well-defined measure of genome compactness that can be compared across annotations, assemblies, and species. At a slightly smaller scale, iLoci can be leveraged to investigate largescale changes in genome organization along the length of the chromosome, with possible interpretation in terms of transposon activity and other dynamic mechanisms of genome expansion and contraction. At the scale of individual genes, iLoci capture local aspects of genome organization, furnishing insight into gene spacing and orientation for specific genes of interest. Insight gained from analysis of genome organization at these various scales also lays a foundation for more detailed modeling of genome architecture, and perhaps even simulation of genome evolutionary dynamics. Simulating transposon activity, gene duplication, and genome rearrangements at various rates, and observing the effect these have on signatures of large-scale genome organization provided by iLoci could yield insight into the dominant mechanisms driving the evolution of genome architecture in particular species or clades of interest.

## Conclusions

Parsing annotated genome sequences into iLoci and then using these iLoci as a new coordinate system provides a robust and reproducible framework for investigating a variety of questions about genome content, architecture, and evolution. iLocus annotation might include contextual information for gene models in the form of up- and down-stream regulatory sequences. iLoci containing overlapping gene models can easily be identified for scrutiny seeking to distinguish gene model prediction errors from true compact gene organization that would likely be missed if analysis were performed at the level of individual genes. iLoci also provide stability across different versions of an annotated genome assembly, preserving gene models or intergenic regions for which local genomic context remained invariant to assembly and annotation updates. Finally, iLoci provide a way to break down the entire genome into distinct blocks that can be filtered based on their composition, gene content, conservation, or a variety of other characteristics of interest, thus providing finely tuned data sets for analyses or training and testing of predictive models.

## Supporting information

AdditionalFile2

## Author’s contributions

D.S.S. implemented the AEGeAn Toolkit, contributed to the initial study design, performed the initial studies of genome content, genome compactness, and iLocus stability, wrote early drafts of the manuscript, and edited the final version of the manuscript. T.L. updated and expanded the original data analysis and Python scripts. In addition, he designed and implemented the procedure to retrieve and analyze genome compactness characteristics for large sets of branch-specific genomes. V.P.B. conceived the concept of iLoci, designed the study, finalized the code and code repositories, and wrote drafts and most of the final version of the manuscript.

## Acknowledgements

The authors are grateful for use of the Extreme Science and Engineering Discovery Environment (XSEDE) Jetstream resource at Indiana University and the Texas Advanced Computing Center through allocation TG-BIO160012 (Computational Genomics) to V.B.. XSEDE is supported by National Science Foundation grant number ACI-1548562. We would like to thank our colleagues Robert Policastro and Haixu Tang for helpful comments on an earlier version of the manuscript.

## Supplementary Figures

**Figure S1:**
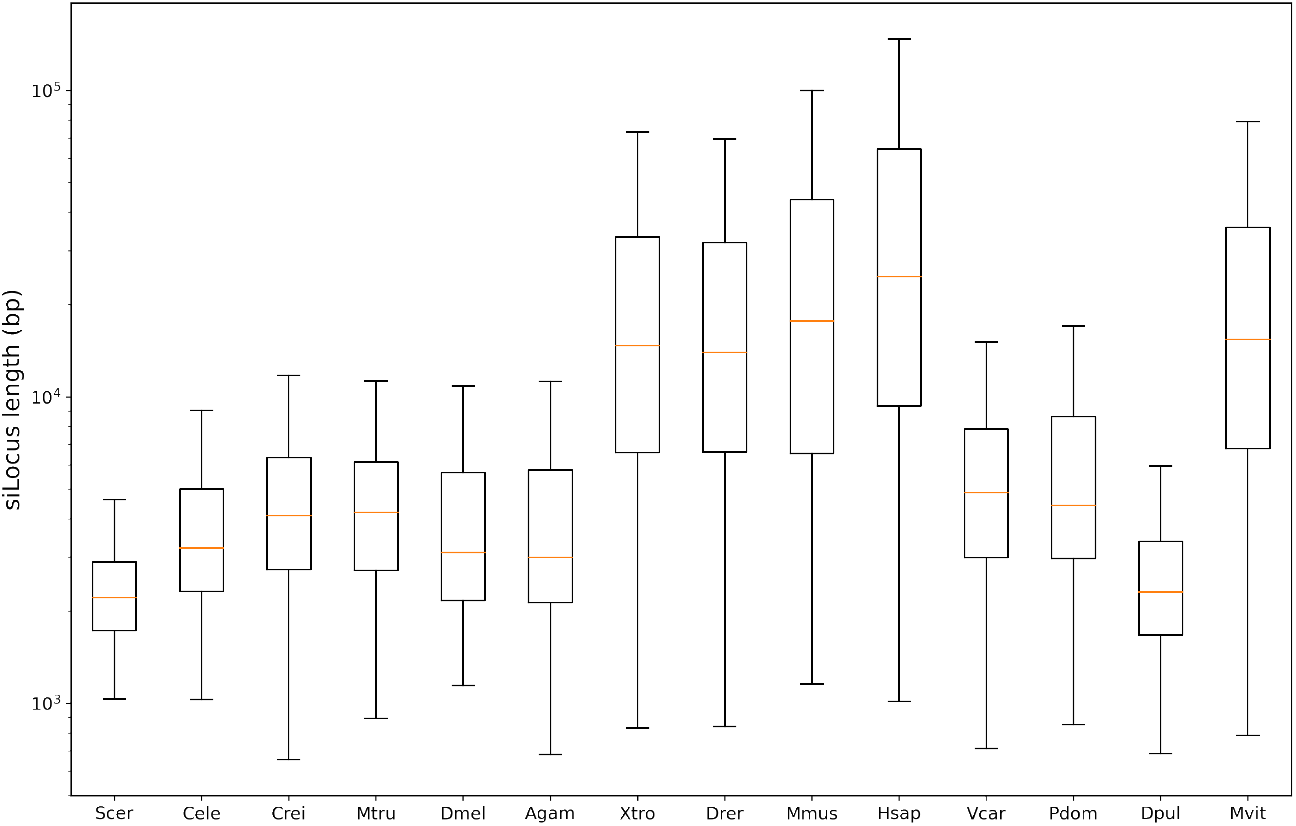
Distribution of siLocus lengths for genomes of ten model organisms.

**Figure S2:**
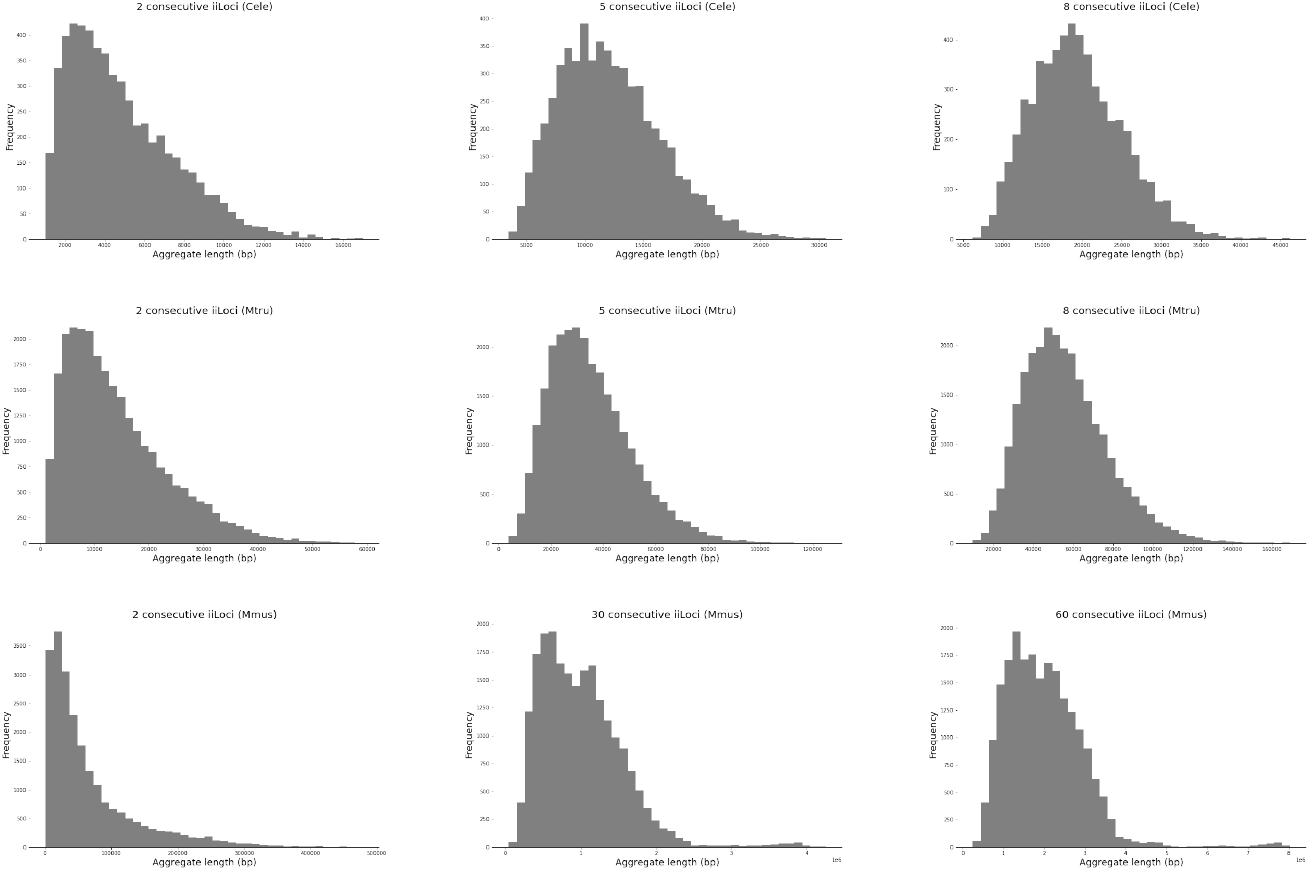
Aggregate lengths of *n* adjacent iiLoci for three species. The value of *n* at which the distribution of aggregate iiLocus lengths becomes near-normal is reflective of the range at which local variations in gene spacing are evened out.

**Figure S3:**
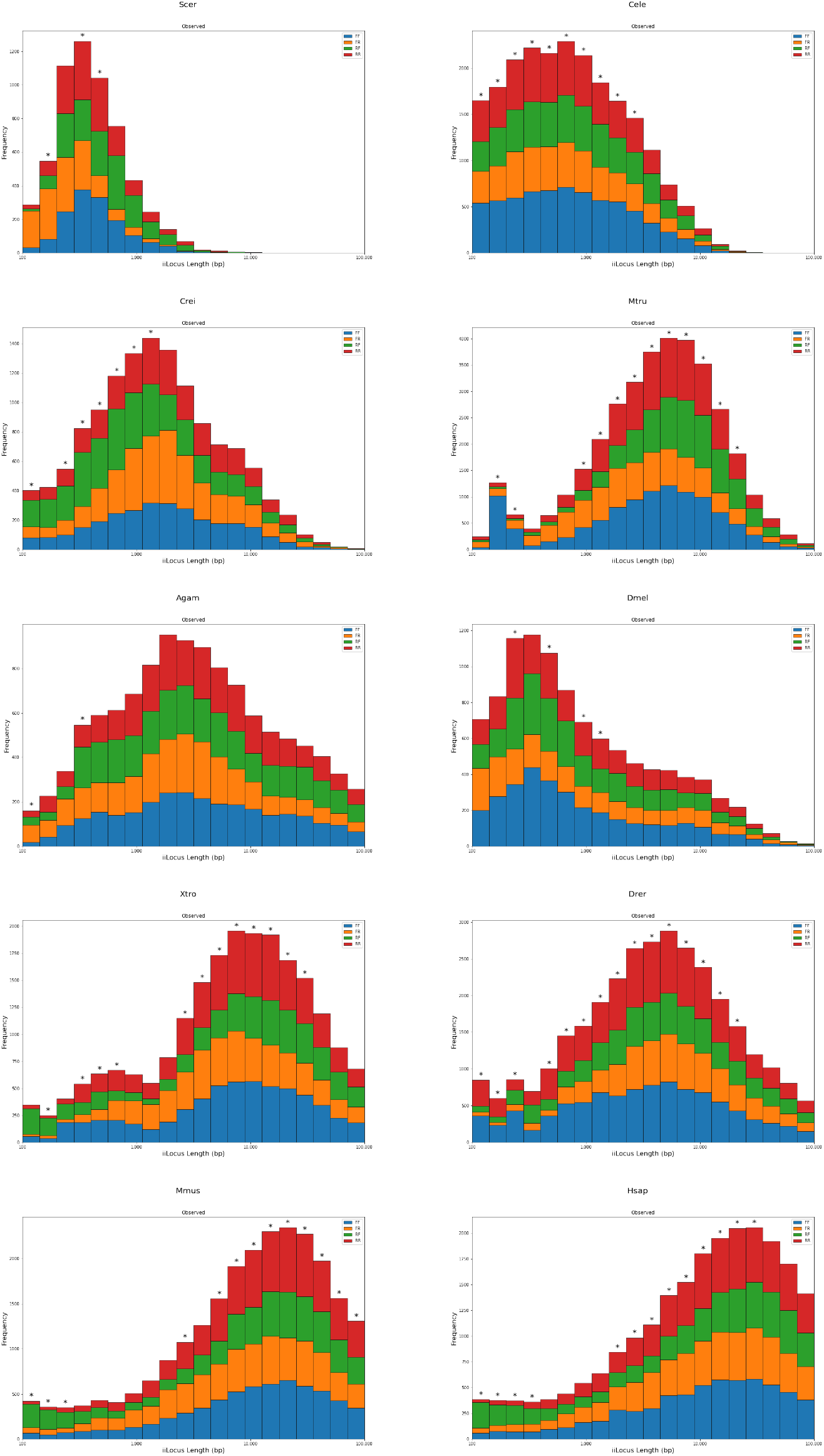
Histograms of frequencies of iiLocus lengths, broken down by the four orientation types. Starred length bins show significant deviation of counts from expectation based on Fisher’s exact test at the 1% significance level, adjusted for multiple testing.

**Figure S4:**
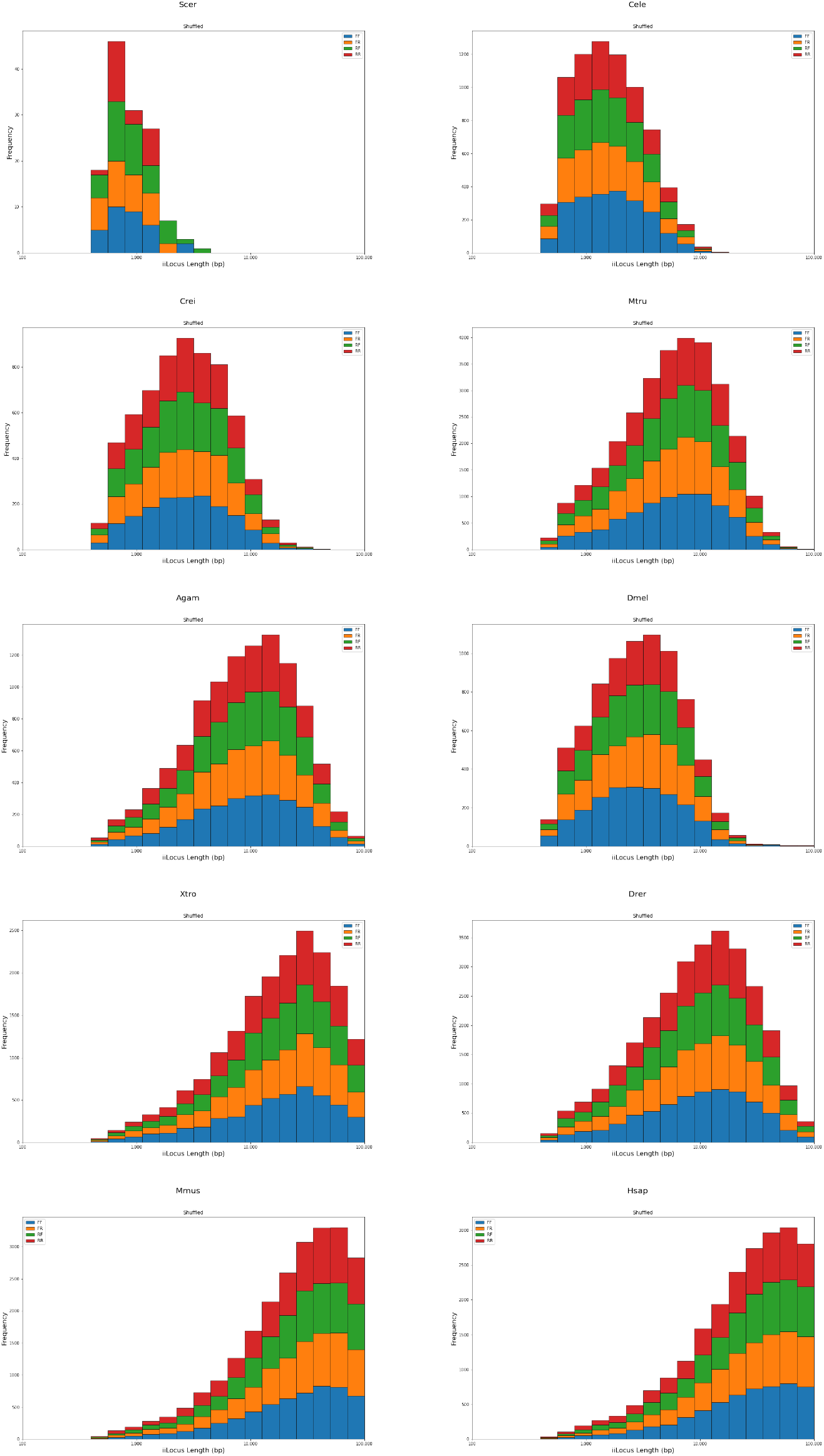
Histograms of frequencies of iiLocus lengths, broken down by the four orientation types, for randomized gene placement in the genome. Only chromosome/scaffold sequences of at least 1 Mb in length were used as input data.

**Figure S5:**
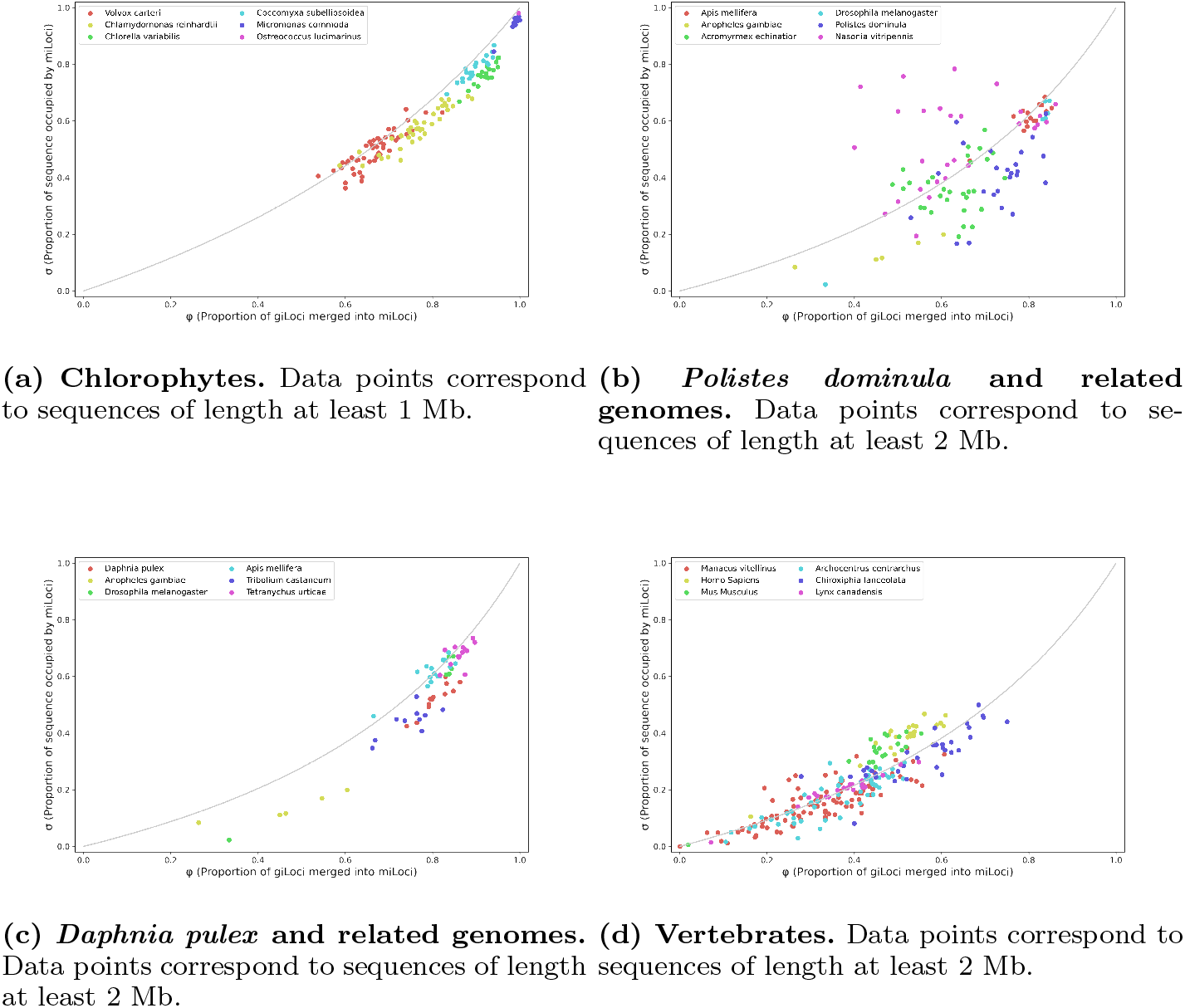
Genome compactness for different groups of species. In all plots, short giLoci (lower 5%-tile) and long iiLoci (top 5%-tile) were removed for each genome prior to calculation.

**Figure S6:**
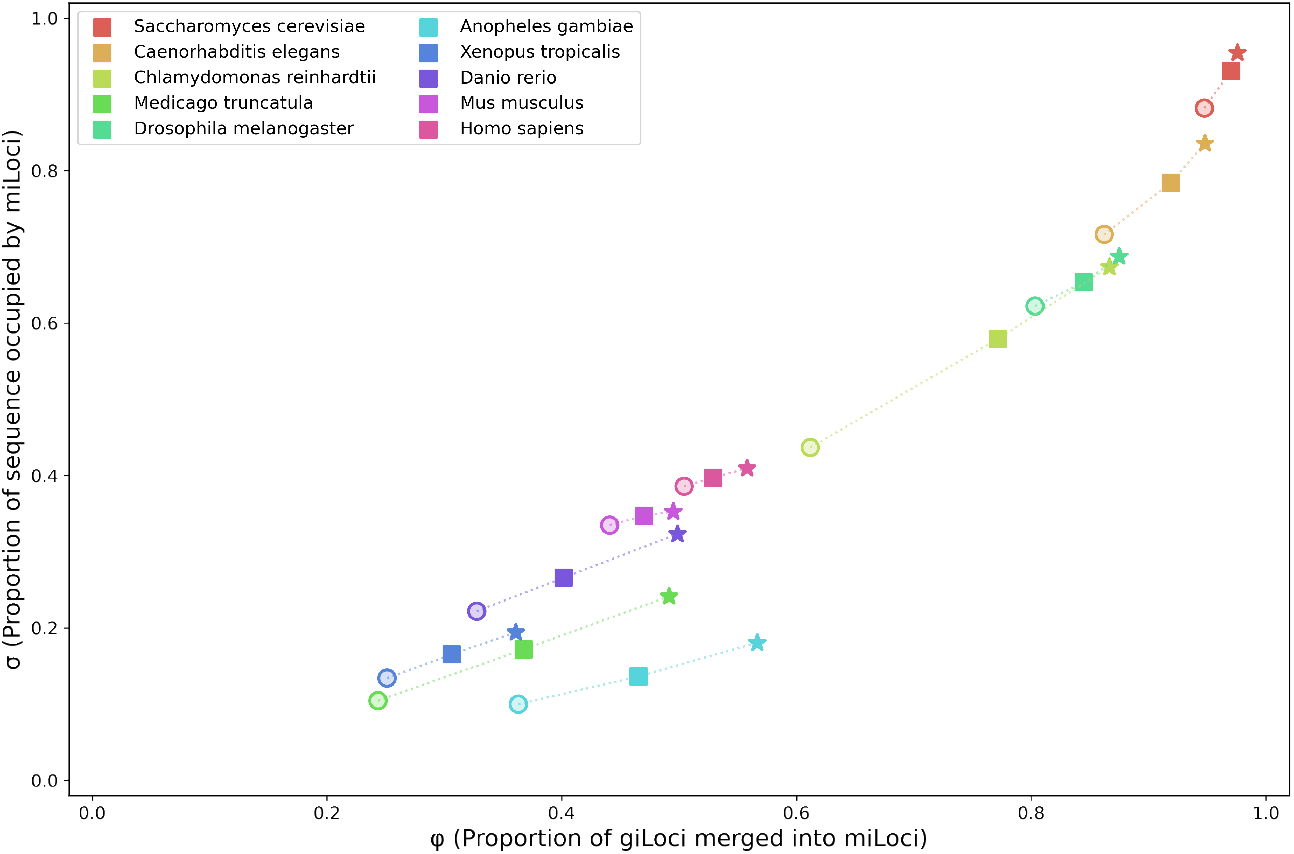
Comparison of (*ϕ, σ*) genome compactness with different values of *δ*. Circles, squares, and stars are for *δ* values 300, 500, or 750, respectively. Data points correspond to sequences of length at least 1 Mb. In all plots, short giLoci (lower 5%-tile) and long iiLoci (top 5%-tile) were removed for each genome prior to calculation.

**Figure S7:**
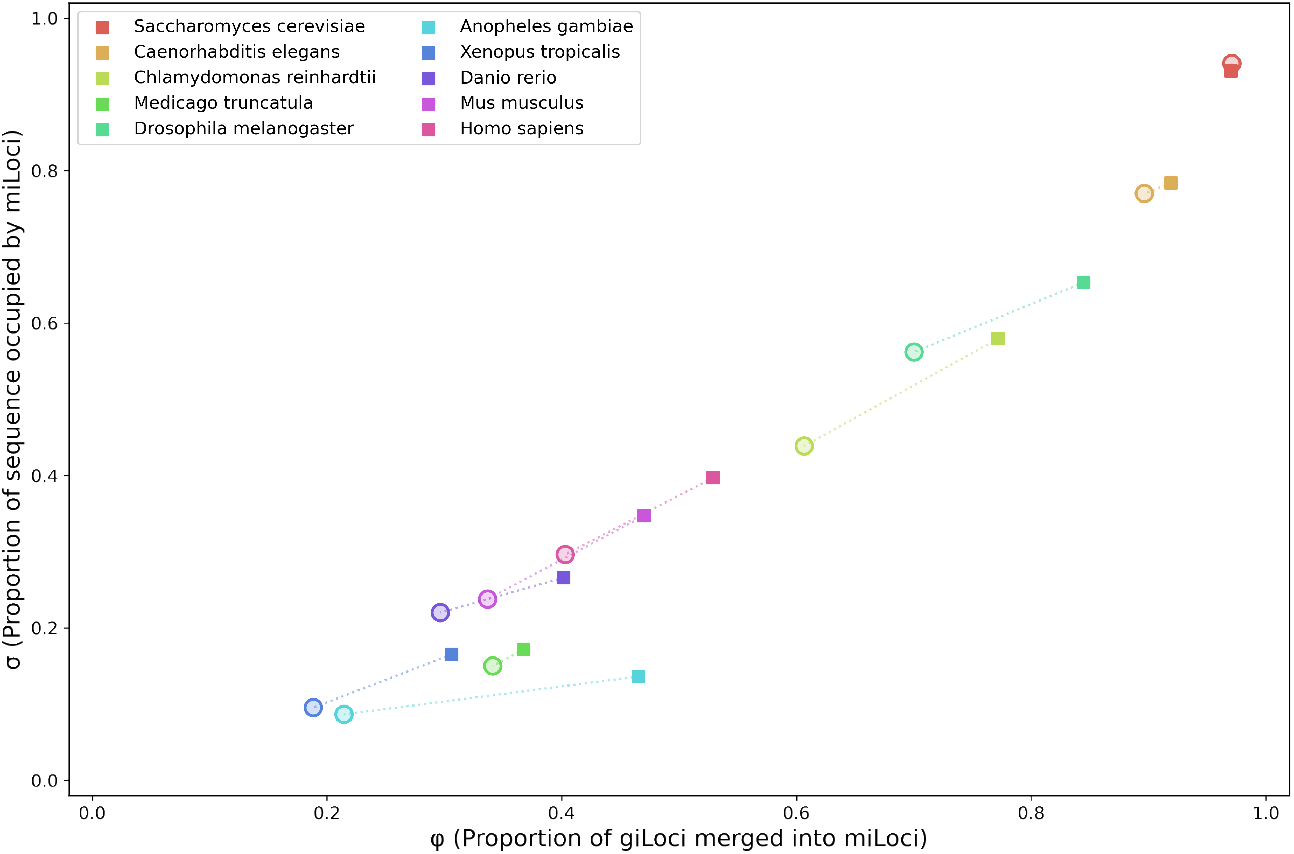
Comparison of (*ϕ, σ*) genome compactness as measured on genomic sequences as annotated (solid squares) versus the same sequences with a random arrangement of genes (hollow circles). Each point represents the average (centroid) of (*ϕ, σ*) values computed on all chromosome and scaffold sequences of lengths at least 1 Mb in the corresponding assembly. Short giLoci (lower 5%-tile) and long iiLoci (top 5%-tile) were removed for each genome prior to calculation.

**Figure S8:**
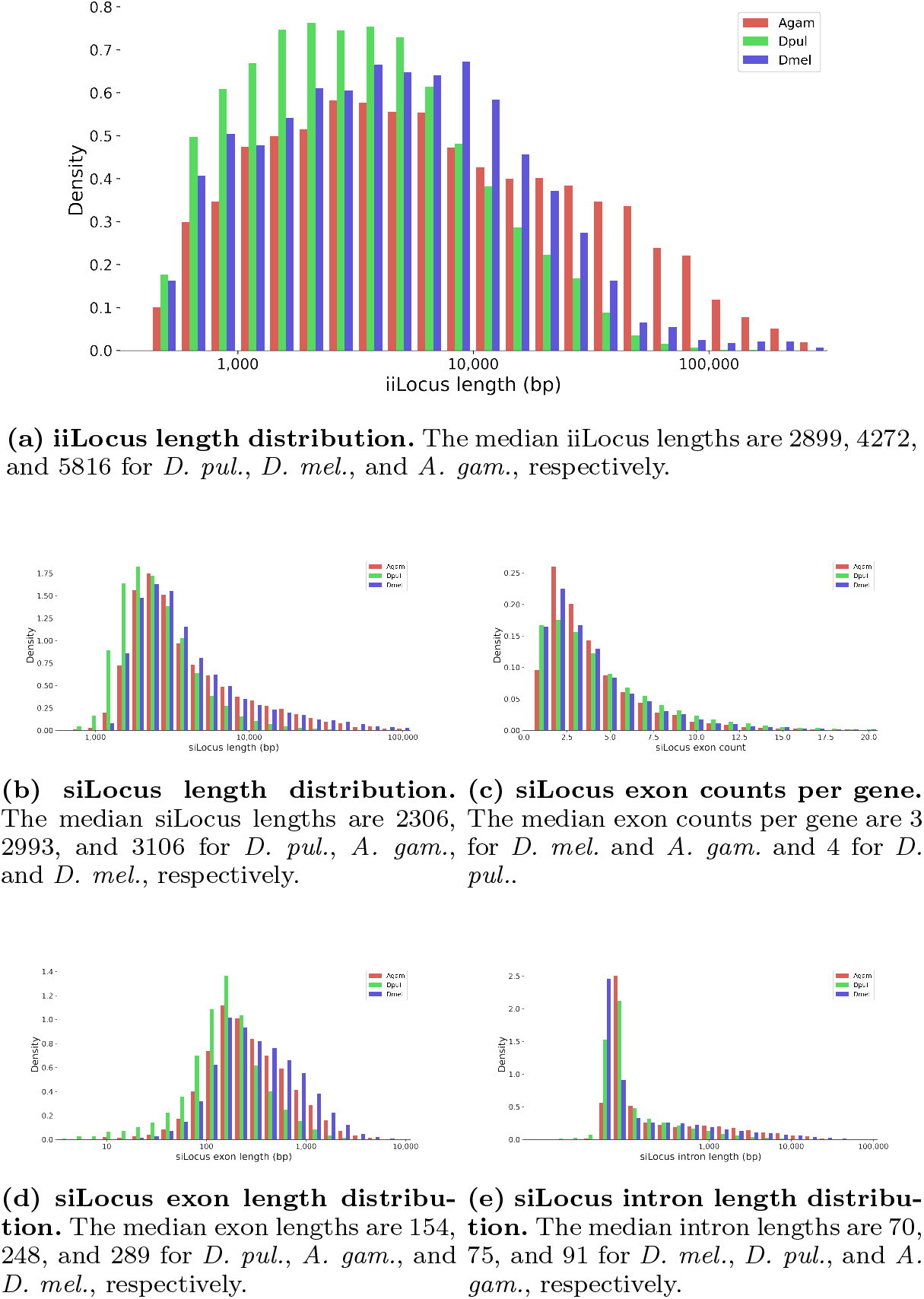
Evaluation of *Daphnia pulex* (green) genome compactness relative to *Drosophila melanogaster* (blue) and *Anopheles gambiae* (red).

## Supplementary Tables

**Table S1:** Summary of miLoci computed on randomly positioned genes.

| Species | miLoci | Occupancy<sup>1</sup> | Median #<br>genes<sup>2</sup> | Singletons<sup>3</sup> |
| --- | --- | --- | --- | --- |
| *Saccharomyces cerevisiae* | 158 | 4.5 Mb (93.1%) | 11 | 59 ( 2.4%) |
| *Caenorhabditis elegans* | 5,465 | 76.2 Mb (76.0%) | 4 | 2,672 ( 6.8%) |
| *Chlamydomonas reinhardtii* | 2,480 | 33.5 Mb (42.1%) | 2 | 3,978 (37.7%) |
| *Medicago truncatula* | 5,655 | 56.6 Mb (13.3%) | 2 | 24,562 (65.8%) |
| *Anopheles gambiae* | 1,322 | 18.3 Mb ( 8.0%) | 2 | 9,474 (76.4%) |
| *Drosophila melanogaster* | 3,724 | 68.7 Mb (50.0%) | 2 | 4,969 (29.9%) |
| *Xenopus tropicalis* | 1,952 | 126.1 Mb ( 9.8%) | 2 | 17,943 (80.2%) |
| *Danio rerio* | 4,675 | 271.9 Mb (20.3%) | 2 | 25,284 (64.4%) |
| *Mus musculus* | 4,800 | 548.1 Mb (20.1%) | 2 | 23,277 (66.4%) |
| *Homo sapiens* | 5,735 | 808.3 Mb (26.2%) | 2 | 21,285 (58.4%) |

**Table S2:** Summary of piLoci in genomes downloaded from NCBI, grouped by taxonomic branches.

Compare to Table 3.**Table S2:** Summary of piLoci in genomes downloaded from NCBI, grouped by taxonomic branches.
| Species | piLoci | Occupancy<sup>1</sup> | Single Exon piLoci |
| --- | --- | --- | --- |
| *Fungi* | 10,368 | 24.7 Mb (81.8%) | 2,619 (34.1%) |
| *Invertebrates* | 12,796 | 146.4 Mb (62.8%) | 1,197 ( 9.8%) |
| *Mammalian vertebrates* | 19,044 | 900.6 Mb (42.2%) | 1,787 ( 9.7%) |
| *Other vertebrates* | 16,032 | 490.9 Mb (59.6%) | 769 ( 4.7%) |
| *Plants* | 26,179 | 130.4 Mb (43.8%) | 4,570 (18.5%) |
| *Protozoa* | 8,266 | 23.3 Mb (79.5%) | 3,897 (50.9%) |

<sup>1</sup>Total number of nucleotides occupied by piLoci and the corresponding fraction of effective genome size.

**Table S3:** Summary of miLoci in genomes downloaded from NCBI, grouped by taxonomic branches.

| Species | miLoci | Occupancy<sup>1</sup> | Median # genes<sup>2</sup> | Singletons<sup>3</sup> |
| --- | --- | --- | --- | --- |
| *Fungi* | 1,644 | 22.6 Mb (76.0%) | 5 | 683 ( 7.5%) |
| *Invertebrates* | 2,306 | 72.9 Mb (33.5%) | 3 | 4,904 (38.9%) |
| *Mammalian vertebrates* | 3,374 | 305.6 Mb (15.0%) | 2 | 15,369 (64.6%) |
| *Other vertebrates* | 3,046 | 170.7 Mb (17.0%) | 2 | 12,880 (62.7%) |
| *Plants* | 5,403 | 69.4 Mb (30.1%) | 2 | 12,788 (42.8%) |
| *Protozoa* | 1,347 | 16.5 Mb (65.6%) | 3 | 1,181 (18.7%) |

<sup>1</sup>Total number of nucleotides occupied by miLoci and, in parentheses, the corresponding fraction of effective genome size.
<sup>2</sup>Median gene count per miLocus.
<sup>3</sup>Total number of giLoci not contained in miLoci and, in parentheses, corresponding fraction of all giLoci.

**Table S4:**
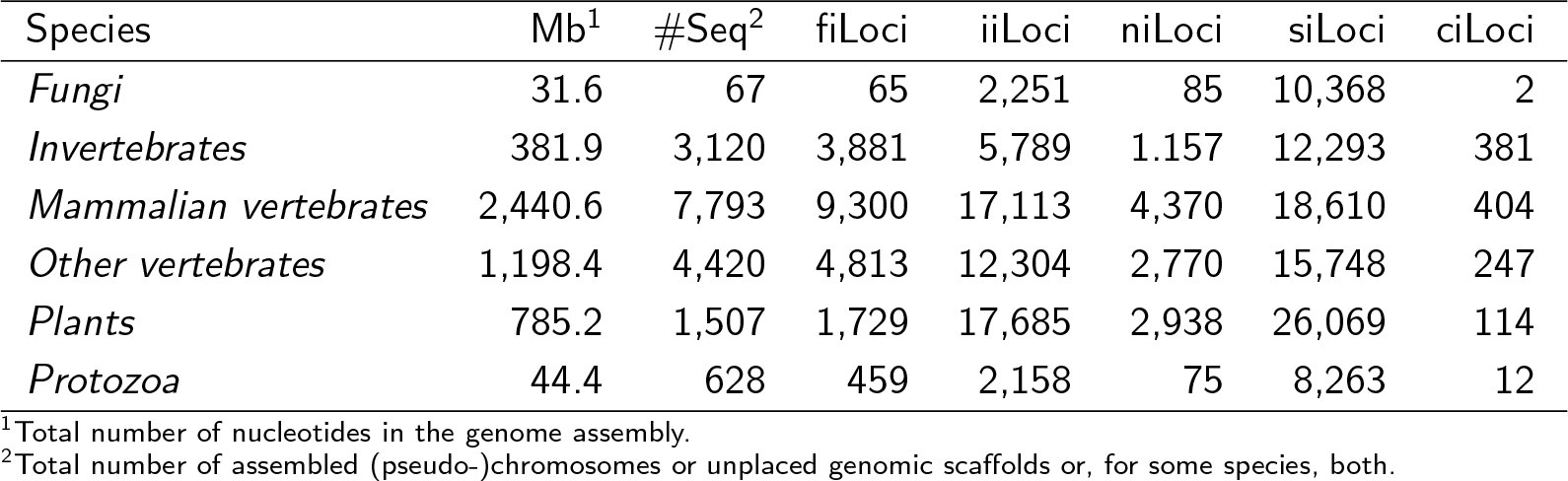
iLocus averages in genomes downloaded from NCBI, grouped by taxonomic branches.

| Species | Mb<sup>1</sup> | #Seq<sup>2</sup> | fiLoci | iiLoci | niLoci | siLoci | ciLoci |
| --- | --- | --- | --- | --- | --- | --- | --- |
| *Fungi* | 31.6 | 67 | 65 | 2,251 | 85 | 10,368 | 2 |
| *Invertebrates* | 381.9 | 3,120 | 3,881 | 5,789 | 1,157 | 12,293 | 381 |
| *Mammalian vertebrates* | 2,440.6 | 7,793 | 9,300 | 17,113 | 4,370 | 18,610 | 404 |
| *Other vertebrates* | 1,198.4 | 4,420 | 4,813 | 12,304 | 2,770 | 15,748 | 247 |
| *Plants* | 785.2 | 1,507 | 1,729 | 17,685 | 2,938 | 26,069 | 114 |
| *Protozoa* | 44.4 | 628 | 459 | 2,158 | 75 | 8,263 | 12 |

<sup>1</sup>Total number of nucleotides in the genome assembly.
<sup>2</sup>Total number of assembled (pseudo-)chromosomes or unplaced genomic scaffolds or, for some species, both.

**Table S5:**
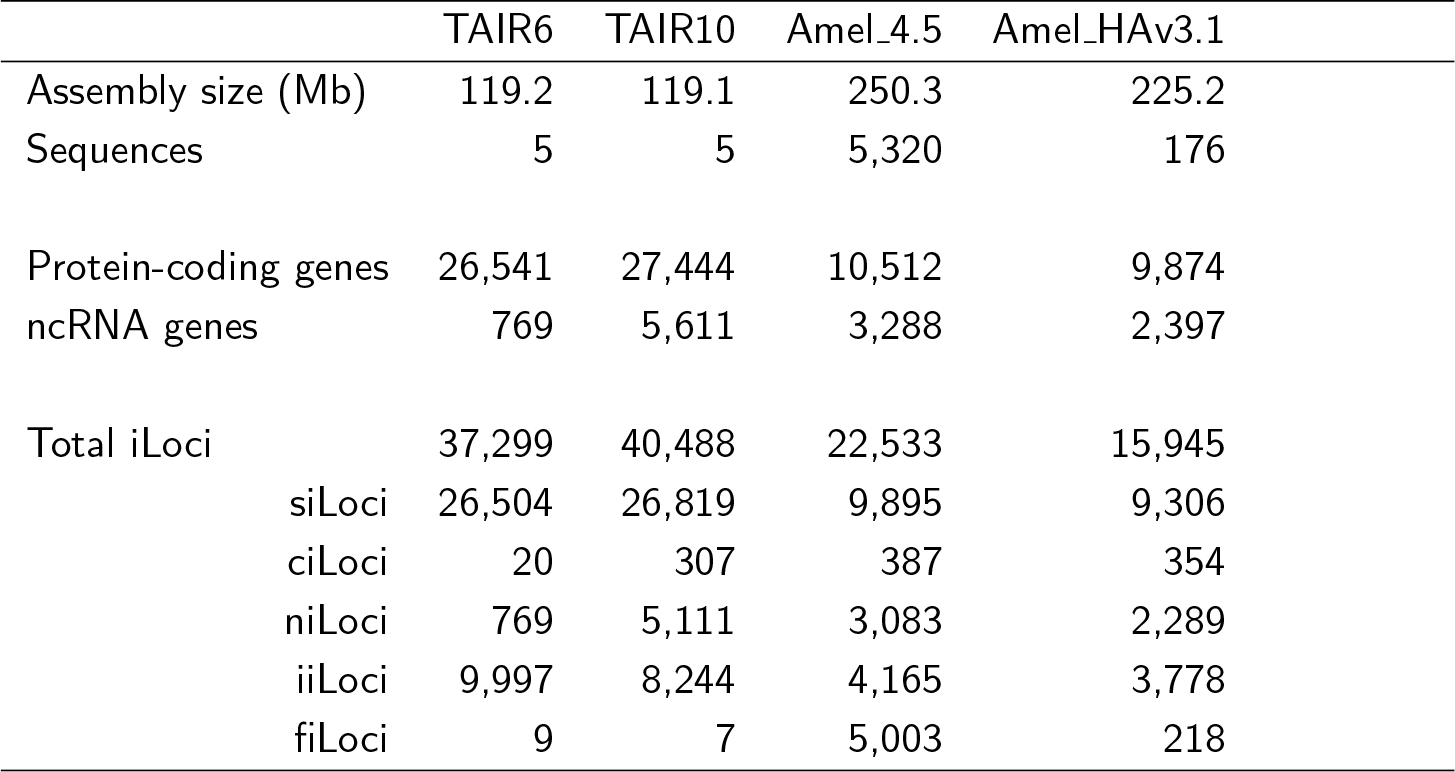
Descriptive summary of annotated genome assemblies for *A. thaliana* and *A. mellifera*.

**Table S6:** iLocus stability across assembly/annotation versions.

| *Arabidopsis thaliana* (TAIR6 → TAIR10) |  |  |  |  |  |  |  |  |  |
| --- | --- | --- | --- | --- | --- | --- | --- | --- | --- |
|  | all | woht | unmd | mapd | cons | uncn | cont | anch | redf |
| siLoci | 26,504 | 3 | 152 | 26,349 | 18,115 | 8,234 | 7,269 | 815 | 150 |
| niLoci | 769 | 0 | 2 | 767 | 690 | 77 | 52 | 21 | 4 |
| ciLoci | 20 | 0 | 0 | 20 | 11 | 9 | 4 | 5 | 0 |
| iiLoci | 9,997 | 13 | 1,210 | 8,774 | 3,085 | 5,689 | 1,354 | 4,266 | 69 |
| fiLoci | 9 | 0 | 1 | 8 | 4 | 4 | 1 | 3 | 0 |
| all iLoci | 37,299 | 16 | 1,365 | 35,918 | 21,905 | 14,013 | 8,680 | 5,110 | 223 |
| *Arabidopsis thaliana* (TAIR10 → TAIR6) |  |  |  |  |  |  |  |  |  |
|  | all | woht | unmd | mapd | cons | uncn | cont | anch | redf |
| siLoci | 26,819 | 2 | 668 | 26,149 | 17,989 | 8,160 | 1,489 | 6,645 | 26 |
| niLoci | 5,111 | 6 | 1,513 | 3,592 | 820 | 2,772 | 2,318 | 448 | 6 |
| ciLoci | 307 | 0 | 8 | 299 | 139 | 160 | 11 | 147 | 2 |
| iiLoci | 8,244 | 27 | 62 | 8,155 | 3,042 | 5,113 | 4,637 | 408 | 68 |
| fiLoci | 7 | 2 | 0 | 5 | 3 | 2 | 2 | 0 | 0 |
| all iLoci | 40,488 | 37 | 2,251 | 38,200 | 21,993 | 16,207 | 8,457 | 7,648 | 102 |
| *Apis mellifera* (4.5 → HAv3.1) |  |  |  |  |  |  |  |  |  |
|  | all | woht | unmd | mapd | cons | uncn | cont | anch | redf |
| siLoci | 9,895 | 280 | 551 | 9,064 | 6,549 | 2,515 | 1,708 | 665 | 142 |
| niLoci | 3,083 | 30 | 268 | 2,785 | 973 | 1,812 | 1,434 | 220 | 158 |
| ciLoci | 387 | 1 | 6 | 380 | 234 | 146 | 31 | 84 | 31 |
| iiLoci | 4,165 | 292 | 653 | 3,220 | 1,367 | 1,853 | 1,209 | 627 | 17 |
| fiLoci | 5,003 | 890 | 620 | 3,493 | 5 | 3,488 | 3,435 | 47 | 6 |
| all iLoci | 22,533 | 1,493 | 2,098 | 18,942 | 9,128 | 9,814 | 7,817 | 1,643 | 354 |
| *Apis mellifera* (HAv3.1 → 4.5) |  |  |  |  |  |  |  |  |  |
|  | all | woht | unmd | mapd | cons | uncn | cont | anch | redf |
| siLoci | 9,306 | 36 | 435 | 8,835 | 6,554 | 2,281 | 815 | 1,248 | 218 |
| niLoci | 2,289 | 15 | 158 | 2,116 | 983 | 1,133 | 701 | 213 | 219 |
| ciLoci | 354 | 1 | 3 | 350 | 232 | 118 | 14 | 86 | 18 |
| iiLoci | 3,778 | 315 | 354 | 3,109 | 1,356 | 1,753 | 876 | 780 | 97 |
| fiLoci | 218 | 36 | 35 | 147 | 2 | 145 | 2 | 125 | 18 |
| all iLoci | 15,945 | 403 | 985 | 14,557 | 9,127 | 5,430 | 2,408 | 2,452 | 570 |

Column abbreviations: woht, without hit; unmd, unmapped; mapd, mapped; cons, conserved; uncn, unconserved; cont, contained; anch, anchored; redf, redefined. Note that "all" comprises iLoci without hits, unmapped iLoci, and mapped iLoci. Mapped iLoci comprise conserved and unconserved iLoci. The unconserved category is comprised of contained, anchored, and redefined iLoci. See **Methods** for details.

**Table S7:** Breakdown of conservation type by iLoci class.

| *Arabidopsis thaliana* (TAIR6 → TAIR10) |  |  |  |  |  |  |
| --- | --- | --- | --- | --- | --- | --- |
|  | All | siLocus | niLocus | ciLocus | iiLocus | fiLocus |
| TAIR6 siLoci conserved | 18,115 | 17,939 | 38 | 132 | 6 | 0 |
| contained in | 7,269 | 6,437 | 25 | 178 | 629 | 0 |
| anchored by | 815 | 712 | 46 | 1 | 56 | 0 |
| redefined to | 150 | 129 | 2 | 2 | 17 | 0 |
| TAIR6 niLoci conserved | 690 | 0 | 690 | 0 | 0 | 0 |
| contained in | 52 | 8 | 43 | 0 | 1 | 0 |
| anchored by | 21 | 2 | 18 | 0 | 1 | 0 |
| redefined to | 4 | 3 | 0 | 1 | 0 | 0 |
| TAIR6 ciLoci conserved | 11 | 4 | 0 | 7 | 0 | 0 |
| contained in | 4 | 0 | 1 | 2 | 1 | 0 |
| anchored by | 5 | 5 | 0 | 0 | 0 | 0 |
| redefined to | 0 | 0 | 0 | 0 | 0 | 0 |
| TAIR6 iiLoci conserved | 3,085 | 21 | 32 | 1 | 3,031 | 0 |
| contained in | 1,354 | 471 | 291 | 21 | 571 | 0 |
| anchored by | 4,266 | 147 | 441 | 3 | 3,675 | 0 |
| redefined to | 69 | 17 | 2 | 1 | 49 | 0 |
| TAIR6 fiLoci conserved | 4 | 0 | 1 | 0 | 0 | 3 |
| contained in | 1 | 1 | 0 | 0 | 0 | 0 |
| anchored by | 3 | 1 | 1 | 0 | 0 | 1 |
| redefined to | 0 | 0 | 0 | 0 | 0 | 0 |
| *Apis mellifera* (4.5 → HAv3.1) |  |  |  |  |  |  |
|  | All | siLocus | niLocus | ciLocus | iiLocus | fiLocus |
| Am45 siLoci conserved | 6,549 | 6,479 | 15 | 51 | 4 | 0 |
| contained in | 1,708 | 1,449 | 25 | 153 | 78 | 3 |
| anchored by | 665 | 524 | 67 | 2 | 71 | 1 |
| redefined to | 142 | 114 | 5 | 22 | 0 | 1 |
| Am45 niLoci conserved | 973 | 4 | 958 | 6 | 5 | 0 |
| contained in | 1,434 | 886 | 169 | 105 | 257 | 17 |
| anchored by | 220 | 24 | 140 | 1 | 55 | 0 |
| redefined to | 158 | 140 | 3 | 13 | 0 | 2 |
| Am45 ciLoci conserved | 234 | 51 | 8 | 175 | 0 | 0 |
| contained in | 31 | 12 | 1 | 14 | 4 | 0 |
| anchored by | 84 | 57 | 10 | 9 | 8 | 0 |
| redefined to | 31 | 17 | 0 | 14 | 0 | 0 |
| Am45 iiLoci conserved | 1,367 | 22 | 0 | 1 | 1,344 | 0 |
| contained in | 1,209 | 375 | 65 | 42 | 722 | 5 |
| anchored by | 627 | 50 | 26 | 0 | 547 | 4 |
| redefined to | 17 | 14 | 0 | 1 | 2 | 0 |
| Am45 fiLoci conserved | 5 | 0 | 2 | 0 | 2 | 1 |
| contained in | 3,435 | 1,805 | 100 | 248 | 780 | 502 |
| anchored by | 47 | 5 | 6 | 0 | 30 | 6 |
| redefined to | 6 | 5 | 0 | 0 | 1 | 0 |

Rows correspond to the type annotated in the earlier annotation version, and columns correspond to the type annotated in the later version.

## Additional Files

Additional file 1 — Supplementary figures and tables

This PDF file contains all the supplementary figures and tables referred to in the text.

Additional file 2 — Workflow documentation, scripts, and notebooks

This file is a zip archive of our github repository https://github.com/BrendelGroup/iLoci_SLB21 frozen at the time of submission. The parent directory contains instructions on how to reproduce the data in this paper and how to modify and extend the scope of the computational work. Reference in the manuscript are relative to the *work* subdirectory.

## References

1. SRA: Sequence Read Archive. http://www.ncbi.nlm.nih.gov/sra

2. Stanke, M., Diekhans, M., Baertsch, R., Haussler, D.: Using native and syntenically mapped cDNA alignments to improve de novo gene finding. Bioinformatics 24(5), 637–644 (2008). doi:10.1093/bioinformatics/btn013

3. Campbell, M.S., Law, M., Holt, C., Stein, J.C., Moghe, G.D., Hufnagel, D.E., Lei, J., Achawanantakun, R., Jiao, D., Lawrence, C.J., Ware, D., Shiu, S.-H., Childs, K.L., Sun, Y., Jiang, N., Yandell, M.: MAKER-P: A tool kit for the rapid creation, management, and quality control of plant genome annotations. Plant Physiology 164(2), 513–524 (2014). doi:10.1104/pp.113.230144

4. Hoff, K.J., Lange, S., Lomsadze, A., Borodovsky, M., Stanke, M.: BRAKER1: Unsupervised RNA-Seq-based genome annotation with GeneMark-ET and AUGUSTUS. Bioinformatics (2015). doi:10.1093/bioinformatics/btv661

5. Souvorov, A., Kapustin, Y., Kiryutin, B., Chetvernin, V., Tatusova, T., Lipman, D.: Gnomon - NCBI Eukaryotic gene prediction tool (2010). http://www.ncbi.nlm.nih.gov/RefSeq/Gnomon-description.pdf

6. Elsik, C., Mackey, A., Reese, J., Milshina, N., Roos, D., Weinstock, G.: Creating a honey bee consensus gene set. Genome Biology 8(1), 13 (2007). doi:10.1186/gb-2007-8-1-r13

7. Elsik, C., Worley, K., Bennett, A., Beye, M., Camara, F., Childers, C., de Graaf, D., Debyser, G., Deng, J., Devreese, B., Elhaik, E., Evans, J., Foster, L., Graur, D., Guigo, R., production teams, H., Hoff, K., Holder, M., Hudson, M., Hunt, G., Jiang, H., Joshi, V., Khetani, R., Kosarev, P., Kovar, C., Ma, J., Maleszka, R., Moritz, R., Munoz-Torres, M., Murphy, T.: Finding the missing honey bee genes: lessons learned from a genome upgrade. BMC Genomics 15(1), 86 (2014). doi:10.1186/1471-2164-15-86

8. NCBI Apis mellifera Annotation Release 102. http://www.ncbi.nlm.nih.gov/genome/annotation_euk/Apis_mellifera/102/

9. Wallberg, A., Bunikis, I., Pettersson, O.V., Mosbech, M.-B., Childers, A.K., Evans, J.D., Mikheyev, A.S., Robertson, H.M., Robinson, G.E., Webster, M.T.: A hybrid de novo genome assembly of the honeybee, apis mellifera, with chromosome-length scaffolds. BMC Genomics 20(1), 275 (2019). doi:10.1186/s12864-019-5642-0

10. Standage, D.S.: The AEGeAn toolkit: Analysis and Evaluation of Genome Annotations. http://brendelgroup.github.io/AEGeAn/

11. Standage, D., Brendel, V.: ParsEval: parallel comparison and analysis of gene structure annotations. BMC Bioinformatics 13(1), 187 (2012). doi:10.1186/1471-2105-13-187

12. Riley, M.C., Clare, A., King, R.D.: Locational distribution of gene functional classes in Arabidopsis thaliana. BMC Bioinformatics 8, 112 (2007). doi:10.1186/1471-2105-8-112

13. Koonin, E.V.: Evolution of genome architecture. The International Journal of Biochemistry & Cell Biology 41(2), 298–306 (2009). doi:10.1016/j.biocel.2008.09.015

14. NCBI Genome. http://www.ncbi.nlm.nih.gov/genome/

15. Gremme, G., Steinbiss, S., Kurtz, S.: GenomeTools: A comprehensive software library for efficient processing of structured genome annotations. IEEE/ACM Transactions on Computational Biology and Bioinformatics 10(3), 645–656 (2013). doi:10.1109/TCBB.2013.68

16. GenomeTools web site. http://genometools.org/

17. Stein, L.: GFF3 Specification, The Sequence Ontology Project. http://www.sequenceontology.org/gff3.shtml

18. Wilbrandt, J., Misof, B., Panfilio, K.A., Niehuis, O.: Repertoire-wide gene structure analyses: a case study comparing automatically predicted and manually annotated gene models. BMC genomics 20(1), 753 (2019). doi:10.1186/s12864-019-6064-8

19. Eilbeck, K., Moore, B., Holt, C., Yandell, M.: Quantitative measures for the management and comparison of annotated genomes. BMC Bioinformatics 10(1), 67 (2009). doi:10.1186/1471-2105-10-67

20. GAEVAL: A Tool for Gene Annotation Evaluation. http://www.plantgdb.org/GAEVAL/docs/index.html

21. Franck, E., Hulsen, T., Huynen, M.A., de Jong, W.W., Lubsen, N.H., Madsen, O.: Evolution of closely linked gene pairs in vertebrate genomes. Molecular Biology and Evolution 25(9), 1909–1921 (2008). doi:10.1093/molbev/msn136

22. Harris, R.S.: Improved pairwise alignment of genomic dna. PhD thesis, The Pennsylvania State University (2007)

23. Li, W., Godzik, A.: Cd-hit: a fast program for clustering and comparing large sets of protein or nucleotide sequences. Bioinformatics 22(13), 1658–1659 (2006). doi:10.1093/bioinformatics/btl158

24. Pascual-Anaya, J., D’Aniello, S., Kuratani, S., Garcia-Fernàndez, J.: Evolution of Hox gene clusters in deuterostomes. BMC Developmental Biology 13(1), 1–15 (2013). doi:10.1186/1471-213X-13-26

25. Yi, G., Sze, S.-H., Thon, M.R.: Identifying clusters of functionally related genes in genomes. Bioinformatics 23(9), 1053–1060 (2007). doi:10.1093/bioinformatics/btl673

26. Howe, K., Clark, M.D., Torroja, C.F., Torrance, J., Berthelot, C., Muffato, M., Collins, J.E., Humphray, S., McLaren, K., Matthews, L., McLaren, S., Sealy, I., Caccamo, M., Churcher, C., Scott, C., Barrett, J.C., Koch, R., Rauch, G.-J., White, S., Chow, W., Kilian, B., Quintais, L.T., Guerra-Assuncao, J.A., Zhou, Y., Gu, Y., Yen, J., Vogel, J.-H., Eyre, T., Redmond, S., Banerjee, R., Chi, J., Fu, B., Langley, E., Maguire, S.F., Laird, G.K., Lloyd, D., Kenyon, E., Donaldson, S., Sehra, H., Almeida-King, J., Loveland, J., Trevanion, S., Jones, M., Quail, M., Willey, D., Hunt, A., Burton, J., Sims, S., McLay, K., Plumb, B., Davis, J., Clee, C., Oliver, K., Clark, R., Riddle, C., Eliott, D., Threadgold, G., Harden, G., Ware, D., Mortimer, B., Kerry, G., Heath, P., Phillimore, B., Tracey, A., Corby, N., Dunn, M., Johnson, C., Wood, J., Clark, S., Pelan, S., Griffiths, G., Smith, M., Glithero, R., Howden, P., Barker, N., Stevens, C., Harley, J., Holt, K., Panagiotidis, G., Lovell, J., Beasley, H., Henderson, C., Gordon, D., Auger, K., Wright, D., Collins, J., Raisen, C., Dyer, L., Leung, K., Robertson, L., Ambridge, K., Leongamornlert, D., McGuire, S., Gilderthorp, R., Griffiths, C., Manthravadi, D., Nichol, S., Barker, G., Whitehead, S., Kay, M., Brown, J., Murnane, C., Gray, E., Humphries, M., Sycamore, N., Barker, D., Saunders, D., Wallis, J., Babbage, A., Hammond, S., Mashreghi-Mohammadi, M., Barr, L., Martin, S., Wray, P., Ellington, A., Matthews, N., Ellwood, M., Woodmansey, R., Clark, G., Cooper, J., Tromans, A., Grafham, D., Skuce, C., Pandian, R., Andrews, R., Harrison, E., Kimberley, A., Garnett, J., Fosker, N., Hall, R., Garner, P., Kelly, D., Bird, C., Palmer, S., Gehring, I., Berger, A., Dooley, C.M., Ersan-Urun, Z., Eser, C., Geiger, H., Geisler, M., Karotki, L., Kirn, A., Konantz, J., Konantz, M., Oberlander, M., Rudolph-Geiger, S., Teucke, M., Osoegawa, K., Zhu, B., Rapp, A., Widaa, S., Langford, C., Yang, F., Carter, N.P., Harrow, J., Ning, Z., Herrero, J., Searle, S.M.J., Enright, A., Geisler, R., Plasterk, R.H.A., Lee, C., Westerfield, M., de Jong, P.J., Zon, L.I., Postlethwait, J.H., Nusslein-Volhard, C., Hubbard, T.J.P., Crollius, H.R., Rogers, J., Stemple, D.L.: The zebrafish reference genome sequence and its relationship to the human genome. Nature 496(7446), 498–503 (2013). doi:10.1038/nature12111

27. Karlin, S., Brendel, V.: Chance and statistical significance in protein and dna sequence analysis. Science 257(5066), 39–49 (1992). doi:10.1126/science.1621093. https://science.sciencemag.org/content/257/5066/39.full.pdf

28. Merchant, S.S., Prochnik, S.E., Vallon, O., Harris, E.H., Karpowicz, S.J., Witman, G.B., Terry, A., Salamov, A., Fritz-Laylin, L.K., Maréchal-Drouard, L., Marshall, W.F., Qu, L.-H., Nelson, D.R., Sanderfoot, A.A., Spalding, M.H., Kapitonov, V.V., Ren, Q., Ferris, P., Lindquist, E., Shapiro, H., Lucas, S.M., Grimwood, J., Schmutz, J., Cardol, P., Cerutti, H., Chanfreau, G., Chen, C.-L., Cognat, V., Croft, M.T., Dent, R., Dutcher, S., Fernández, E., Fukuzawa, H., González-Ballester, D., González-Halphen, D., Hallmann, A., Hanikenne, M., Hippler, M., Inwood, W., Jabbari, K., Kalanon, M., Kuras, R., Lefebvre, P.A., Lemaire, S.D., Lobanov, A.V., Lohr, M., Manuell, A., Meier, I., Mets, L., Mittag, M., Mittelmeier, T., Moroney, J.V., Moseley, J., Napoli, C., Nedelcu, A.M., Niyogi, K., Novoselov, S.V., Paulsen, I.T., Pazour, G., Purton, S., Ral, J.-P., Rianño-Pachón, D.M., Riekhof, W., Rymarquis, L., Schroda, M., Stern, D., Umen, J., Willows, R., Wilson, N., Zimmer, S.L., Allmer, J., Balk, J., Bisova, K., Chen, C.-J., Elias, M., Gendler, K., Hauser, C., Lamb, M.R., Ledford, H., Long, J.C., Minagawa, J., Page, M.D., Pan, J., Pootakham, W., Roje, S., Rose, A., Stahlberg, E., Terauchi, A.M., Yang, P., Ball, S., Bowler, C., Dieckmann, C.L., Gladyshev, V.N., Green, P., Jorgensen, R., Mayfield, S., Mueller-Roeber, B., Rajamani, S., Sayre, R.T., Brokstein, P., Dubchak, I., Goodstein, D., Hornick, L., Huang, Y.W., Jhaveri, J., Luo, Y., Martínez, D., Ngau, W.C.A., Otillar, B., Poliakov, A., Porter, A., Szajkowski, L., Werner, G., Zhou, K., Grigoriev, I.V., Rokhsar, D.S., Grossman, A.R.: The Chlamydomonas genome reveals the evolution of key animal and plant functions. Science 318(5848), 245–250 (2007). doi:10.1126/science.1143609

29. Prochnik, S.E., Umen, J., Nedelcu, A.M., Hallmann, A., Miller, S.M., Nishii, I., Ferris, P., Kuo, A., Mitros, T., Fritz-Laylin, L.K., Hellsten, U., Chapman, J., Simakov, O., Rensing, S.A., Terry, A., Pangilinan, J., Kapitonov, V., Jurka, J., Salamov, A., Shapiro, H., Schmutz, J., Grimwood, J., Lindquist, E., Lucas, S., Grigoriev, I.V., Schmitt, R., Kirk, D., Rokhsar, D.S.: Genomic analysis of organismal complexity in the multicellular green alga Volvox carteri. Science 329(5988), 223–226 (2010). doi:10.1126/science.1188800

30. RefSeq: NCBI Reference Sequence Database. http://www.ncbi.nlm.nih.gov/refseq/

31. Standage, D.S., Berens, A.J., Glastad, K.M., Severin, A.J., Brendel, V.P., Toth, A.L.: Genome, transcriptome and methylome sequencing of a primitively eusocial wasp reveal a greatly reduced dna methylation system in a social insect. Molecular Ecology 25(8), 1769–1784 (2016). doi:10.1111/mec.13578

32. Colbourne, J.K., Pfrender, M.E., Gilbert, D., Thomas, W.K., Tucker, A., Oakley, T.H., Tokishita, S., Aerts, A., Arnold, G.J., Basu, M.K., Bauer, D.J., Cáceres, C.E., Carmel, L., Casola, C., Choi, J.-H., Detter, J.C., Dong, Q., Dusheyko, S., Eads, B.D., Fröhlich, T., Geiler-Samerotte, K.A., Gerlach, D., Hatcher, P., Jogdeo, S., Krijgsveld, J., Kriventseva, E.V., Kültz, D., Laforsch, C., Lindquist, E., Lopez, J., Manak, J.R., Muller, J., Pangilinan, J., Patwardhan, R.P., Pitluck, S., Pritham, E.J., Rechtsteiner, A., Rho, M., Rogozin, I.B., Sakarya, O., Salamov, A., Schaack, S., Shapiro, H., Shiga, Y., Skalitzky, C., Smith, Z., Souvorov, A., Sung, W., Tang, Z., Tsuchiya, D., Tu, H., Vos, H., Wang, M., Wolf, Y.I., Yamagata, H., Yamada, T., Ye, Y., Shaw, J.R., Andrews, J., Crease, T.J., Tang, H., Lucas, S.M., Robertson, H.M., Bork, P., Koonin, E.V., Zdobnov, E.M., Grigoriev, I.V., Lynch, M., Boore, J.L.: The ecoresponsive genome of Daphnia pulex. Science 331(6017), 555–561 (2011). doi:10.1126/science.1197761

33. Zhang, G., Li, B., Li, C., Gilbert, M.T., Jarvis, E.D., Wang, J.: Comparative genomic data of the avian phylogenomics project. Gigascience, 789–804 (2014). doi:10.1186/2047-217X-3-26

34. Ncbi manacus vitellinus annotation release 103

35. The Arabidopsis Information Resource. http://www.arabidopsis.org

36. Lamesch, P., Berardini, T.Z., Li, D., Swarbreck, D., Wilks, C., Sasidharan, R., Muller, R., Dreher, K., Alexander, D.L., Garcia-Hernandez, M., Karthikeyan, A.S., Lee, C.H., Nelson, W.D., Ploetz, L., Singh, S., Wensel, A., Huala, E.: The Arabidopsis Information Resource (TAIR): improved gene annotation and new tools. Nucleic Acids Research 40(D1), 1202–1210 (2012). doi:10.1093/nar/gkr1090

37. Cheng, C.-Y., Krishnakumar, V., Chan, A.P., Thibaud-Nissen, F., Schobel, S., Town, C.D.: Araport11: a complete reannotation of the arabidopsis thaliana reference genome. The Plant Journal 89(4), 789–804 (2017). doi:10.1111/tpj.13415. https://onlinelibrary.wiley.com/doi/pdf/10.1111/tpj.13415

38. Ncbi apis mellifera annotation release 103

