## AdditionalFile2 for "iLoci: Robust evaluation of genome content and organization for provisional and mature genome assemblies": README.md

# iLoci_SLB21
Procedures and data for the iLoci study described in Lai, Standage, and
Brendel (2021)


## Overview

This repository documents application of
[AEGeAn](https://github.com/BrendelGroup/AEGeAn) code to derive, classify, and
analyze genomic interval loci (iLoci) as described in the paper
Standage, Lai, and Brendel (2021).
Please refer to the
[AEGeAn Installation](https://github.com/BrendelGroup/AEGeAn/blob/master/INSTALL.md)
site for details on how to obtain the relevant software.
The simplest way to get going is to use the AEGeAn
[Singularity](https://sylabs.io/docs/) container, e.g. as follows:

```bash
cd
git clone https://github.com/BrendelGroup/iLoci_SLB21
cd iLoci_SLB21/work
wget https://BrendelGroup.org/SingularityHub/aegean.sif
alias rws="singularity exec -e -B ~/iLoci_SLB21 ~/iLoci_SLB21/work/aegean.simg"
rws fidibus -h
```

In the above example, you clone this repository into your Linux home directory,
go into the iLoci_SLB21/work directory that has been created, download the AEGeAn
Singularity container, define the shell variable _rws_ ("run with singularity"),
and check that everything works by showing the "help" lines for the AEGeAn
command _fididbus_.

To reproduce the data discussed in Standage, Lai, and Brendel (2021), go to
[./work](./work) and follow the instructions in the [README](./work/README.md)
file.
