## AdditionalFile2 for "iLoci: Robust evaluation of genome content and organization for provisional and mature genome assemblies": det-Chlorophyta-status-post-blast.py

```
#!/usr/bin/env python
from __future__ import print_function
import argparse


def get_parser():
    parser = argparse.ArgumentParser()
    parser.add_argument('blast_match_ids', type=argparse.FileType('r'))
    parser.add_argument('status', type=argparse.FileType('r'),
                        help='hilocus status table')
    return parser


def main(args):
    blast_matches = dict()
    for line in args.blast_match_ids:
        protid = line.strip()
        blast_matches[protid] = True

    for line in args.status:
        values = line.strip().split('\t')
        status = values[2]
        protid = values[4]
        if status == 'Unmatched':
            if protid in blast_matches:
                values[2] = 'Matched'
            else:
                values[2] = 'Orphan'
        print(*values, sep='\t')

if __name__ == '__main__':
    main(get_parser().parse_args())
```
