## AdditionalFile2 for "iLoci: Robust evaluation of genome content and organization for provisional and mature genome assemblies": det-Chlorophyta-status.py

```
#!/usr/bin/env python
from __future__ import print_function
import argparse
import glob


iloc2prot = dict()
mapfiles = ('data/Vcar/Vcar.protein2ilocus.tsv', \
        'data/Apro/Apro.protein2ilocus.tsv',
        'data/Crei/Crei.protein2ilocus.tsv',
        'data/Csub/Csub.protein2ilocus.tsv',
        'data/Cvar/Cvar.protein2ilocus.tsv',
        'data/Mcom/Mcom.protein2ilocus.tsv',
        'data/Mpus/Mpus.protein2ilocus.tsv',
        'data/Oluc/Oluc.protein2ilocus.tsv',
        'data/Otau/Otau.protein2ilocus.tsv',
        'data/Mtru/Mtru.protein2ilocus.tsv',
        'data/Atha/Atha.protein2ilocus.tsv',
        'data/Bdis/Bdis.protein2ilocus.tsv',
        'data/Osat/Osat.protein2ilocus.tsv' )
repfiles = ('data/Vcar/Vcar.protids.txt', \
        'data/Apro/Apro.protids.txt',
        'data/Crei/Crei.protids.txt',
        'data/Csub/Csub.protids.txt',
        'data/Cvar/Cvar.protids.txt',
        'data/Mcom/Mcom.protids.txt',
        'data/Mpus/Mpus.protids.txt',
        'data/Oluc/Oluc.protids.txt',
        'data/Otau/Otau.protids.txt',
        'data/Mtru/Mtru.protids.txt',
        'data/Atha/Atha.protids.txt',
        'data/Bdis/Bdis.protids.txt',
        'data/Osat/Osat.protids.txt' )
for mapfile, repfile in zip(mapfiles, repfiles):
    with open(mapfile, 'r') as mapstream, open(repfile, 'r') as repstream:
        reps = dict()
        for line in repstream:
            protid = line.strip()
            reps[protid] = True
        for line in mapstream:
            protid, locusid = line.strip().split('\t')
            if protid in reps:
                iloc2prot[locusid] = protid

chlorophytes = ['Vcar', 'Apro', 'Crei', 'Csub', 'Cvar', \
                'Mcom', 'Mpus', 'Oluc', 'Otau']

class HiLocus(object):

    def __init__(self, record):
        self._rawdata = record
        values = record.strip().split('\t')
        self.iloci = values[2].split(',')
        self.species = values[3].split(',')

    def status(self):
        assert len(self.species) >= 1 and len(self.species) <= 13
        chlorophyte_count = 0
        nonchlorophyte_count = 0
        for species in self.species:
            if species in chlorophytes:
                chlorophyte_count += 1
            else:
                nonchlorophyte_count += 1
        assert chlorophyte_count <= 9 and nonchlorophyte_count <= 4 and chlorophyte_count + nonchlorophyte_count > 0

        if chlorophyte_count == 9:
            return 'HighlyConserved'
        elif chlorophyte_count > 3:
            return 'Conserved'
        elif chlorophyte_count + nonchlorophyte_count > 1:
            return 'Matched'
        else:
            return 'Unmatched'


def get_parser():
    parser = argparse.ArgumentParser()
    parser.add_argument('hiloci', type=argparse.FileType('r'),
                        help='hilocus data table')
    return parser


def main(args):
    hilocus_count = 0
    print('hiLocus', 'iLocus', 'Status', 'Species', 'Protein', sep='\t')
    for line in args.hiloci:
        hilocus = HiLocus(line)
        hilocus_count += 1
        hid = 'HILC-{}'.format(hilocus_count)
        for locusid in hilocus.iloci:
            print(hid, locusid, hilocus.status(), locusid[0:4],
                  iloc2prot[locusid], sep='\t')

if __name__ == '__main__':
    main(get_parser().parse_args())
```
