## AdditionalFile2 for "iLoci: Robust evaluation of genome content and organization for provisional and mature genome assemblies": hiLoci-breakdown.py

```
#!/usr/bin/env python
from __future__ import print_function
import argparse
import sys


def status(histat):
    ilocus_status = dict()
    next(histat)
    for record in histat:
        values = record.strip().split('\t')
        assert len(values) == 5, values
        ilocus = values[1]
        status = values[2]
        ilocus_status[ilocus] = status
    return ilocus_status


def get_parser():
    """Define the program command-line interface."""
    parser = argparse.ArgumentParser()
    parser.add_argument('-c', '--counts', action='store_true',
                        help='report iLocus counts; by default, the amount of '
                        'the genome occupied (in bp) is reported')
    parser.add_argument('iloci', type=argparse.FileType('r'),
                        help='ilocus data table')
    parser.add_argument('histat', type=argparse.FileType('r'),
                        help='table of hiLocus conservation status')
    return parser


def main(args):
    """Main procedure."""
    ilocus_status = status(args.histat)
    outcols = ['HighlyConserved', 'Conserved', 'Matched', 'Orphan', 'Complex',
               'ncRNA', 'Intergenic', 'Fragment']
    breakdown = dict()
    for record in args.iloci:
        if record.startswith('Species'):
            continue
        values = record.rstrip().split('\t')
        species = values[0]
        ilcid = values[1]
        eff_len = int(values[5])
        ilclass = values[9]

        if species not in breakdown:
            breakdown[species] = dict((col, list()) for col in outcols)

        if ilclass == 'iiLocus':
            breakdown[species]['Intergenic'].append(values)
        elif ilclass == 'fiLocus':
            breakdown[species]['Fragment'].append(values)
        elif ilclass == 'ciLocus':
            breakdown[species]['Complex'].append(values)
        elif ilclass == 'niLocus':
            breakdown[species]['ncRNA'].append(values)
        else:
            assert ilclass == 'siLocus', ilclass
            if ilcid not in ilocus_status:
                print('warning: error determining conservation status for '
                      'iLocus {}'.format(ilcid), file=sys.stderr)
                continue
            stat = ilocus_status[ilcid]
            breakdown[species][stat].append(values)

    print('\t'.join(['Species'] + outcols))
    for species in sorted(breakdown):
        print(species, end='', sep='')
        for col in outcols:
            iloci = breakdown[species][col]
            if args.counts:
                print('\t%d' % len(iloci), end='', sep='')
            else:
                cumlength = sum([int(x[5]) for x in iloci])
                print('\t%d' % cumlength, end='', sep='')
        print('\n', end='')


if __name__ == '__main__':
    main(get_parser().parse_args())
```
