## AdditionalFile2 for "iLoci: Robust evaluation of genome content and organization for provisional and mature genome assemblies": README.md

# Annotation/Assembly comparisons

We investigate the transition of iLoci between assembly and annotation versions for two genomes: Apis mellifera and Arabidopsis thaliana. 

If the data files are not present, they may be downloaded and processed with the following:

```bash
Amels=(Amel Amh3)

fidibus --numprocs=3 --refr=Amel,Amh3,Atha download prep iloci breakdown stats
for name in ${Amels[@]}
    mv species/${species}/${species}.iloci.tsv Amel/ 
    mv species/${species}/${species}.iloci.fa Amel/
done
mv species/Atha/Atha.iloci.tsv Atha/
mv species/Atha/Atha.iloci.fa Atha/
mv species/Atha/GCF_000001735.3_TAIR10_genomic.fna Atha/
```
The second version of Arabidopsis is from Araport and requires local downloading and processing. 
Obtain the genome annotation (GFF3 file) and the protein FASTA file and place them in the Atha directory. Then process with the xprepAt11 and run with xAt11 scripts. 

Part of the analysis involves a breakdown of the query iLocus types. Since this functionality is not currently present in fidibus, we do so with a custom Python script.

```bash
cd Amel/
python parse_loci.py
cd ../Atha
python parse_loci.py
```

Now that we have the data files, we may begin the comparison analysis. First, we compute chain alignments using LASTZ and postprocess with some custom 
Python scripts. 

We begin by calling LASTZ with a command such as:
```bash
lastz Amh3.iloci.fa[multiple] Amel.iloci.fa --match=1,9 --filter=identity:95 --chain \
		  format=general:name1,length1,size1,name2,length2,size2,identity,nmatch \
		  > entire.tsv
```
To run the entire analysis broken down by iLocus type, run the corresponding chain.sh script in each subdirectory. 
This generates a TSV file containing the query iLoci and all of the chains against the target, along with the attributes of both. We are mainly interested in the maximal chain lengths for each query. Additional python scripts 
then processes the TSV files and produces the relevant counts. These may be done by calling hsp.sh and then counts.sh

See [comparisons_plot.ipynb](https://github.com/timlai4/IntervalLoci/blob/comparisons/compare/comparison_plots.ipynb) for visualizations of these data.
