## AdditionalFile2 for "iLoci: Robust evaluation of genome content and organization for provisional and mature genome assemblies": runAmel.sh

```
#!/bin/bash
#
set -e
set -u
set -o pipefail

#1) Data set preparation:
#
mkdir Amel
labels=(Am45 Amel)

fidibus --numprocs=2 --refr=Am45,Amel download prep iloci breakdown stats

for species in ${labels[@]}
do
    ln -s ../species/${species}/${species}.iloci.tsv  Amel/ 
    ln -s ../species/${species}/${species}.iloci.fa   Amel/
    ln -s ../species/${species}/${species}.iloci.gff3 Amel/
done

cd ./Amel
makeblastdb -in Am45.iloci.fa -dbtype nucl -out Am45ILC -parse_seqids
makeblastdb -in Amel.iloci.fa -dbtype nucl -out AmelILC -parse_seqids
python3 ../scripts/split_iloci_by_type.py Am45
python3 ../scripts/split_iloci_by_type.py Amel
cd ..


#2) Running lastz:
#
cd ./Amel

# Assuming we have at least 5 processors, we run each lastz job in the background:
#
lastz Amel.iloci.fa[multiple] Am45.filoci.fa --allocate:traceback=500M \
		--ambiguous=iupac --filter=identity:95 --chain \
		--format=general:name1,length1,size1,name2,length2,size2,identity,nmatch \
		> Am45.filoci-vs-Amel.lastz &
lastz Amel.iloci.fa[multiple] Am45.ciloci.fa --allocate:traceback=500M \
		--ambiguous=iupac --filter=identity:95 --chain \
		--format=general:name1,length1,size1,name2,length2,size2,identity,nmatch \
		> Am45.ciloci-vs-Amel.lastz &
lastz Amel.iloci.fa[multiple] Am45.niloci.fa --allocate:traceback=500M \
		--ambiguous=iupac --filter=identity:95 --chain \
		--format=general:name1,length1,size1,name2,length2,size2,identity,nmatch \
		> Am45.niloci-vs-Amel.lastz &
lastz Amel.iloci.fa[multiple] Am45.iiloci.fa --allocate:traceback=500M \
		--ambiguous=iupac --filter=identity:95 --chain \
		--format=general:name1,length1,size1,name2,length2,size2,identity,nmatch \
		> Am45.iiloci-vs-Amel.lastz &
lastz Amel.iloci.fa[multiple] Am45.siloci.fa --allocate:traceback=500M \
		--ambiguous=iupac --filter=identity:95 --chain \
		--format=general:name1,length1,size1,name2,length2,size2,identity,nmatch \
		> Am45.siloci-vs-Amel.lastz &
wait
cd ..


#3) Processing the results:
#
iltypes=(filoci ciloci iiloci niloci siloci)

cd ./Amel
for iltype in ${iltypes[@]}
do
  n=`cat Am45.${iltype}.fa | egrep "^>" | wc -l`
  python3 ../scripts/process_lastz_output.py -n $n -i Am45.${iltype}-vs-Amel.lastz -v -o Am45.${iltype}-vs-Amel.summary
done
cd ..


###Reciprocal runs. Let's see whether we need them.===

cd ./Amel

# Assuming we have at least 5 processors, we run each lastz job in the background:
#
lastz Am45.iloci.fa[multiple] Amel.filoci.fa --allocate:traceback=500M \
		--ambiguous=iupac --filter=identity:95 --chain \
		--format=general:name1,length1,size1,name2,length2,size2,identity,nmatch \
		> Amel.filoci-vs-Am45.lastz &
lastz Am45.iloci.fa[multiple] Amel.ciloci.fa --allocate:traceback=500M \
		--ambiguous=iupac --filter=identity:95 --chain \
		--format=general:name1,length1,size1,name2,length2,size2,identity,nmatch \
		> Amel.ciloci-vs-Am45.lastz &
lastz Am45.iloci.fa[multiple] Amel.niloci.fa --allocate:traceback=500M \
		--ambiguous=iupac --filter=identity:95 --chain \
		--format=general:name1,length1,size1,name2,length2,size2,identity,nmatch \
		> Amel.niloci-vs-Am45.lastz &
lastz Am45.iloci.fa[multiple] Amel.iiloci.fa --allocate:traceback=500M \
		--ambiguous=iupac --filter=identity:95 --chain \
		--format=general:name1,length1,size1,name2,length2,size2,identity,nmatch \
		> Amel.iiloci-vs-Am45.lastz &
lastz Am45.iloci.fa[multiple] Amel.siloci.fa --allocate:traceback=500M \
		--ambiguous=iupac --filter=identity:95 --chain \
		--format=general:name1,length1,size1,name2,length2,size2,identity,nmatch \
		> Amel.siloci-vs-Am45.lastz &
wait
cd ..


#Processing the results:
#
iltypes=(filoci ciloci iiloci niloci siloci)

cd Amel
for iltype in ${iltypes[@]}
do
  n=`cat Amel.${iltype}.fa | egrep "^>" | wc -l`
  python3 ../scripts/process_lastz_output.py -n $n -i Amel.${iltype}-vs-Am45.lastz -v -o Amel.${iltype}-vs-Am45.summary
done
cd ..
```
