## AdditionalFile2 for "iLoci: Robust evaluation of genome content and organization for provisional and mature genome assemblies": runAtha.sh

```
#!/bin/bash
#
set -e
set -u
set -o pipefail

#1) Data set preparation:
#
mkdir Atha
labels=(Att6 Atha)

fidibus --numprocs=2 --refr=Att6,Atha download prep iloci breakdown stats

for species in ${labels[@]}
do
    ln -s ../species/${species}/${species}.iloci.tsv  Atha/ 
    ln -s ../species/${species}/${species}.iloci.fa   Atha/
    ln -s ../species/${species}/${species}.iloci.gff3 Atha/
done

cd ./Atha
makeblastdb -in Att6.iloci.fa -dbtype nucl -out Att6ILC -parse_seqids
makeblastdb -in Atha.iloci.fa -dbtype nucl -out AthaILC -parse_seqids
python3 ../scripts/split_iloci_by_type.py Att6
python3 ../scripts/split_iloci_by_type.py Atha
cd ..


#2) Running lastz:
#
cd ./Atha

# Assuming we have at least 5 processors, we run each lastz job in the background:
#
lastz Atha.iloci.fa[multiple] Att6.filoci.fa --allocate:traceback=500M \
		--ambiguous=iupac --filter=identity:95 --chain \
		--format=general:name1,length1,size1,name2,length2,size2,identity,nmatch \
		> Att6.filoci-vs-Atha.lastz &
lastz Atha.iloci.fa[multiple] Att6.ciloci.fa --allocate:traceback=500M \
		--ambiguous=iupac --filter=identity:95 --chain \
		--format=general:name1,length1,size1,name2,length2,size2,identity,nmatch \
		> Att6.ciloci-vs-Atha.lastz &
lastz Atha.iloci.fa[multiple] Att6.niloci.fa --allocate:traceback=500M \
		--ambiguous=iupac --filter=identity:95 --chain \
		--format=general:name1,length1,size1,name2,length2,size2,identity,nmatch \
		> Att6.niloci-vs-Atha.lastz &
lastz Atha.iloci.fa[multiple] Att6.iiloci.fa --allocate:traceback=500M \
		--ambiguous=iupac --filter=identity:95 --chain \
		--format=general:name1,length1,size1,name2,length2,size2,identity,nmatch \
		> Att6.iiloci-vs-Atha.lastz &
lastz Atha.iloci.fa[multiple] Att6.siloci.fa --allocate:traceback=500M \
		--ambiguous=iupac --filter=identity:95 --chain \
		--format=general:name1,length1,size1,name2,length2,size2,identity,nmatch \
		> Att6.siloci-vs-Atha.lastz &
wait
cd ..


#3) Processing the results:
#
iltypes=(filoci ciloci iiloci niloci siloci)

cd ./Atha
for iltype in ${iltypes[@]}
do
  n=`cat Att6.${iltype}.fa | egrep "^>" | wc -l`
  python3 ../scripts/process_lastz_output.py -n $n -i Att6.${iltype}-vs-Atha.lastz -v -o Att6.${iltype}-vs-Atha.summary
done
cd ..


###Reciprocal runs. Let's see whether we need them.===

cd ./Atha

# Assuming we have at least 5 processors, we run each lastz job in the background:
#
lastz Att6.iloci.fa[multiple] Atha.filoci.fa --allocate:traceback=500M \
		--ambiguous=iupac --filter=identity:95 --chain \
		--format=general:name1,length1,size1,name2,length2,size2,identity,nmatch \
		> Atha.filoci-vs-Att6.lastz &
lastz Att6.iloci.fa[multiple] Atha.ciloci.fa --allocate:traceback=500M \
		--ambiguous=iupac --filter=identity:95 --chain \
		--format=general:name1,length1,size1,name2,length2,size2,identity,nmatch \
		> Atha.ciloci-vs-Att6.lastz &
lastz Att6.iloci.fa[multiple] Atha.niloci.fa --allocate:traceback=500M \
		--ambiguous=iupac --filter=identity:95 --chain \
		--format=general:name1,length1,size1,name2,length2,size2,identity,nmatch \
		> Atha.niloci-vs-Att6.lastz &
lastz Att6.iloci.fa[multiple] Atha.iiloci.fa --allocate:traceback=500M \
		--ambiguous=iupac --filter=identity:95 --chain \
		--format=general:name1,length1,size1,name2,length2,size2,identity,nmatch \
		> Atha.iiloci-vs-Att6.lastz &
lastz Att6.iloci.fa[multiple] Atha.siloci.fa --allocate:traceback=500M \
		--ambiguous=iupac --filter=identity:95 --chain \
		--format=general:name1,length1,size1,name2,length2,size2,identity,nmatch \
		> Atha.siloci-vs-Att6.lastz &
wait
cd ..


#Processing the results:
#
iltypes=(filoci ciloci iiloci niloci siloci)

cd Atha
for iltype in ${iltypes[@]}
do
  n=`cat Atha.${iltype}.fa | egrep "^>" | wc -l`
  python3 ../scripts/process_lastz_output.py -n $n -i Atha.${iltype}-vs-Att6.lastz -v -o Atha.${iltype}-vs-Att6.summary
done
cd ..
```
