## AdditionalFile2 for "iLoci: Robust evaluation of genome content and organization for provisional and mature genome assemblies": filterAt11dna.py

```
from Bio import SeqIO

input_seq_iter = SeqIO.parse("ATgdna.fa", "fasta")

target_seq_iter = (record for record in input_seq_iter
                      if record.id.split()[0] not in ['ChrC','ChrM'])

SeqIO.write(target_seq_iter, "test.fa", "fasta")
```
