## AdditionalFile2 for "iLoci: Robust evaluation of genome content and organization for provisional and mature genome assemblies": gff3splitmultiparentlines.pl

```
#!/usr/bin/perl
#
#gff3splitmultiparentlines.pl
#
#	Usage: gff3splitmultiparentlines.pl input.gff3 > output.gff
#
#Purpose: Although a GFF3 feature may have mulitple parents annotated on the
#         the same line (comma-separated), some programs processing GFF3 files
#         don't parse this correctly and rather need separate lines for each
#         annotated feature/parent relationship. This script does that.
#         
#         GFF3 specification:
#         https://github.com/The-Sequence-Ontology/Specifications/blob/master/gff3.md

while (<>) {
	my ($parents) = $_ =~ /.*Parent=([^;]*[;]*).*/;
	chop $parents;
	my @prnt;
	if ($parents =~ /.*,.*/) {
		@prnt = split /,/, $parents;
		foreach my $p ( @prnt ) {
			my $newfeatureline = $_ =~ s/$parents/$p/r;
			print $newfeatureline;
		}
	}
	else {
		print $_;
	}
}
```
