## AdditionalFile2 for "iLoci: Robust evaluation of genome content and organization for provisional and mature genome assemblies": process_lastz_output.py

```
#!/usr/bin/env python
#
# process_lastz_output.py

import argparse
import sys
import pandas as pd
import json

pd.options.display.width = 0


def parse_args():
    parser = argparse.ArgumentParser()
    parser.add_argument('-i', '--input', type=str, help='lastz output file to read')
    parser.add_argument('-n', '--nquery', type=int, help='number of query iloci')
    parser.add_argument('-o', '--output', type=argparse.FileType('w'),
                        default=sys.stdout, help='Output file; default is '
                        'terminal (stdout)')
    parser.add_argument('-v', '--verbose',  action='store_true', help='verbose output')
    return parser.parse_args()

args = parse_args()

nbr_qloci = 0
if not args.input:
    print("\nPlease specify the input file with the -i option.\n\n", file=sys.stderr)
    exit()
if args.nquery:
    nbr_qloci = args.nquery
if args.output:
    sys.stdout = args.output


# lastztsv = LASTZ tsv output, read from file
#
lastztsv = pd.read_csv(args.input, sep='\t')

lastztsv.rename(columns = {'#name1' : 'name1'}, inplace = True)
# splitting the identity field to nbrid (number of identities) and
# nbral (number of aligned positions):
#
lastztsv[['nbrid','nbral']] = lastztsv['identity'].str.split('/',expand=True)
lastztsv[['nbrid','nbral']] = lastztsv[['nbrid','nbral']].apply(pd.to_numeric)

# qloci is the list of query loci in the LASTZ output:
#
qloci = list(sorted(set(lastztsv.name2)))
nbr_mapped = 0
nbr_unmapped =0
nbr_conserved = 0
nbr_unconserved = 0
nbr_redefined = 0
nbr_contained = 0
nbr_anchored = 0
conservediloci = {}
qsrelations = {}

for qlocus in qloci:
    if args.verbose:
        print("\n\n================================================================================")
        print("Working on query locus: ", qlocus)

    qhits = lastztsv[lastztsv.name2 == qlocus]
    if args.verbose: print(" with qhits:\n", qhits)
    qlength = qhits.iloc[0]['size2']

    # sloci is the list of subject loci hit by the query:
    #
    sloci = list(sorted(set(qhits['name1'])))
    candidatematches = {}
    conserved_as =[]
    contained_in =[]
    anchored_by =[]
    for slocus in sloci:
        if args.verbose: print("\nNow processing slocus", slocus)
        hsps = qhits[qhits.name1 == slocus]
        if args.verbose: print(" with hsps:\n", hsps)
        clength = hsps['nmatch'].sum()
        slength = qhits[qhits.name1 == slocus]['size1'].iloc[-1]
        if args.verbose:
            print("\n chain length cl= %d\tquery length ql= %d\t subject length sl= %d\tratios:\tcl/ql= %5.3f\tcl/sl= %5.3f"
              % (clength, qlength, slength, clength/qlength, clength/slength))
        if clength / qlength >= 0.90 or clength / slength >= 0.90:
            candidatematches[slocus] = clength
    if args.verbose: print("\nDone with processing sloci for query %s\n" % qlocus)

    #At this point we have either no candidate matches or candidate matches
    #with chain length at least 90% of either query or subject length.
    if len(candidatematches) == 0:
        if args.verbose: print("Status: %s could not be mapped by our criteria." % qlocus)
        nbr_unmapped += 1
        continue
    else:
        nbr_mapped += 1
        # let's see which criteria 1 hits also meet criterion 2
        for slocus,clength in candidatematches.items():
            slength = qhits[qhits.name1 == slocus]['size1'].iloc[-1]
            if clength >= qlength * 0.90  and  clength >= slength * 0.90:
               if args.verbose: print("%s is conserved as %s" % (qlocus, slocus))
               conserved_as.extend([slocus])
            elif clength >= qlength * 0.90:
               if args.verbose: print("%s is contained in %s" % (qlocus, slocus))
               contained_in.extend([slocus])
            else:
               if args.verbose: print("%s is anchored  by %s" % (qlocus, slocus))
               anchored_by.extend([slocus])
        if len(conserved_as) == 0:
            nbr_unconserved += 1
            if len(contained_in) > 0  and  len(anchored_by) > 0:
                if args.verbose: print("\nStatus: %s is unconserved-redefined" % (qlocus))
                nbr_redefined += 1
                qsrelations.update( {qlocus: ['redefined_to']} )
                qsrelations[qlocus].extend(contained_in)
                qsrelations[qlocus].extend(anchored_by)
            elif len(contained_in) > 0:
                if args.verbose: print("\nStatus: %s is unconserved-contained" % (qlocus))
                nbr_contained+= 1
                qsrelations.update( {qlocus: ['contained_in']} )
                qsrelations[qlocus].extend(contained_in)
            else:
                if args.verbose: print("\nStatus: %s is unconserved-anchored" % (qlocus))
                nbr_anchored += 1
                qsrelations.update( {qlocus: ['anchored_by']} )
                qsrelations[qlocus].extend(anchored_by)
            continue
        if len(conserved_as) == 1:
            conservediloci[qlocus] = conserved_as
            qsrelations.update( {qlocus: ['conserved']} )
            qsrelations[qlocus].extend(conserved_as)
            if args.verbose: print("\nStatus: %s is conserved as %s\n" % (qlocus,conservediloci[qlocus][0]))
        else:
            #We have multiple conserved targets. Is one best?
            trgtcl = {}
            trgtid = {}
            for target in conserved_as:
                hsps = qhits[qhits.name1 == target]
                #Tiebreaker one: maximal chain length; tiebreaker two: maximal nbrid/nbral ratio
                clength = hsps['nmatch'].sum()
                trgtcl[target] = clength
                trgtid[target] = hsps['nbrid'].sum() / hsps['nbral'].sum()
            tiebreakers = [key for m in [max(trgtcl.values())] for key,val in trgtcl.items() if val == m]
            if len(tiebreakers) > 1:
                tiebreakers = [key for m in [max(trgtid.values())] for key,val in trgtid.items() if val == m]
            conservediloci[qlocus] = tiebreakers
            if len(conservediloci[qlocus]) == 1:
                if args.verbose: print("\nStatus: %s is conserved as %s (and repetitive)\n" % (qlocus,conservediloci[qlocus][0]))
            else:
                if args.verbose: print("\nStatus: %s is multiply conserved as %s\n" % (qlocus,conservediloci[qlocus]))
            qsrelations.update( {qlocus: ['conserved']} )
            qsrelations[qlocus].extend(conservediloci[qlocus])
        nbr_conserved += 1


print("\n\n================================================================================")
print("Summary statistics:\n")
if nbr_qloci > 0:
    print("Number of qloci            :\t", nbr_qloci)
    print(" Number of qloci    w/o hits:\t", nbr_qloci - nbr_unmapped - nbr_mapped)

print(" Number of qloci    unmapped:\t", nbr_unmapped)
print(" Number of qloci      mapped:\t", nbr_mapped)
print("  Number of qloci   conserved:\t", nbr_conserved)
print("  Number of qloci unconserved:\t", nbr_unconserved)
print("   Number of qloci   contained:\t", nbr_contained)
print("   Number of qloci    anchored:\t", nbr_anchored)
print("   Number of qloci   redefined:\t", nbr_redefined)

counts = {}
counts['qloci_all']         = nbr_qloci
counts['qloci_wohits']      = nbr_qloci - nbr_unmapped - nbr_mapped
counts['qloci_unmapped']    = nbr_unmapped
counts['qloci_mapped']      = nbr_mapped
counts['qloci_unconserved'] = nbr_unconserved
counts['qloci_redefined']   = nbr_redefined
counts['qloci_contained']   = nbr_contained
counts['qloci_anchored']    = nbr_anchored
counts['qloci_conserved']   = nbr_conserved


if args.output:
    with open(args.output.name + '.counts', 'w') as f:
        f.write(json.dumps(counts))
        f.write("\n")
        f.close
    with open(args.output.name + '.qsrelations', 'w') as f:
        f.write(json.dumps(qsrelations))
        f.write("\n")
        f.close
###with open(args.output.name + '.counts', 'r') as f:
###    j = json.load(f)
###print(json.dumps(j, sort_keys=False, indent=4))
###with open(args.output.name + '.qsrelations', 'r') as f:
###    j = json.load(f)
###print(json.dumps(j, sort_keys=False, indent=4))
```
