## AdditionalFile2 for "iLoci: Robust evaluation of genome content and organization for provisional and mature genome assemblies": split_iloci_by_type.py

```
#!/usr/bin/env pythone
#
import sys
from Bio import SeqIO
import pandas as pd

if len(sys.argv) < 2:
    print("\nPlease provide the species label for input files\n",
          "<species>.iloci.tsv and <species>.iloci.fa.\n\n",
          file=sys.stderr)
    exit()

species = sys.argv[1]
iloci = pd.read_csv(species + '.iloci.tsv',sep='\t')
ii = iloci['LocusClass'] == "iiLocus"
iiloci = iloci[ii]
fi = iloci['LocusClass'] == "fiLocus"
filoci = iloci[fi]
siloci = iloci[iloci['LocusClass'] == "siLocus"]
niloci = iloci[iloci['LocusClass'] == "niLocus"]
ciloci = iloci[iloci['LocusClass'] == "ciLocus"]

ii_ids = set(iiloci['LocusId'])
fi_ids = set(filoci['LocusId'])
si_ids = set(siloci['LocusId'])
ni_ids = set(niloci['LocusId'])
ci_ids = set(ciloci['LocusId'])

input_seq_iterator = list(SeqIO.parse(species + ".iloci.fa", "fasta"))

ii_seq_iterator = (record for record in input_seq_iterator \
                      if record.id.split()[0] in ii_ids)

SeqIO.write(ii_seq_iterator, species + ".iiloci.fa", "fasta")

fi_seq_iterator = (record for record in input_seq_iterator \
                      if record.id.split()[0] in fi_ids)

SeqIO.write(fi_seq_iterator, species + ".filoci.fa", "fasta")

si_seq_iterator = (record for record in input_seq_iterator \
                      if record.id.split()[0] in si_ids)

SeqIO.write(si_seq_iterator, species + ".siloci.fa", "fasta")

ni_seq_iterator = (record for record in input_seq_iterator \
                      if record.id.split()[0] in ni_ids)

SeqIO.write(ni_seq_iterator, species + ".niloci.fa", "fasta")

ci_seq_iterator = (record for record in input_seq_iterator \
                      if record.id.split()[0] in ci_ids)

SeqIO.write(ci_seq_iterator, species + ".ciloci.fa", "fasta")
```
