## AdditionalFile2 for "iLoci: Robust evaluation of genome content and organization for provisional and mature genome assemblies": xgetDpul

#The following would be nice, but in reality, you will need to set up an
# account with JGI (https://signon.jgi.doe.gov/signon), open the links,
# choose to download the files, and then move the files into this directory ...
#
#curl -O http://genome.jgi.doe.gov/Dappu1/download/Daphnia_pulex.fasta.gz
#curl -O http://genome.jgi.doe.gov/Dappu1/download/FrozenGeneCatalog20110204.gff3.gz
#curl -O http://genome.jgi.doe.gov/Dappu1/download/FrozenGeneCatalog20110204.proteins.fasta.gz

#We need to simplify the protein identifiers to standard format to make potential fididbus
# sequence extraction work:
#
cp FrozenGeneCatalog20110204.proteins.fasta.gz FrozenGeneCatalog20110204.proteins.fasta.gzORIG
gunzip FrozenGeneCatalog20110204.proteins.fasta.gz
sed -i -e "s#>jgi|Dappu1|[^|]*|#>#" -e "s#|#-#g" -e "s#-\$##"  FrozenGeneCatalog20110204.proteins.fasta
gzip FrozenGeneCatalog20110204.proteins.fasta
