## AdditionalFile2 for "iLoci: Robust evaluation of genome content and organization for provisional and mature genome assemblies": Dpul.yml

```
Dpul:
    species: Daphnia pulex
    common: water flea
    source: local
    branch: invertebrate
    build: FrozenGeneCatalog20110204
    gdna: downloads/Daphnia_pulex.fasta.gz
    gff3: downloads/FrozenGeneCatalog20110204.gff3.gz
    prot: downloads/FrozenGeneCatalog20110204.proteins.fasta.gz
    checksums:
        gdna: 07772286ad596c9e4e295dcd4e11d51a7f9ee47d
        gff3: 1d443306d22805bb0bf92bd3b456722ee4eeec48
        prot: 53dd650185f313160cd89569a77d13bc231fa0f7
    description: from JGI; protein IDs modified
```
