## AdditionalFile2 for "iLoci: Robust evaluation of genome content and organization for provisional and mature genome assemblies": explore-iiLoci.ipynb

```
# %% [markdown]
# Exploring aggregate lengths of *r* consecutive iiLoci

Investigating different values of *r* to see if/when the distribution of consecutive iiLocus lengths (*r-scans*) approaches normal.
The first time you execute the notebook, you'll want to choose `Cell --> Run All`.
For subsequent executions, you'll want to edit the simple configuration cell and then while that cell is selected choose `Cell --> Run All Below`.

# %% [markdown]
## Supporting code

Code for accumulating adjacent iiLoci, as well as the obligatory boilerplate code to make figures that don't look horrible.
Only extremely curious tinkerers will want to tread here.

# %% [code]
from __future__ import print_function
%matplotlib inline

import glob
import matplotlib.pyplot as plt
import pandas
import seaborn
import numpy
from collections import deque
seaborn.set_context({"figure.figsize": (12, 8)})

# %% [code]
def len_consec_iiloci(data, r=2, filt=None):
    """
    Compute combined lengths for each set of r consecutive iiLoci.

    Iterate over iiLoci, accumulating sets of r consecutive iiLoci in a deque
    and yield the aggregate length of each set of r. Reset the deque when a new
    sequence is encountered.
    
    `data` is a pandas.DataFrame object, each row corresponding to a single
    iiLocus.

    `filt` is an (optional) maximum length threshold. If set, iiLoci whose
    length exceeds the threshold will be discarded.
    """
    iiloci = deque()
    for index, row in data.iterrows():
        if filt and row['Length'] > filt:
            continue
        if len(iiloci) > 0 and iiloci[0]['SeqID'] != row['SeqID']:
            iiloci = deque()
        iiloci.append(row)
        if len(iiloci) == r:
            aggregate_length = sum([ilocus['Length'] for ilocus in iiloci])
            yield aggregate_length
            iiloci.popleft()

# %% [code]
def make_hist(data, log10=False, bins=40, xlab=None, ylab=None,
              xlim=None, ylim=None, title=None, color='grey'):
    if xlim:
        data = [dat for dat in data if dat < xlim[1]]
    if log10:
        _ = plt.hist(numpy.log10(data), bins=bins, color=color)
    else:
        _ = plt.hist(data, bins=bins, color=color)
    seaborn.despine(left=True)
    if xlab is not None:
        _ = plt.xlabel(xlab, fontsize=18)
    if ylab is not None:
        _ = plt.ylabel(ylab, fontsize=18)
    if xlim is not None:
        _ = plt.xlim(xlim[0], xlim[1])
    if ylim is not None:
        _ = plt.ylim(ylim[0], ylim[1])
    if title:
        plt.title(title, fontsize=20)
    plt.show()

# %% [markdown]
## Configuration

Okay, here is where it gets interesting.
Specify what you want to look at with the configuration options below.

# %% [code]
# Species (choices: Scer Cele Crei Mtru Agam Dmel Xtro Drer Mmus Hsap Pdom Dpul Vcar)
species = 'Mmus'

# Range of r values to consider
rrange = [2, 30, 60, 120]

# The next line specifies the quantile at which outliers in the long tail of the
# length distribution will be discarded. Set it to None to disable the filter.
maxlenqnt = 0.95
# maxlenqnt = None


# The next line controls the x-axis range of viewing window, which can make a 
# difference (especially when the filter is disabled) in the histogram's
# resolution.
xrange = None
# xrange = [0, 100000]

# %% [markdown]
You shouldn't have to touch anything below this line.
Once you've set the configuration values, go ahead and select that cell and then click `Cell --> Run All Below` in the menu above.

# %% [code]
filename = '../data/{}/{}.iloci.tsv'.format(species, species)
iloci = pandas.read_csv(filename, sep='\t')
iiloci = iloci.loc[iloci.LocusClass == 'iiLocus']

maxlen = 0
if maxlenqnt:
    maxlen = int(iiloci['Length'].quantile(maxlenqnt))
    print('Maximal iiLocus length: {} bp'.format(maxlen))

# %% [code]
iiloci.head()

# %% [code]
for k in rrange:
    title = '{} consecutive iiLoci ({})'.format(k, species)
    agglens = [x for x in len_consec_iiloci(iiloci, r=k, filt=maxlen)]
    make_hist(agglens, xlab='Aggregate length (bp)', ylab='Frequency',
              title=title, xlim=xrange)

# %% [code]
species_list = ['Cele','Mtru']
rrange = [2, 5, 8]
for species in species_list:
    filename = '../data/{}/{}.iloci.tsv'.format(species, species)
    iloci = pandas.read_csv(filename, sep='\t')
    iiloci = iloci.loc[iloci.LocusClass == 'iiLocus']

    maxlen = 0
    if maxlenqnt:
        maxlen = int(iiloci['Length'].quantile(maxlenqnt))
        print('Maximal iiLocus length: {} bp'.format(maxlen))
    for k in rrange:
        title = '{} consecutive iiLoci ({})'.format(k, species)
        agglens = [x for x in len_consec_iiloci(iiloci, r=k, filt=maxlen)]
        make_hist(agglens, xlab='Aggregate length (bp)', ylab='Frequency',
                  title=title, xlim=xrange)

# %% [code]
```
