## AdditionalFile2 for "iLoci: Robust evaluation of genome content and organization for provisional and mature genome assemblies": explore-lengthANDprcntGC.ipynb

```
# %% [code]
%matplotlib inline

import numpy
import matplotlib
import matplotlib.pyplot as plt
import pandas
import seaborn
seaborn.set_context({"figure.figsize": (16, 8)})
matplotlib.rcParams['axes.labelsize'] = 16
matplotlib.rcParams['xtick.labelsize'] = 12
matplotlib.rcParams['ytick.labelsize'] = 12

# %% [code]
def do_plots(species, label, root="../data", lengthticks=[3, 4, 5, 6],
             lengthticklabs=['1,000', '10,000', '100,000', '1,000,000'],
             lengthbins=numpy.linspace(2.75, 6.0, 40),
             gcbins=numpy.linspace(20, 80, 40)):
    ilocusdata = pandas.read_table('{}/{}/{}.iloci.tsv'.format(root, species, species))
    milocusdata = pandas.read_table('{}/{}/{}.miloci.tsv'.format(root, species, species))
    ilocusdata['LogLength'] = numpy.log10(ilocusdata['Length'])
    milocusdata['LogLength'] = numpy.log10(milocusdata['Length'])
    siloci = ilocusdata.loc[(ilocusdata.LocusClass == 'siLocus')]
    miloci = milocusdata.loc[(milocusdata.LocusClass == 'miLocus')]
    singleton_siloci = milocusdata.loc[(milocusdata.LocusClass == 'siLocus')]
    singleton_locuspos = [lp[6:-1] for lp in singleton_siloci['LocusPos']]
    milocus_siloci = siloci.loc[~(siloci.LocusPos.isin(singleton_locuspos))]
    fig, axes = plt.subplots(1, 2)
    ax1, ax2 = axes
    _ = ax1.hist(singleton_siloci['LogLength'], bins=lengthbins, alpha=0.5)
    _ = ax1.hist(milocus_siloci['LogLength'], bins=lengthbins, alpha=0.5)
    _ = ax1.set_title('siLocus Length ({})'.format(label))
    _ = ax1.set_xlabel('Length (bp)')
    _ = ax1.set_ylabel('Frequency')
    _ = ax1.legend(loc='upper right', labels=('Unmerged', 'Merged'))
    if lengthticks:
        _ = ax1.set_xticks(lengthticks)
    if lengthticklabs:
        _ = ax1.set_xticklabels(lengthticklabs)
    _ = ax2.hist(singleton_siloci['GCContent'] * 100, bins=gcbins, alpha=0.5)
    _ = ax2.hist(milocus_siloci['GCContent'] * 100, bins=gcbins, alpha=0.5)
    _ = ax2.set_title('siLocus Nucleotide Composition ({})'.format(label))
    _ = ax2.set_xlabel('% GC Content')
    _ = ax2.set_ylabel('Frequency')
    _ = ax2.legend(loc='upper right', labels=('Unmerged', 'Merged'))

# %% [code]
do_plots('Cele', 'C. elegans', "../data", lengthticks=[3, 4, 5], lengthticklabs=['1,000', '10,000', '100,000'],
         lengthbins=numpy.linspace(2.5, 5.5, 40), gcbins=numpy.linspace(20, 55, 40))

# %% [code]
do_plots('Crei', 'Chlamydomonas', '../data', lengthticks=[3, 4, 5], lengthticklabs=['1,000', '10,000', '100,000'],
         lengthbins=numpy.linspace(2.5, 5.5, 40), gcbins=numpy.linspace(50, 80, 40))

# %% [code]
do_plots('Mtru', 'Medicago', lengthticks=[3, 4, 5], lengthticklabs=['1,000', '10,000', '100,000'],
         lengthbins=numpy.linspace(2.5, 5.5, 40), gcbins=numpy.linspace(20, 45, 40))

# %% [code]
do_plots('Agam', 'Anopheles gambiae', gcbins=numpy.linspace(25, 65, 40))

# %% [code]
do_plots('Dmel', 'Drosphila', gcbins=numpy.linspace(25, 65, 40))

# %% [code]
do_plots('Xtro', 'Xenopus', lengthbins=numpy.linspace(2.5, 6.5, 40), gcbins=numpy.linspace(30, 60, 40))

# %% [code]
do_plots('Drer', 'Zebrafish', lengthbins=numpy.linspace(2.5, 6.5, 40), gcbins=numpy.linspace(25, 55, 40))

# %% [code]
do_plots('Mmus', 'Mouse', lengthbins=numpy.linspace(2.5, 6.5, 40), gcbins=numpy.linspace(30, 70, 40))

# %% [code]
do_plots('Hsap', 'Human', lengthbins=numpy.linspace(2.5, 6.5, 40), gcbins=numpy.linspace(30, 75, 40))

# %% [code]
root = '../data'
species = 'Mmus'
ilocusdata = pandas.read_table('{}/{}/{}.iloci.tsv'.format(root,species, species))
milocusdata = pandas.read_table('{}/{}/{}.miloci.tsv'.format(root,species, species))
ilocusdata['LogLength'] = numpy.log10(ilocusdata['Length'])
milocusdata['LogLength'] = numpy.log10(milocusdata['Length'])
siloci = ilocusdata.loc[(ilocusdata.LocusClass == 'siLocus')]
miloci = milocusdata.loc[(milocusdata.LocusClass == 'miLocus')]
singleton_siloci = milocusdata.loc[(milocusdata.LocusClass == 'siLocus')]
peak = singleton_siloci.loc[(singleton_siloci.Length > 1800) & (singleton_siloci.Length < 2200)]
_ = plt.hist(peak['Length'], bins=40)

# %% [code]
ioi = siloci.loc[(siloci.Length > 1920) & (siloci.Length < 1980)]
ioi.sample(n=5)

# %% [code]
ioi.sample(n=5)

# %% [code]
root = '../data'
species = 'Hsap'
ilocusdata = pandas.read_table('{}/{}/{}.iloci.tsv'.format(root,species, species))
milocusdata = pandas.read_table('{}/{}/{}.miloci.tsv'.format(root,species, species))
ilocusdata['LogLength'] = numpy.log10(ilocusdata['Length'])
milocusdata['LogLength'] = numpy.log10(milocusdata['Length'])
siloci = ilocusdata.loc[(ilocusdata.LocusClass == 'siLocus')]
miloci = milocusdata.loc[(milocusdata.LocusClass == 'miLocus')]
singleton_siloci = milocusdata.loc[(milocusdata.LocusClass == 'siLocus')]
peak = singleton_siloci.loc[(singleton_siloci.Length > 1800) & (singleton_siloci.Length < 2200)]
_ = plt.hist(peak['Length'], bins=40)

# %% [code]
ioi = siloci.loc[(siloci.Length > 1920) & (siloci.Length < 1980)]
ioi.sample(n=5)

# %% [code]
ioi.sample(n=5)

# %% [code]
```
