## AdditionalFile2 for "iLoci: Robust evaluation of genome content and organization for provisional and mature genome assemblies": explore-miLoci.ipynb

```
# %% [code]
from __future__ import division
%matplotlib inline

import matplotlib
import matplotlib.pyplot as plt
import pandas
import re
import seaborn

# %% [code]
def parse_seqreg(entry):
    seqregmatch = re.search('##sequence-region\s+(\S+)\s+(\d+)\s+(\d+)', entry)
    assert seqregmatch
    seqid = seqregmatch.group(1)
    start = int(seqregmatch.group(2))
    assert start == 1
    end = int(seqregmatch.group(3))
    return seqid, start, end

# %% [code]
def gff3_to_hist(species, seqid, root='../data'):
    seqlength = 0
    filename = '{}/{}/{}.iloci.gff3'.format(root, species, species)
    with open(filename, 'r') as infile:
        for line in infile:
            if line.startswith('##sequence-region'):
                seq, start, end = parse_seqreg(line)
                if seq == seqid:
                    seqlength = end
    
    milocus_starts = list()
    filename = '{}/{}/{}.miloci.gff3'.format(root, species, species)
    with open(filename, 'r') as infile:
        for line in infile:
            if 'iLocus_type=miLocus' not in line:
                continue
            fields = line.split('\t')
            assert len(fields) == 9
            seq = fields[0]
            if seq != seqid:
                continue
            start = int(fields[3]) / seqlength
            milocus_starts.append(start)
    return milocus_starts

# %% [code]
seaborn.set_context({"figure.figsize": (12, 6)})
chromosomes = 'NC_000067.6 NC_000068.7 NC_000069.6 NC_000070.6 NC_000071.6 NC_000072.6 NC_000073.6 NC_000074.6 NC_000075.6 NC_000076.6 NC_000077.6 NC_000078.6 NC_000079.6 NC_000080.6 NC_000081.6 NC_000082.6 NC_000083.6 NC_000084.6 NC_000085.6 NC_000086.7 NC_000087.7'
for chrom in chromosomes.split(' '):
    mipos = gff3_to_hist('Mmus', chrom)
    _ = plt.hist(mipos, bins=75)
    _ = plt.title(chrom)
    _ = plt.show()

# %% [code]
seaborn.set_context({"figure.figsize": (12, 6)})
chromosomes = 'NC_004353.4 NC_004354.4 NC_024512.1 NT_033777.3 NT_033778.4 NT_033779.5 NT_037436.4'
for chrom in chromosomes.split(' '):
    mipos = gff3_to_hist('Dmel', chrom)
    _ = plt.hist(mipos, bins=100)
    _ = plt.title(chrom)
    _ = plt.show()

# %% [code]
seaborn.set_context({"figure.figsize": (12, 6)})
chromosomes = 'NC_003279.8 NC_003280.10 NC_003281.10 NC_003282.8 NC_003283.11 NC_003284.9'
for chrom in chromosomes.split(' '):
    mipos = gff3_to_hist('Cele', chrom)
    _ = plt.hist(mipos, bins=100)
    _ = plt.title(chrom)
    _ = plt.show()

# %% [code]
seaborn.set_context({"figure.figsize": (12, 6)})
chromosomes = 'NC_053042.1 NC_053043.1 NC_053044.1 NC_053045.1 NC_053046.1 NC_053047.1 NC_053048.1 NC_053049.1'
for chrom in chromosomes.split(' '):
    mipos = gff3_to_hist('Mtru', chrom)
    _ = plt.hist(mipos, bins=100)
    _ = plt.title(chrom)
    _ = plt.show()

# %% [code]
seaborn.set_context({"figure.figsize": (12, 6)})
chromosomes = 'NC_004818.2 NT_078265.2 NT_078266.2 NT_078267.5 NT_078268.4'
for chrom in chromosomes.split(' '):
    mipos = gff3_to_hist('Agam', chrom)
    _ = plt.hist(mipos, bins=100)
    _ = plt.title(chrom)
    _ = plt.show()

# %% [code]
seaborn.set_context({"figure.figsize": (12, 6)})
chromosomes = 'NC_007112.6 NC_007113.6 NC_007114.6 NC_007115.6 NC_007116.6 NC_007117.6 NC_007118.6 NC_007119.6 NC_007120.6 NC_007121.6 NC_007122.6 NC_007123.6 NC_007124.6 NC_007125.6 NC_007126.6 NC_007127.6 NC_007128.6 NC_007129.6 NC_007130.6 NC_007131.6 NC_007132.6 NC_007133.6 NC_007134.6 NC_007135.6 NC_007136.6'
for chrom in chromosomes.split(' '):
    mipos = gff3_to_hist('Drer', chrom)
    _ = plt.hist(mipos, bins=100)
    _ = plt.title(chrom)
    _ = plt.show()

# %% [code]
seaborn.set_context({"figure.figsize": (12, 6)})
chromosomes = 'NC_000001.11 NC_000002.12 NC_000003.12 NC_000004.12 NC_000005.10 NC_000006.12 NC_000007.14 NC_000008.11 NC_000009.12 NC_000010.11 NC_000011.10 NC_000012.12 NC_000013.11 NC_000014.9 NC_000015.10 NC_000016.10 NC_000017.11 NC_000018.10 NC_000019.10 NC_000020.11 NC_000021.9 NC_000022.11 NC_000023.11 NC_000024.10'
for chrom in chromosomes.split(' '):
    mipos = gff3_to_hist('Hsap', chrom)
    _ = plt.hist(mipos, bins=75)
    _ = plt.title(chrom)
    _ = plt.show()

# %% [code]
```
