## AdditionalFile2 for "iLoci: Robust evaluation of genome content and organization for provisional and mature genome assemblies": make-F3a-SF6-SF7.ipynb

```
# %% [code]
%matplotlib inline

import math
import matplotlib
import matplotlib.pyplot as plt
import matplotlib.lines as ml
import pandas
import random
import seaborn

matplotlib.rcParams['axes.labelsize'] = 16
matplotlib.rcParams['xtick.labelsize'] = 12
matplotlib.rcParams['ytick.labelsize'] = 12
matplotlib.rcParams['savefig.dpi'] = 300
seaborn.set_context({"figure.figsize": (15, 10)})

# %% [code]
def calc_rho(species, basedir='../data'):
    """
    Calculate \rho, the average fraction of iiLocus length to giLocus length,
    for a particular genome.
    """
    files = ['{}/{}/{}.iloci.tsv'.format(basedir, species, species)]
    data = pandas.concat([pandas.read_table(f) for f in files])
    giloci = data.loc[data.LocusClass.isin(['siLocus', 'ciLocus', 'niLocus'])]
    iiloci = data.loc[data.LocusClass.isin(['iiLocus'])]
    quant = list(giloci['Length'].quantile([0.05, 0.95]))
    giloci_q = giloci.loc[(giloci.Length >= quant[0]) & (giloci.Length <= quant[1])]
    quant = list(iiloci['Length'].quantile([0.05, 0.95]))
    iiloci_q = iiloci.loc[(iiloci.Length >= quant[0]) & (iiloci.Length <= quant[1])]
    rho = iiloci_q['Length'].mean() / giloci_q['Length'].mean()
    return rho

# %% [markdown]
## Genome compactness in theory

If the average iiLocus length is ρ-times the average giLocus length, then the relationship between φ and σ is given by $\sigma = \frac{\phi}{1 + 1(1-\phi)\rho}$.
Below we show (φ, σ) curves with ρ equal to 0.1, 1, 2, 4, and 8.

# %% [code]
x = [i / 1000 for i in range(1000)]
colors = seaborn.color_palette("hls", 5)
plots = list()
rhos = [0.1, 1, 2, 4, 8]
labels = ['ρ = {}'.format(num) for num in rhos]
for j, rho in enumerate(rhos):
    y = [(i/(1+(1-i)*rho)) for i in x]
    theplot = ml.Line2D(x, y, color=colors[j], label=labels[j])
    _ = plt.plot(x, y, color=colors[j], label=labels[j])
    plots.append(theplot)
plt.xlabel('φ (Proportion of giLoci merged into miLoci)')
plt.ylabel('σ (Proportion of sequence occupied by miLoci)')
plt.legend(plots, labels,
               loc='upper left', ncol=1, fontsize=14)
plt.show()

# %% [markdown]
## Empirical genome compactness

Now let's look at empirical (φ, σ) values calculated on chromosome/scaffold sequences from 10 reference genomes.

# %% [code]
data = pandas.read_csv('../tables/phisigma-refr.tsv',sep='\t')
speciesnames = ['Saccharomyces cerevisiae', 'Caenorhabditis elegans',
                'Chlamydomonas reinhardtii', 'Medicago truncatula',
                'Drosophila melanogaster', 'Anopheles gambiae',
                'Xenopus tropicalis', 'Danio rerio',
                'Mus musculus', 'Homo sapiens']
specieslist = ['Scer', 'Cele', 'Crei', 'Mtru', 'Dmel',
               'Agam', 'Xtro', 'Drer', 'Mmus', 'Hsap']

plots = list()
palette = seaborn.color_palette("hls", len(specieslist))
for i, species in enumerate(specieslist):
    specdata = data.loc[(data.Species == species)]
    plot = plt.scatter(specdata['Phi'], specdata['Sigma'], color=palette[i])
    plots.append(plot)
_ = plt.xlim(-0.02, 1.02)
_ = plt.ylim(-0.02, 1.02)
_ = plt.xlabel('φ (Proportion of giLoci merged into miLoci)')
_ = plt.ylabel('σ (Proportion of sequence occupied by miLoci)')
_ = plt.legend(plots, speciesnames, scatterpoints=1,
           loc='upper left',
           ncol=2,
           fontsize=12)
_ = plt.show()

# %% [code]
# Calculating rho:
#
for s in specieslist:
    print(s, calc_rho(s))

# %% [markdown]
## Now let's put it together:

# %% [code]
data = pandas.read_csv('../tables/phisigma-refr.tsv',sep='\t')
speciesnames = ['Saccharomyces cerevisiae', 'Caenorhabditis elegans',
                'Chlamydomonas reinhardtii', 'Medicago truncatula',
                'Drosophila melanogaster', 'Anopheles gambiae',
                'Xenopus tropicalis', 'Danio rerio',
                'Mus musculus', 'Homo sapiens']
specieslist = ['Scer', 'Cele', 'Crei', 'Mtru', 'Dmel',
               'Agam', 'Xtro', 'Drer', 'Mmus', 'Hsap']

rhos = [0.1, 1, 2, 4, 8]
for j, rho in enumerate(rhos):
    y = [(i/(1+(1-i)*rho)) for i in x]
    _ = plt.plot(x, y, color='#cc6666', alpha=0.2)

plots = list()
palette = seaborn.color_palette("hls", len(specieslist))
for i, species in enumerate(specieslist):
    specdata = data.loc[(data.Species == species)]
    plot = plt.scatter(specdata['Phi'], specdata['Sigma'], color=palette[i])
    plots.append(plot)

_ = plt.xlim(-0.02, 1.02)
_ = plt.ylim(-0.02, 1.02)
_ = plt.xlabel('φ (Proportion of giLoci merged into miLoci)')
_ = plt.ylabel('σ (Proportion of sequence occupied by miLoci)')
_ = plt.legend(plots, speciesnames, scatterpoints=1,
           loc='upper left',
           ncol=2,
           fontsize=12)
#_= plt.show()
plt.savefig('../figures/SLB21-Fig3a.png')
plt.show()

# %% [markdown]
## Different values of δ

To evaluate the robustness of the (φ, σ) measures with respect to the δ (delta) parameter, we recomputed iLoci at δ=300 and δ=750 for comparison with the default δ=500.
Again, we show centroids in favor of the full scatter plot to facilitate comparison.
The circles, boxes, and stars correspond to δ=300, δ=500, and δ=750, respectively.

# %% [code]
data500 = pandas.read_csv('../tables/phisigma-refr-centroids-delta500.tsv',sep='\t')
data300 = pandas.read_csv('../tables/phisigma-refr-centroids-delta300.tsv',sep='\t')
data750 = pandas.read_csv('../tables/phisigma-refr-centroids-delta750.tsv',sep='\t')
speciesnames = ['Saccharomyces cerevisiae', 'Caenorhabditis elegans',
                'Chlamydomonas reinhardtii', 'Medicago truncatula',
                'Drosophila melanogaster', 'Anopheles gambiae',
                'Xenopus tropicalis', 'Danio rerio',
                'Mus musculus', 'Homo sapiens']
specieslist = ['Scer', 'Cele', 'Crei', 'Mtru', 'Dmel',
               'Agam', 'Xtro', 'Drer', 'Mmus', 'Hsap']

plots = list()
palette = seaborn.color_palette("hls", len(specieslist))
for i, species in enumerate(specieslist):
    phis = list()
    sigmas = list()

    specdata = data300.loc[(data300.Species == species)]
    phis.append(list(specdata['Phi'])[0])
    sigmas.append(list(specdata['Sigma'])[0])
    plot = plt.scatter(specdata['Phi'], specdata['Sigma'], color=palette[i], s=128, alpha=0.25)
    plot = plt.scatter(specdata['Phi'], specdata['Sigma'], color=palette[i], s=128, facecolor='none', lw=2)
            
    specdata = data500.loc[(data500.Species == species)]
    phis.append(list(specdata['Phi'])[0])
    sigmas.append(list(specdata['Sigma'])[0])
    plot = plt.scatter(specdata['Phi'], specdata['Sigma'], color=palette[i], s=128, marker='s')
    # ... use the delta500 marker for the legend:
    plots.append(plot)

    specdata = data750.loc[(data750.Species == species)]
    phis.append(list(specdata['Phi'])[0])
    sigmas.append(list(specdata['Sigma'])[0])
    plot = plt.scatter(specdata['Phi'], specdata['Sigma'], color=palette[i], s=128, marker='*', lw=2)
    plot = plt.plot(phis, sigmas, color=palette[i], alpha=0.5, ls=':')
    
_ = plt.xlim(-0.02, 1.02)
_ = plt.ylim(-0.02, 1.02)
_ = plt.xlabel('φ (Proportion of giLoci merged into miLoci)')
_ = plt.ylabel('σ (Proportion of sequence occupied by miLoci)')
_ = plt.legend(plots, speciesnames, scatterpoints=1,
           loc='upper left',
           ncol=2,
           fontsize=12)
plt.savefig('../figures/SLB21-SuppFig6.png')

# %% [markdown]
## Comparison of real with random gene arrangements

# %% [code]
data = pandas.read_csv('../tables/phisigma-refr-centroids-delta500.tsv',sep='\t')
sdata = pandas.read_csv('../tables/phisigma-refr-shuffled-centroids-delta500.tsv',sep='\t')
speciesnames = ['Saccharomyces cerevisiae', 'Caenorhabditis elegans',
                'Chlamydomonas reinhardtii', 'Medicago truncatula',
                'Drosophila melanogaster', 'Anopheles gambiae',
                'Xenopus tropicalis', 'Danio rerio',
                'Mus musculus', 'Homo sapiens']
specieslist = ['Scer', 'Cele', 'Crei', 'Mtru', 'Dmel',
               'Agam', 'Xtro', 'Drer', 'Mmus', 'Hsap']

plots = list()
palette = seaborn.color_palette("hls", len(specieslist))
for i, species in enumerate(specieslist):
    phis = list()
    sigmas = list()
        
    specdata = data.loc[(data.Species == species)]
    phis.append(list(specdata['Phi'])[0])
    sigmas.append(list(specdata['Sigma'])[0])
    plot = plt.scatter(specdata['Phi'], specdata['Sigma'], color=palette[i], marker='s', s=64)
    plots.append(plot)

    specdata = sdata.loc[(sdata.Species == species)]
    phis.append(list(specdata['Phi'])[0])
    sigmas.append(list(specdata['Sigma'])[0])
    plot = plt.scatter(specdata['Phi'], specdata['Sigma'], color=palette[i], s=128, alpha=0.25)
    plot = plt.scatter(specdata['Phi'], specdata['Sigma'], color=palette[i], s=128, facecolor='none', lw=2)
    #plots.append(plot)
    plot = plt.plot(phis, sigmas, color=palette[i], alpha=0.5, ls=':')
    
_ = plt.xlim(-0.02, 1.02)
_ = plt.ylim(-0.02, 1.02)
_ = plt.xlabel('φ (Proportion of giLoci merged into miLoci)')
_ = plt.ylabel('σ (Proportion of sequence occupied by miLoci)')
_ = plt.legend(plots, speciesnames, scatterpoints=1,
           loc='upper left',
           ncol=2,
           fontsize=12)
plt.savefig('../figures/SLB21-SuppFig7.png')

# %% [code]
```
