## AdditionalFile2 for "iLoci: Robust evaluation of genome content and organization for provisional and mature genome assemblies": make-F3b.ipynb

```
# %% [code]
%matplotlib inline

import math
import matplotlib
import matplotlib.pyplot as plt
import matplotlib.lines as ml
import pandas as pd
import seaborn

matplotlib.rcParams['axes.labelsize'] = 16
matplotlib.rcParams['xtick.labelsize'] = 12
matplotlib.rcParams['ytick.labelsize'] = 12
matplotlib.rcParams['savefig.dpi'] = 300
seaborn.set_context({"figure.figsize": (15, 10)})

# %% [code]
file_names = ['fungi','invertebrate','vertebrate_mammalian','vertebrate_other','plant','protozoa']
branches = ['Fungi', 'Invertebrates', 'Mammalian vertebrates', 'Other vertebrates', 'Plants', 'Protozoa']

x = [i / 1000 for i in range(1000)]
rhos = [0.1, 1, 2, 4, 8]
for j, rho in enumerate(rhos):
    y = [(i/(1+(1-i)*rho)) for i in x]
    _ = plt.plot(x, y, color='#cc6666', alpha=0.2)

plots = []
palette = seaborn.color_palette("hls", len(branches))
i = 0
for file in file_names:
    specdata = pd.read_csv('../taxa/' + file + '_centroids.tsv', sep = '\t')
    plot = plt.scatter(specdata['Phi'], specdata['Sigma'], color=palette[i])
    plots.append(plot)
    i += 1
    
_ = plt.xlim(-0.02, 1.02)
_ = plt.ylim(-0.02, 1.02)
_ = plt.xlabel('φ (Proportion of giLoci merged into miLoci)')
_ = plt.ylabel('σ (Proportion of sequence occupied by miLoci)')
_ = plt.legend(plots, branches, scatterpoints=1,
           loc='upper left',
           ncol=2,
           fontsize=12)
plotfile = "../figures/SLB21-Fig3b.png"
plt.savefig(plotfile)
_ = plt.show()

# %% [code]
```
