## AdditionalFile2 for "iLoci: Robust evaluation of genome content and organization for provisional and mature genome assemblies": make-F4a-F4b-SF5a.ipynb

```
# %% [code]
%matplotlib inline

import glob
import math
import matplotlib
import matplotlib.pyplot as plt
import matplotlib.lines as ml
import numpy
import pandas
import random
import seaborn

matplotlib.rcParams['axes.labelsize'] = 16
matplotlib.rcParams['xtick.labelsize'] = 12
matplotlib.rcParams['ytick.labelsize'] = 12
matplotlib.rcParams['savefig.dpi'] = 300
seaborn.set_context({"figure.figsize": (12, 8)})

# %% [code]
def calc_rho(species, basedir='../data'):
    """
    Calculate \rho, the proportion of iiLocus length to giLocus length, for a particular genome.
    """
    files = ['{}/{}/{}.iloci.tsv'.format(basedir, s, s) for s in species]
    data = pandas.concat([pandas.read_csv(f, sep='\t') for f in files])
    giloci = data.loc[data.LocusClass.isin(['siLocus', 'ciLocus', 'niLocus'])]
    iiloci = data.loc[data.LocusClass.isin(['iiLocus'])]
    quant = list(giloci['Length'].quantile([0.05, 0.95]))
    giloci_q = giloci.loc[(giloci.Length >= quant[0]) & (giloci.Length <= quant[1])]
    quant = list(iiloci['Length'].quantile([0.05, 0.95]))
    iiloci_q = iiloci.loc[(iiloci.Length >= quant[0]) & (iiloci.Length <= quant[1])]
    rho = iiloci_q['Length'].mean() / giloci_q['Length'].mean()
    return rho

# %% [code]
colors = ['#a6cee3', '#1f78b4', '#b2df8a', '#33a02c', '#cc33cc',
          '#fb9a99', '#e31a1c', '#fdbf6f']

def load_and_prep(filename, specieslist, index='Species'):
    """Boilerplate code: load data into memory"""
    dataframe = pandas.read_csv(filename,sep='\t')
    df = dataframe[dataframe[index].isin(specieslist)]
    df = df.set_index([index]).reindex(specieslist)
    return df

def make_barplot(dataframe, xlab=None, ylab=None, ylim=None, lloc=2,
                 legend=True, yticks=None, yticklabs=None):
    """Biolerplate code: make a pretty barplot"""

    plot = dataframe.plot(kind='bar', stacked=True, figsize=(15, 10),
                          fontsize=16, rot=0, color=colors)
    seaborn.despine(left=True)
    if legend is True:
        _ = plot.legend(loc=lloc, prop={'size': 16})
    if xlab is not None:
        _ = plt.xlabel(xlab)
    if ylab is not None:
        _ = plt.ylabel(ylab)
    if ylim is not None:
        _ = plot.set_ylim(ylim[0], ylim[1])
    if yticks:
        _ = plot.set_yticks(yticks)
    if yticklabs:
        _ = plot.set_yticklabels(yticklabs)
    _ = plot.yaxis.label.set_fontsize(18)
    

# %% [code]
speciesnames = ['Volvox carteri', 'Chlamydomonas reinhardtii',
                'Chlorella variabilis', 'Coccomyxa subellipsoidea', 'Micromonas commoda',
                'Ostreococcus lucimarinus']
specieslist = ['Vcar', 'Crei', 'Cvar', 'Csub', 'Mcom', 'Oluc']
palette = seaborn.color_palette("hls", len(speciesnames))
# This is using an outdated directory structure. 
#rho = calc_rho(specieslist, basedir='scratch/chlorophyta/species')
rho = calc_rho(specieslist)

data = pandas.read_csv('../tables/phisigma-Vcar.tsv',sep='\t')
plots = list()
for i, species in enumerate(specieslist):
    # Uncomment and edit the next two lines if you want to exclude/restrict the plot
    # if species in ['Vcar']:
    #    continue
    specdata = data.loc[(data.Species == species)]
    plot = plt.scatter(specdata['Phi'], specdata['Sigma'], color=palette[i])
    plots.append(plot)
    x = [k / 1000 for k in range(1000)]
    y = [(k/(1+(1-k)*rho)) for k in x]
    _ = plt.plot(x, y, color='#cccccc')
_ = plt.xlim(-0.02, 1.02)
_ = plt.ylim(-0.02, 1.02)
_ = plt.xlabel('φ (Proportion of giLoci merged into miLoci)')
_ = plt.ylabel('σ (Proportion of sequence occupied by miLoci)')
_ = plt.legend(plots, speciesnames, scatterpoints=1,
           loc='upper left',
           ncol=2,
           fontsize=12)
plt.savefig('../figures/SLB21-SuppFig5a.png')
plt.show()

# %% [code]
specieslist = ['Apro','Cvar','Csub','Mcom',
               'Mpus','Oluc','Otau','Crei','Vcar']
bd_bp = load_and_prep('../Chlorophyta/Chlorophyta-breakdown-bp.tsv', specieslist)
bd_counts = load_and_prep('../Chlorophyta/Chlorophyta-breakdown-counts.tsv', specieslist)
make_barplot(bd_counts, xlab='', ylim=(0, 2.5e4),
             ylab='Total iLocus count')
plt.savefig('../figures/SLB21-Fig4a.png')
make_barplot(bd_bp, xlab='', ylim=(0, 1.5e8),
             yticks=[2.5e7 * x for x in [0,1,2,3,4,5,6]],
             yticklabs=['0', '25', '50', '75', '100', '125', '150'],
             ylab='Genome size (Mb)')
plt.savefig('../figures/SLB21-Fig4b.png')
plt.show()

# %% [code]
```
