## AdditionalFile2 for "iLoci: Robust evaluation of genome content and organization for provisional and mature genome assemblies": make-F5-F6.ipynb

```
# %% [code]
%matplotlib inline

import pandas as pd
import matplotlib.pyplot as plt
import numpy as np
import matplotlib
import seaborn
import json

#Set debug = True to print out variable values for verification:
debug = False

# %% [code]
def draw_hist(alldata, numbins=25, mybins=None, density=False, stacked=False,
              xlabel=None, xlim=None, xticks=None, xticklabs=None,
              ylabel=None, ylim=None, yticks=None, yticklabs=None,
              palette=None, Title=None):
    if not mybins:
        mybins = numbins
        if xlim:
            mybins = np.linspace(xlim[0], xlim[1], numbins)

    fig, ax = plt.subplots()
    
    plot = ax.hist(alldata, bins=mybins, color=palette, density=density, stacked=stacked,
                   label=["Old All", "New All", "Mapped", "Old Unique", "New Unique"])

    if xlabel is not None:
        _ = ax.set_xlabel(xlabel)
    if xticks is not None:
        _ = ax.set_xticks(xticks)
    if xticklabs is not None:
        _ = ax.set_xticklabels(xticklabs)
    if ylabel is not None:
        _ = ax.set_ylabel(ylabel)
    if yticks is not None:
        _ = ax.set_yticks(yticks)
    if yticklabs is not None:
        _ = ax.set_yticklabels(yticklabs)
    if xlim:
        _ = ax.set_xlim(xlim[0], xlim[1])
    if ylim:
        _ = ax.set_ylim(ylim[0], ylim[1])
    _ = ax.legend(loc='upper right', ncol=1, fontsize=16)
    # Cleanup
    seaborn.despine(left=True)
    if Title is not None:
        _ = matplotlib.pyplot.title(Title, fontsize=24)
        _ = matplotlib.pyplot.suptitle("")
    else:
        _ = matplotlib.pyplot.title("")
        _ = matplotlib.pyplot.suptitle("")

# %% [code]
#Importing status counts, query/subject relations, and iLoci information
# from external files:
#
iltypes = ['ci','fi','ii','ni','si']


#For Arabidopsis thaliana: variables statuscountsAtt6 etc. contain the results
# of mapping Att6 onto Atha; variables statuscountsAtha etc. contain the
# results of the reverse mapping, Atha onto Att6

statuscountsAtt6 = {}
qsrelationsAtt6  = {}
qsrelationsAtt6['all'] = {}
statuscountsAtha = {}
qsrelationsAtha  = {}
qsrelationsAtha['all'] = {}

for iltype in iltypes:
    with open('../comparisons/Atha/Att6.' + iltype + 'loci-vs-Atha.summary.counts', 'r') as f:
        statuscountsAtt6[iltype] = json.load(f)
    with open('../comparisons/Atha/Atha.' + iltype + 'loci-vs-Att6.summary.counts', 'r') as f:
        statuscountsAtha[iltype] = json.load(f)
    if debug:
        print("\n\nstatuscountsAtha %sloci:\n" % iltype)
        print(statuscountsAtha[iltype])
    with open('../comparisons/Atha/Att6.' + iltype + 'loci-vs-Atha.summary.qsrelations', 'r') as f:
        qsrelationsAtt6[iltype] = json.load(f)
        qsrelationsAtt6['all'].update(qsrelationsAtt6[iltype])        
    with open('../comparisons/Atha/Atha.' + iltype + 'loci-vs-Att6.summary.qsrelations', 'r') as f:
        qsrelationsAtha[iltype] = json.load(f)
        qsrelationsAtha['all'].update(qsrelationsAtha[iltype])        
    if debug:
        print("\n\nAtha %sloci-qsrelations" % iltype)
        print(qsrelationsAtha[iltype])

ilociAtt6 = pd.read_csv('../comparisons/Atha/Att6.iloci.tsv',sep='\t')
ilociAtha = pd.read_csv('../comparisons/Atha/Atha.iloci.tsv',sep='\t')
ilociAtt6['LogLength'] = np.log10(ilociAtt6['Length'])
ilociAtha['LogLength'] = np.log10(ilociAtha['Length'])


#For Apis mellifera: variables statuscountsAm45 etc. contain the results
# of mapping Am45 onto Amel; variables statuscountsAmel etc. contain the
# results of the reverse mapping, Amel onto Am45

statuscountsAm45 = {}
qsrelationsAm45 = {}
qsrelationsAm45['all'] = {}
statuscountsAmel = {}
qsrelationsAmel  = {}
qsrelationsAmel['all'] = {}

for iltype in iltypes:
    with open('../comparisons/Amel/Am45.' + iltype + 'loci-vs-Amel.summary.counts', 'r') as f:
        statuscountsAm45[iltype] = json.load(f)
    with open('../comparisons/Amel/Amel.' + iltype + 'loci-vs-Am45.summary.counts', 'r') as f:
        statuscountsAmel[iltype] = json.load(f)
    if debug:
        print("\n\nstatuscountsAmel %sloci:\n" % iltype)
        print(statuscountsAmel[iltype])

    with open('../comparisons/Amel/Am45.' + iltype + 'loci-vs-Amel.summary.qsrelations', 'r') as f:
        qsrelationsAm45[iltype] = json.load(f)
        qsrelationsAm45['all'].update(qsrelationsAm45[iltype])
    with open('../comparisons/Amel/Amel.' + iltype + 'loci-vs-Am45.summary.qsrelations', 'r') as f:
        qsrelationsAmel[iltype] = json.load(f)
        qsrelationsAmel['all'].update(qsrelationsAmel[iltype])
    if debug:
        print("\n\nAmel %sloci-qsrelations" % iltype)
        print(qsrelationsAmel[iltype])

ilociAm45 = pd.read_csv('../comparisons/Amel/Am45.iloci.tsv',sep='\t')
ilociAmel = pd.read_csv('../comparisons/Amel/Amel.iloci.tsv',sep='\t')
ilociAm45['LogLength'] = np.log10(ilociAm45['Length'])
ilociAmel['LogLength'] = np.log10(ilociAmel['Length'])


if debug:
    print("\n\nilociAm45:\n")
    print(ilociAm45)
    print("\n\niLociAm45 iiLoci only:\n")
    print(ilociAmel[ilociAmel.LocusClass == "iiLocus"])

    print("\n\nilociAtha:\n")
    print(ilociAtha)
    print("\n\niLociAtha iiLoci only:\n")
    print(ilociAtha[ilociAtha.LocusClass == "iiLocus"])
    
    print("\n\nqsrelationsAm45:\n")
    print(len(qsrelationsAm45['all']))
    print("\n\nSample entries:\n\n")
    for i in range(4):
        print("key: ", list(qsrelationsAm45['all'])[i], "\t",
              "value: ", qsrelationsAm45['all'].get(list(qsrelationsAm45['all'])[i]))

# %% [code]
#Arabidopsis iLoci mapping status

mydf = pd.DataFrame(data = statuscountsAtt6)
mydf['all i'] = mydf.sum(axis=1)
Att6VsAtha = mydf.transpose()
Att6VsAtha.columns = Att6VsAtha.columns.str.replace('qloci_','')
Att6VsAtha.index = Att6VsAtha.index.str.replace('i','iLoci')
Att6VsAtha = Att6VsAtha.rename(index={'iLociiLoci':'iiLoci'}, columns={'wohits':'w/o hits'})
Att6VsAtha = Att6VsAtha[['all','w/o hits','unmapped','mapped','conserved','unconserved','contained','anchored','redefined']]
Att6VsAtha = Att6VsAtha.loc[ ['siLoci', 'niLoci', 'ciLoci', 'iiLoci', 'fiLoci', 'all iLoci'], :]
print("\n\nAtt6 iLoci mapped to Atha:")
display(Att6VsAtha)  
tablefp = open("../tables/SLB21-SuppTable6a-mapping-Atha.tex", "wt")
tablefp.write(Att6VsAtha.to_latex(index=True))
tablefp.close()  
    
mydf = pd.DataFrame(data = statuscountsAtha)
mydf['all i'] = mydf.sum(axis=1)
AthaVsAtt6 = mydf.transpose()
AthaVsAtt6.columns = AthaVsAtt6.columns.str.replace('qloci_','')
AthaVsAtt6.index = AthaVsAtt6.index.str.replace('i','iLoci')
AthaVsAtt6 = AthaVsAtt6.rename(index={'iLociiLoci':'iiLoci'}, columns={'wohits':'w/o hits'})
AthaVsAtt6 = AthaVsAtt6[['all','w/o hits','unmapped','mapped','conserved','unconserved','contained','anchored','redefined']]
AthaVsAtt6 = AthaVsAtt6.loc[ ['siLoci', 'niLoci', 'ciLoci', 'iiLoci', 'fiLoci', 'all iLoci'], :]
print("\n\nAtha iLoci mapped to Att6:")
display(AthaVsAtt6)
tablefp = open("../tables/SLB21-SuppTable6b-mapping-Atha.tex", "wt")
tablefp.write(AthaVsAtt6.to_latex(index=True))
tablefp.close()

# %% [code]
#Apis mellifera mapping status

mydf = pd.DataFrame(data = statuscountsAm45)
mydf['all i'] = mydf.sum(axis=1)
Am45VsAmel = mydf.transpose()
Am45VsAmel.columns = Am45VsAmel.columns.str.replace('qloci_','')
Am45VsAmel.index = Am45VsAmel.index.str.replace('i','iLoci')
Am45VsAmel = Am45VsAmel.rename(index={'iLociiLoci':'iiLoci'}, columns={'wohits':'w/o hits'})
Am45VsAmel = Am45VsAmel[['all','w/o hits','unmapped','mapped','conserved','unconserved','contained','anchored','redefined']]
Am45VsAmel = Am45VsAmel.loc[ ['siLoci', 'niLoci', 'ciLoci', 'iiLoci', 'fiLoci', 'all iLoci'], :]
print("\n\nAmel4.5 iLoci mapped to AmelHAv3.1:")
display(Am45VsAmel)
tablefp = open("../tables/SLB21-SuppTable6c-mapping-Amel.tex", "wt")
tablefp.write(Am45VsAmel.to_latex(index=True))
tablefp.close()
    

mydf = pd.DataFrame(data = statuscountsAmel)
mydf['all i'] = mydf.sum(axis=1)
AmelVsAm45 = mydf.transpose()
AmelVsAm45.columns = AmelVsAm45.columns.str.replace('qloci_','')
AmelVsAm45.index = AmelVsAm45.index.str.replace('i','iLoci')
AmelVsAm45 = AmelVsAm45.rename(index={'iLociiLoci':'iiLoci'}, columns={'wohits':'w/o hits'})
AmelVsAm45 = AmelVsAm45[['all','w/o hits','unmapped','mapped','conserved','unconserved','contained','anchored','redefined']]
AmelVsAm45 = AmelVsAm45.loc[ ['siLoci', 'niLoci', 'ciLoci', 'iiLoci', 'fiLoci', 'all iLoci'], :]
print("\n\nAmelHAv3.1 iLoci mapped to Amel4.5:")
display(AmelVsAm45)
tablefp = open("../tables/SLB21-SuppTable6d-mapping-Amel.tex", "wt")
tablefp.write(AmelVsAm45.to_latex(index=True))
tablefp.close()

# %% [code]
#Visualization of Arabidopsis iLoci mapping status: TAIR6 to TAIR10

Att6VsAtha['Unique to TAIR6'] = Att6VsAtha['w/o hits'] + Att6VsAtha['unmapped']
bdf = Att6VsAtha.loc[['siLoci', 'iiLoci', 'niLoci', 'ciLoci', 'fiLoci'],
                     ['conserved', 'contained', 'anchored', 'redefined', 'Unique to TAIR6']]
print(bdf)

#We multiply small number for purposes of visualization on the same plot,
# changing the index names appropriately:
bdf.loc[['niLoci'], :] = 10 * bdf.loc[['niLoci'], :]
bdf.loc[['ciLoci', 'fiLoci'], :] = 1000 * bdf.loc[['ciLoci', 'fiLoci'], :]
bdf = bdf.rename(index = {'niLoci' : r"niLoci $\times 10$",
                'ciLoci' : r"ciLoci $\times$ 1000", 'fiLoci' : r"fiLoci $\times 1000$"})

plot = bdf.plot.bar(stacked = True, color = ['red','blue', 'purple', 'cyan', 'black'],
                    fontsize=16, rot=0, figsize=(15,10))
_ = plot.set_title("Arabidopsis thaliana (TAIR6 to TAIR10)", fontsize = 28)
_ = plot.legend(prop={'size': 16})
_ = plt.xlabel('')
_ = plt.ylabel('iLocus count')
_ = plot.yaxis.label.set_fontsize(18)
plotfile = "../figures/SLB21-Fig5a.png"
plt.savefig(plotfile)
_ = plt.show()

# %% [code]
#Visualization of Arabidopsis iLoci mapping status: TAIR10 to TAIR6

AthaVsAtt6['Unique to TAIR10'] = AthaVsAtt6['w/o hits'] + AthaVsAtt6['unmapped']
bdf = AthaVsAtt6.loc[['siLoci', 'iiLoci', 'niLoci', 'ciLoci', 'fiLoci'],
                     ['conserved', 'contained', 'anchored', 'redefined', 'Unique to TAIR10']]
print(bdf)

#We multiply small number for purposes of visualization on the same plot,
# changing the index names appropriately:
bdf.loc[['ciLoci'], :] = 10 * bdf.loc[['ciLoci'], :]
bdf.loc[['fiLoci'], :] = 1000 * bdf.loc[['fiLoci'], :]
bdf = bdf.rename(index = {'ciLoci' : r"ciLoci $\times 10$", 'fiLoci' : r"fiLoci $\times 1000$"})

plot = bdf.plot.bar(stacked = True, color = ['red','blue', 'purple', 'cyan', 'black'],
                    fontsize=16, rot=0, figsize=(15,10))
_ = plot.set_title("Arabidopsis thaliana (TAIR10 to TAIR6)", fontsize = 28)
_ = plot.legend(prop={'size': 16})
_ = plt.xlabel('')
_ = plt.ylabel('iLocus count')
_ = plot.yaxis.label.set_fontsize(18)

# %% [code]
#Visualization of Apis mellifera iLoci mapping status: 4.5 to HAv3.1

Am45VsAmel['Unique to Amel4.5'] = Am45VsAmel['w/o hits'] + Am45VsAmel['unmapped']
bdf = Am45VsAmel.loc[['siLoci', 'iiLoci', 'niLoci', 'ciLoci', 'fiLoci'],
                     ['conserved', 'contained', 'anchored', 'redefined', 'Unique to Amel4.5']]
print(bdf)

#We multiply small number for purposes of visualization on the same plot,
# changing the index names appropriately:
bdf.loc[['ciLoci'], :] = 10 * bdf.loc[['ciLoci'], :]
bdf = bdf.rename(index = {'ciLoci' : r"ciLoci $\times 10$"})

plot = bdf.plot.bar(stacked = True, color = ['red','blue', 'purple', 'cyan', 'black'],
                    fontsize=16, rot=0, figsize=(15,10))
_ = plot.set_title("Apis mellifera (Amel4.5 to HAv3.1)", fontsize = 28)
_ = plot.legend(prop={'size': 16})
_ = plt.xlabel('')
_ = plt.ylabel('iLocus count')
_ = plot.yaxis.label.set_fontsize(18)
plotfile = "../figures/SLB21-Fig5b.png"
plt.savefig(plotfile)
_ = plt.show()

# %% [code]
#Visualization of Apis mellifera iLoci mapping status: HAv3.1 to 4.5

AmelVsAm45['Unique to AmelHAv3.1'] = AmelVsAm45['w/o hits'] + AmelVsAm45['unmapped']
bdf = AmelVsAm45.loc[['siLoci', 'iiLoci', 'niLoci', 'ciLoci', 'fiLoci'],
                     ['conserved', 'contained', 'anchored', 'redefined', 'Unique to AmelHAv3.1']]
print(bdf)

#We multiply small number for purposes of visualization on the same plot,
# changing the index names appropriately:
bdf.loc[['ciLoci'], :] = 10 * bdf.loc[['ciLoci'], :]
bdf.loc[['fiLoci'], :] = 10 * bdf.loc[['fiLoci'], :]
bdf = bdf.rename(index = {'ciLoci' : r"ciLoci $\times 10$", 'fiLoci' : r"fiLoci $\times 10$"})

plot = bdf.plot.bar(stacked = True, color = ['red','blue', 'purple', 'cyan', 'black'],
                    fontsize=16, rot=0, figsize=(15,10))
_ = plot.set_title("Apis mellifera (AmelHAv3.1 to Amel4.5)", fontsize = 28)
_ = plot.legend(prop={'size': 16})
_ = plt.xlabel('')
_ = plt.ylabel('iLocus count')
_ = plot.yaxis.label.set_fontsize(18)

# %% [code]
#Exploring target iLocus type for mapped Att6 iLoci

z = zip(ilociAtha['LocusId'], ilociAtha['LocusClass'])
AthaLC = dict(z)

qltypes = ['si', 'ni', 'ci', 'ii', 'fi']
sttypes = ['conserved', 'contained_in', 'anchored_by', 'redefined_to']
iltypes = ['siLocus', 'niLocus', 'ciLocus', 'iiLocus', 'fiLocus']

myd = {}
for qltype in qltypes:
    q = qsrelationsAtt6[qltype]
    for status in sttypes:
        qs = { key:value for key, value in q.items() if value[0] == status }
        qsl = list(qs.values())
        ftarget = [el[1] for el in qsl]
        targetclass = []
        for i in range(0,len(ftarget)):
            targetclass.append(AthaLC[ftarget[i]])
        tmpl = [len(ftarget)]
        for iltype in iltypes:
            tmpl.append(len([i for i, x in enumerate(targetclass) if x == iltype]))
        myd.update({qltype + '-' + status : tmpl})

mydf = pd.DataFrame.from_dict(myd,orient = 'index')
mydf.columns = ['All'] + iltypes

mydf = mydf.rename(index = {'si-conserved' : 'TAIR6 siLoci conserved',
                            'si-contained_in' : 'contained in',
                            'si-anchored_by' : 'anchored by',
                            'si-redefined_to' : 'redefined to',
                            'ni-conserved' : 'TAIR6 niLoci conserved',
                            'ni-contained_in' : 'contained in',
                            'ni-anchored_by' : 'anchored by',
                            'ni-redefined_to' : 'redefined to',
                            'ci-conserved' : 'TAIR6 ciLoci conserved',
                            'ci-contained_in' : 'contained in',
                            'ci-anchored_by' : 'anchored by',
                            'ci-redefined_to' : 'redefined to',
                            'ii-conserved' : 'TAIR6 iiLoci conserved',
                            'ii-contained_in' : 'contained in',
                            'ii-anchored_by' : 'anchored by',
                            'ii-redefined_to' : 'redefined to',
                            'fi-conserved' : 'TAIR6 fiLoci conserved',
                            'fi-contained_in' : 'contained in',
                            'fi-anchored_by' : 'anchored by',
                            'fi-redefined_to' : 'redefined to'
                   },
                  columns = {'siLocus' : 'TAIR10 siLocus', 'niLocus' : 'TAIR10 niLocus',
                             'ciLocus' : 'TAIR10 ciLocus', 'iiLocus' : 'TAIR10 iiLocus',
                             'fiLocus' : 'TAIR10 fiLocus'})
display(mydf)
tablefp = open("../tables/SLB21-SuppTable7a-conservation-Atha.tex", "wt")
tablefp.write(mydf.to_latex(index=True))
tablefp.close()

# %% [code]
#Exploring target iLocus type for mapped Atha iLoci

z = zip(ilociAtt6['LocusId'], ilociAtt6['LocusClass'])
Att6LC = dict(z)

qltypes = ['si', 'ni', 'ci', 'ii', 'fi']
sttypes = ['conserved', 'contained_in', 'anchored_by', 'redefined_to']
iltypes = ['siLocus', 'niLocus', 'ciLocus', 'iiLocus', 'fiLocus']

myd = {}
for qltype in qltypes:
    q = qsrelationsAtha[qltype]
    for status in sttypes:
        qs = { key:value for key, value in q.items() if value[0] == status }
        qsl = list(qs.values())
        ftarget = [el[1] for el in qsl]
        targetclass = []
        for i in range(0,len(ftarget)):
            targetclass.append(Att6LC[ftarget[i]])
        tmpl = [len(ftarget)]
        for iltype in iltypes:
            tmpl.append(len([i for i, x in enumerate(targetclass) if x == iltype]))
        myd.update({qltype + '-' + status : tmpl})
        
mydf = pd.DataFrame.from_dict(myd,orient = 'index')
mydf.columns = ['All'] + iltypes

mydf = mydf.rename(index = {'si-conserved' : 'TAIR10 siLoci conserved',
                            'si-contained_in' : 'contained in',
                            'si-anchored_by' : 'anchored by',
                            'si-redefined_to' : 'redefined to',
                            'ni-conserved' : 'TAIR10 niLoci conserved',
                            'ni-contained_in' : 'contained in',
                            'ni-anchored_by' : 'anchored by',
                            'ni-redefined_to' : 'redefined to',
                            'ci-conserved' : 'TAIR10 ciLoci conserved',
                            'ci-contained_in' : 'contained in',
                            'ci-anchored_by' : 'anchored by',
                            'ci-redefined_to' : 'redefined to',
                            'ii-conserved' : 'TAIR10 iiLoci conserved',
                            'ii-contained_in' : 'contained in',
                            'ii-anchored_by' : 'anchored by',
                            'ii-redefined_to' : 'redefined to',
                            'fi-conserved' : 'TAIR10 fiLoci conserved',
                            'fi-contained_in' : 'contained in',
                            'fi-anchored_by' : 'anchored by',
                            'fi-redefined_to' : 'redefined to'
                   },
                  columns = {'siLocus' : 'TAIR6 siLocus', 'niLocus' : 'TAIR6 niLocus',
                             'ciLocus' : 'TAIR6 ciLocus', 'iiLocus' : 'TAIR6 iiLocus',
                             'fiLocus' : 'TAIR6 fiLocus'})
display(mydf)

# %% [code]
#Exploring target iLocus type for mapped Am45 iLoci

z = zip(ilociAmel['LocusId'], ilociAmel['LocusClass'])
AmelLC = dict(z)

qltypes = ['si', 'ni', 'ci', 'ii', 'fi']
sttypes = ['conserved', 'contained_in', 'anchored_by', 'redefined_to']
iltypes = ['siLocus', 'niLocus', 'ciLocus', 'iiLocus', 'fiLocus']

myd = {}
for qltype in qltypes:
    q = qsrelationsAm45[qltype]
    for status in sttypes:
        qs = { key:value for key, value in q.items() if value[0] == status }
        qsl = list(qs.values())
        ftarget = [el[1] for el in qsl]
        targetclass = []
        for i in range(0,len(ftarget)):
            targetclass.append(AmelLC[ftarget[i]])
        tmpl = [len(ftarget)]
        for iltype in iltypes:
            tmpl.append(len([i for i, x in enumerate(targetclass) if x == iltype]))
        myd.update({qltype + '-' + status : tmpl})
        
mydf = pd.DataFrame.from_dict(myd,orient = 'index')
mydf.columns = ['All'] + iltypes

mydf = mydf.rename(index = {'si-conserved' : 'Am45 siLoci conserved',
                            'si-contained_in' : 'contained in',
                            'si-anchored_by' : 'anchored by',
                            'si-redefined_to' : 'redefined to',
                            'ni-conserved' : 'Am45 niLoci conserved',
                            'ni-contained_in' : 'contained in',
                            'ni-anchored_by' : 'anchored by',
                            'ni-redefined_to' : 'redefined to',
                            'ci-conserved' : 'Am45 ciLoci conserved',
                            'ci-contained_in' : 'contained in',
                            'ci-anchored_by' : 'anchored by',
                            'ci-redefined_to' : 'redefined to',
                            'ii-conserved' : 'Am45 iiLoci conserved',
                            'ii-contained_in' : 'contained in',
                            'ii-anchored_by' : 'anchored by',
                            'ii-redefined_to' : 'redefined to',
                            'fi-conserved' : 'Am45 fiLoci conserved',
                            'fi-contained_in' : 'contained in',
                            'fi-anchored_by' : 'anchored by',
                            'fi-redefined_to' : 'redefined to'
                   },
                  columns = {'siLocus' : 'AmHAv3.1 siLocus', 'niLocus' : 'AmHAv3.1 niLocus',
                             'ciLocus' : 'AmHAv3.1 ciLocus', 'iiLocus' : 'AmHAv3.1 iiLocus',
                             'fiLocus' : 'AmHAv3.1 fiLocus'})
display(mydf)
tablefp = open("../tables/SLB21-SuppTable7b-conservation-Amel.tex", "wt")
tablefp.write(mydf.to_latex(index=True))
tablefp.close()

# %% [code]
#Exploring target iLocus type for mapped Amel iLoci

z = zip(ilociAm45['LocusId'], ilociAm45['LocusClass'])
Am45LC = dict(z)

qltypes = ['si', 'ni', 'ci', 'ii', 'fi']
sttypes = ['conserved', 'contained_in', 'anchored_by', 'redefined_to']
iltypes = ['siLocus', 'niLocus', 'ciLocus', 'iiLocus', 'fiLocus']

myd = {}
for qltype in qltypes:
    q = qsrelationsAmel[qltype]
    for status in sttypes:
        qs = { key:value for key, value in q.items() if value[0] == status }
        qsl = list(qs.values())
        ftarget = [el[1] for el in qsl]
        targetclass = []
        for i in range(0,len(ftarget)):
            targetclass.append(Am45LC[ftarget[i]])
        tmpl = [len(ftarget)]
        for iltype in iltypes:
            tmpl.append(len([i for i, x in enumerate(targetclass) if x == iltype]))
        myd.update({qltype + '-' + status : tmpl})
        
mydf = pd.DataFrame.from_dict(myd,orient = 'index')
mydf.columns = ['All'] + iltypes

mydf = mydf.rename(index = {'si-conserved' : 'AmHAv3.1 siLoci conserved',
                            'si-contained_in' : 'contained in',
                            'si-anchored_by' : 'anchored by',
                            'si-redefined_to' : 'redefined to',
                            'ni-conserved' : 'AmHAv3.1 niLoci conserved',
                            'ni-contained_in' : 'contained in',
                            'ni-anchored_by' : 'anchored by',
                            'ni-redefined_to' : 'redefined to',
                            'ci-conserved' : 'AmHAv3.1 ciLoci conserved',
                            'ci-contained_in' : 'contained in',
                            'ci-anchored_by' : 'anchored by',
                            'ci-redefined_to' : 'redefined to',
                            'ii-conserved' : 'AmHAv3.1 iiLoci conserved',
                            'ii-contained_in' : 'contained in',
                            'ii-anchored_by' : 'anchored by',
                            'ii-redefined_to' : 'redefined to',
                            'fi-conserved' : 'AmHAv3.1 fiLoci conserved',
                            'fi-contained_in' : 'contained in',
                            'fi-anchored_by' : 'anchored by',
                            'fi-redefined_to' : 'redefined to'
                   },
                  columns = {'siLocus' : 'Am45 siLocus', 'niLocus' : 'Am45 niLocus',
                             'ciLocus' : 'Am45 ciLocus', 'iiLocus' : 'Am45 iiLocus',
                             'fiLocus' : 'Am45 fiLocus'})
display(mydf)

# %% [code]
#Exploring whether either very short or very long iLoci are more prone to change ...
#TAIR6 mapped to TAIR10

mapped = []
matched_new = []

for key in qsrelationsAtt6['all']:
    mapped.append(key)
    for ilocus in qsrelationsAtt6['all'][key][1:]:
        matched_new.append(ilocus)    

Att6_iloci = set(ilociAtt6.LocusId)
Atha_iloci = set(ilociAtha.LocusId)

mapped = set(mapped)
matched_new = set(matched_new)
unique_old = Att6_iloci - mapped
unique_new = Atha_iloci - matched_new

Att6_mapped = ilociAtt6.loc[(ilociAtt6.LocusId.isin(mapped))]
Att6_unique = ilociAtt6.loc[(ilociAtt6.LocusId.isin(unique_old))]
Atha_unique = ilociAtha.loc[(ilociAtha.LocusId.isin(unique_new))]

data = [ilociAtt6['LogLength'], ilociAtha['LogLength'],
        Att6_mapped['LogLength'], Att6_unique['LogLength'], Atha_unique['LogLength']]
        
seaborn.set_context({"figure.figsize": (12, 6)})
plot = draw_hist(data, ylabel='Frequency', numbins=8, density=False,
          palette=seaborn.color_palette("hls", len(data)),
          xlabel='iLocus Length (bp)', xlim=(2.5, 5.0),
          xticks=(2.7, 3.0, 3.4, 3.7, 4.0, 4.4, 4.7, 5.0),
          xticklabs=('500', '1,000', '2,500', '5,000', '10,000', '25,000', '50,000', '100,000'),
          Title="Arabidopsis thaliana (TAIR6 vs. TAIR10)")
plot = draw_hist(data, ylabel='Density', numbins=8, density=True,
          palette=seaborn.color_palette("hls", len(data)),
          xlabel='iLocus Length (bp)', xlim=(2.5, 5.0),
          xticks=(2.7, 3.0, 3.4, 3.7, 4.0, 4.4, 4.7, 5.0),
          xticklabs=('500', '1,000', '2,500', '5,000', '10,000', '25,000', '50,000', '100,000'),
          Title="Arabidopsis thaliana (TAIR6 vs. TAIR10)")
plotfile = "../figures/SLB21-Fig6a.png"
plt.savefig(plotfile)
_ = plt.show()

# %% [code]
#... now checking the reverse mapping, TAIR10 onto TAIR6

mapped = []
matched_new = []

for key in qsrelationsAtha['all']:
    mapped.append(key)
    for ilocus in qsrelationsAtha['all'][key][1:]:
        matched_new.append(ilocus)

Atha_iloci = set(ilociAtha.LocusId)
Att6_iloci = set(ilociAtt6.LocusId)
mapped = set(mapped)
matched_new = set(matched_new)
unique_old = Atha_iloci - mapped
unique_new = Att6_iloci - matched_new

Atha_mapped = ilociAtha.loc[(ilociAtha.LocusId.isin(mapped))]
Atha_unique = ilociAtha.loc[(ilociAtha.LocusId.isin(unique_old))]
Att6_unique = ilociAtt6.loc[(ilociAtt6.LocusId.isin(unique_new))]

data = [ilociAtha['LogLength'], ilociAtt6['LogLength'],
        Atha_mapped['LogLength'], Atha_unique['LogLength'], Att6_unique['LogLength']]

seaborn.set_context({"figure.figsize": (12, 6)})
plot = draw_hist(data, ylabel='Frequency', numbins=10, density=False,
          palette=seaborn.color_palette("hls", len(data)),
          xlabel='iLocus Length (bp)', xlim=(2.5, 5.0),
          xticks=(2.7, 3.0, 3.4, 3.7, 4.0, 4.4, 4.7, 5.0),
          xticklabs=('500', '1,000', '2,500', '5,000', '10,000', '25,000', '50,000', '100,000'),
          Title="Arabidopsis thaliana (TAIR10 vs. TAIR6)")
plot = draw_hist(data, ylabel='Density', numbins=10, density=True,
          palette=seaborn.color_palette("hls", len(data)),
          xlabel='iLocus Length (bp)', xlim=(2.5, 5.0),
          xticks=(2.7, 3.0, 3.4, 3.7, 4.0, 4.4, 4.7, 5.0),
          xticklabs=('500', '1,000', '2,500', '5,000', '10,000', '25,000', '50,000', '100,000'),
          Title="Arabidopsis thaliana (TAIR10 vs. TAIR6)")
_ = plt.show()

# %% [code]
#... Apis mellifera Amel4.5 mapped onto AmelHAv3.1

mapped = []
matched_new = []

for key in qsrelationsAm45['all']:
    mapped.append(key)
    for ilocus in qsrelationsAm45['all'][key][1:]:
        matched_new.append(ilocus)

Am45_iloci = set(ilociAm45.LocusId)
Amel_iloci = set(ilociAmel.LocusId)
mapped = set(mapped)
matched_new = set(matched_new)
unique_old = Am45_iloci - mapped
unique_new = Amel_iloci - matched_new
Amel_mapped = ilociAm45.loc[(ilociAm45.LocusId.isin(mapped))]
Am45_unique = ilociAm45.loc[(ilociAm45.LocusId.isin(unique_old))]
Amel_unique = ilociAmel.loc[(ilociAmel.LocusId.isin(unique_new))]
data = [ilociAm45['LogLength'], ilociAmel['LogLength'],
        Amel_mapped['LogLength'], Am45_unique['LogLength'], Amel_unique['LogLength']]

seaborn.set_context({"figure.figsize": (12, 6)})
draw_hist(data, ylabel='Frequency', numbins=8, density=False,
          palette=seaborn.color_palette("hls", len(data)),
          xlabel='iLocus Length (bp)', xlim=(2.5, 5.0),
          xticks=(2.7, 3.0, 3.4, 3.7, 4.0, 4.4, 4.7, 5.0),
          xticklabs=('500', '1,000', '2,500', '5,000', '10,000', '25,000', '50,000', '100,000'),
          Title="Apis mellifera (4.5 vs. HAv3.1)")
draw_hist(data, ylabel='Density', numbins=8, density=True,
          palette=seaborn.color_palette("hls", len(data)),
          xlabel='iLocus Length (bp)', xlim=(2.5, 5.0),
          xticks=(2.7, 3.0, 3.4, 3.7, 4.0, 4.4, 4.7, 5.0),
          xticklabs=('500', '1,000', '2,500', '5,000', '10,000', '25,000', '50,000', '100,000'),
          Title="Apis mellifera (4.5 vs. HAv3.1)")
plotfile = "../figures/SLB21-Fig6b.png"
plt.savefig(plotfile)
_ = plt.show()

# %% [code]
```
