## AdditionalFile2 for "iLoci: Robust evaluation of genome content and organization for provisional and mature genome assemblies": make-SF1.ipynb

```
# %% [code]
%matplotlib inline

import matplotlib
import pandas as pd
import matplotlib.pyplot as plt
import seaborn

matplotlib.rcParams['axes.labelsize'] = 16
matplotlib.rcParams['xtick.labelsize'] = 12
matplotlib.rcParams['ytick.labelsize'] = 12
matplotlib.rcParams['savefig.dpi'] = 300
seaborn.set_context({"figure.figsize": (15, 10)})

# %% [code]
specieslist = ['Scer', 'Cele', 'Crei', 'Mtru', 'Dmel',
               'Agam', 'Xtro', 'Drer', 'Mmus', 'Hsap',
               'Vcar', 'Pdom', 'Dpul', 'Mvit']

# %% [code]
files = ['../data/{}/{}.iloci.tsv'.format(s, s) for s in specieslist]
silgth = []
for f in files:
    data = pd.read_csv(f, sep='\t')
    siloci = data.loc[data.LocusClass.isin(['siLocus'])]
    silgth.append(siloci.Length)
plt.boxplot(silgth, labels = specieslist, showfliers = False)
plt.ylabel('siLocus length (bp)')
plt.yscale('log')
plt.savefig('../figures/SLB21-SuppFig1.png')
plt.show()

# %% [code]
```
