## AdditionalFile2 for "iLoci: Robust evaluation of genome content and organization for provisional and mature genome assemblies": make-SF3.ipynb

```
# %% [code]
from __future__ import division
%matplotlib inline

import pandas
import matplotlib.pyplot as plt
import numpy
import scipy
import seaborn
seaborn.set_context({"figure.figsize": (15, 10)})

# %% [code]
verbose = True
root = '../data-delta0'
orientations = ['FF', 'FR', 'RF', 'RR']
datasource = "" #set to shuffled for randomized data input

if verbose:
    print("Data for stacked bar plot. Significantly non-random 2x2 tables are starred.")
    
for species in ['Scer', 'Cele', 'Crei', 'Mtru', 'Agam', 'Dmel', 'Xtro', 'Drer', 'Mmus', 'Hsap']:
    if (datasource == 'shuffled'):
      datafile = '{}/{}/{}.iloci.shuffled.tsv'.format(root,species,species)
    else:
      datafile = '{}/{}/{}.iloci.tsv'.format(root,species,species)
    data = pandas.read_csv(datafile,sep='\t')
    data['LogLength'] = numpy.log10(data['Length'])
    
    iiloci = data.loc[data.LocusClass == 'iiLocus']
    
    FRmatrix = numpy.zeros((2,2))
    i = 1
    for o in orientations:
        subset = iiloci.loc[(iiloci.FlankGeneOrient == o)]
        if i == 1:
            FRmatrix[0,0] = len(subset)
        if i == 2:
            FRmatrix[0,1] = len(subset)
        if i == 3:
            FRmatrix[1,0] = len(subset)
        if i == 4:
            FRmatrix[1,1] = len(subset)
        i = i + 1
    
    if verbose:
        print("")
        print("Species: ",species)
        print("Overall FR type counts (total: ",len(iiloci),"):")
        print(FRmatrix.astype(int))
        
    subsets = list()
    FRmatrix = numpy.zeros((2,2))
    llist = []
    bins=numpy.log10(numpy.logspace(2, 5, 21))

    #Note: We use 20 bins on 10 species. Thus, the Fisher Exact Test
    #      significance level is conservatively set to 0.01/20.
    
    for b in range(0,20):
        i = 1
        for orient in orientations:
            subset = iiloci.loc[(iiloci.FlankGeneOrient == orient) &
                                (iiloci.LogLength >= bins[b]) &
                                (iiloci.LogLength <  bins[b+1])]
            if i == 1:
              FRmatrix[0,0] = len(subset)
            if i == 2:
              FRmatrix[0,1] = len(subset)
            if i == 3:
              FRmatrix[1,0] = len(subset)
            if i == 4:
              FRmatrix[1,1] = len(subset)
            i = i + 1
        
        oddsration, pvalue = scipy.stats.fisher_exact(FRmatrix.astype(int))
        height = FRmatrix[0,0] + FRmatrix[0,1] + FRmatrix[1,0] + FRmatrix[1,1]
        if pvalue <= 0.01/20:
            psymbol = "*"
        else:
            psymbol = ""
        if verbose:
            print("")
            print("Length bin: ",int(10**bins[b])," to ",int(10**bins[b+1]+1))
            print("Number of iiLoci in this bin: ",int(height))
            print("FR type matrix (",psymbol,"):")
            print(FRmatrix.astype(int))
        llist.append([(bins[b]+bins[b+1])/2,height,psymbol])
    
    for o in orientations:
        subset = iiloci.loc[(iiloci.FlankGeneOrient == o)]
        subsets.append(subset['LogLength'])
    
    fig, axis = plt.subplots()
    _ = plt.hist(subsets, stacked=True,
                 bins=numpy.log10(numpy.logspace(2, 5, 21)),
                 label=orientations, edgecolor='black', linewidth=0.6)
    if (datasource == 'shuffled'):
      _ = plt.title('Shuffled')
    else:
      _ = plt.title('Observed')
    _ = plt.legend()
    _ = axis.set_xlabel('iiLocus Length (bp)', fontsize=16)
    _ = axis.set_ylabel('Frequency', fontsize=16)
    _ = axis.set_xlim((numpy.log10(100), numpy.log10(100000)))
    _ = axis.set_xticks([numpy.log10(100), 3, 4, 5])
    _ = axis.set_xticklabels(['100','1,000', '10,000', '100,000'])
    _ = plt.suptitle(species, fontsize=20)
        
    for i in range(len(llist)):
        axis.text(llist[i][0], llist[i][1] + 5, llist[i][2],
                  fontsize=20, ha='center', va='bottom')
           
    plotfile = "../figures/SLB21-SuppFig3_%s%s.png" % (datasource, species)
    plt.savefig(plotfile)
    _ = plt.show()

# %% [code]
```
