## AdditionalFile2 for "iLoci: Robust evaluation of genome content and organization for provisional and mature genome assemblies": make-SF5b.ipynb

```
# %% [code]
%matplotlib inline

# Boilerplate code: set up the environment
import matplotlib
import matplotlib.pyplot as plt
import pandas
import scipy
import seaborn
colors = ['#a6cee3', '#1f78b4', '#b2df8a', '#33a02c', '#cc33cc',
          '#fb9a99', '#e31a1c', '#fdbf6f']
matplotlib.rcParams['savefig.dpi'] = 300
seaborn.set_context({"figure.figsize": (15, 10)})

# %% [code]
def load_and_prep(filename, specieslist, index='Species'):
    """Boilerplate code: load data into memory"""
    dataframe = pandas.read_csv(filename,sep='\t')
    df = dataframe[dataframe[index].isin(specieslist)]
    df = df.set_index([index]).reindex(specieslist)
    return df

# %% [code]
def calc_rho(species, basedir='../data'):
    """
    Calculate \rho, the proportion of iiLocus length to giLocus length, for a particular genome.
    """
    files = ['{}/{}/{}.iloci.tsv'.format(basedir, s, s) for s in species]
    data = pandas.concat([pandas.read_csv(f,sep='\t') for f in files])
    giloci = data.loc[data.LocusClass.isin(['siLocus', 'ciLocus', 'niLocus'])]
    iiloci = data.loc[data.LocusClass.isin(['iiLocus'])]
    quant = list(giloci['Length'].quantile([0.05, 0.95]))
    giloci_q = giloci.loc[(giloci.Length >= quant[0]) & (giloci.Length <= quant[1])]
    quant = list(iiloci['Length'].quantile([0.05, 0.95]))
    iiloci_q = iiloci.loc[(iiloci.Length >= quant[0]) & (iiloci.Length <= quant[1])]
    rho = iiloci_q['Length'].mean() / giloci_q['Length'].mean()
    return rho

# %% [code]
matplotlib.rcParams['axes.labelsize'] = 16
matplotlib.rcParams['xtick.labelsize'] = 12
matplotlib.rcParams['ytick.labelsize'] = 12
seaborn.set_context({"figure.figsize": (12, 8)})

speciesnames = ['Apis mellifera', 'Anopheles gambiae',
                'Acromyrmex echinatior', 'Drosophila melanogaster',
                'Polistes dominula', 'Nasonia vitripennis']
specieslist = ['Amel', 'Agam', 'Aech', 'Dmel', 'Pdom', 'Nvit']
palette = seaborn.color_palette("hls", len(specieslist))

for filename in ['../tables/phisigma-Pdom-min1Mb.tsv', '../tables/phisigma-Pdom-min2Mb.tsv', '../tables/phisigma-Pdom-min3Mb.tsv']:
    data = pandas.read_csv(filename,'\t')
    plots = list()
    for i, species in enumerate(specieslist):
        specdata = data.loc[(data.Species == species)]
        plot = plt.scatter(specdata['Phi'], specdata['Sigma'], color=palette[i])
        plots.append(plot)
    _ = plt.xlim(-0.02, 1.02)
    _ = plt.ylim(-0.02, 1.02)
    _ = plt.xlabel('φ (Proportion of giLoci merged into miLoci)')
    _ = plt.ylabel('σ (Proportion of sequence occupied by miLoci)')
    _ = plt.legend(plots, speciesnames, scatterpoints=1,
               loc='upper left',
               ncol=2,
               fontsize=12)
    _ = plt.show()

# %% [code]
speciesnames = ['Apis mellifera', 'Anopheles gambiae',
                'Acromyrmex echinatior', 'Drosophila melanogaster',
                'Polistes dominula', 'Nasonia vitripennis']
specieslist = ['Amel', 'Agam', 'Aech', 'Dmel', 'Pdom', 'Nvit']
palette = seaborn.color_palette("hls", len(specieslist))
rho = calc_rho(specieslist, basedir='../data')

data = pandas.read_csv('../tables/phisigma-Pdom-min2Mb.tsv',sep='\t')
plots = list()
for i, species in enumerate(specieslist):
    specdata = data.loc[(data.Species == species)]
    plot = plt.scatter(specdata['Phi'], specdata['Sigma'], color=palette[i])
    plots.append(plot)
    x = [k / 1000 for k in range(1000)]
    y = [(k/(1+(1-k)*rho)) for k in x]
    _ = plt.plot(x, y, color='#cccccc')
_ = plt.xlim(-0.02, 1.02)
_ = plt.ylim(-0.02, 1.02)
_ = plt.xlabel('φ (Proportion of giLoci merged into miLoci)')
_ = plt.ylabel('σ (Proportion of sequence occupied by miLoci)')
_ = plt.legend(plots, speciesnames, scatterpoints=1,
           loc='upper left',
           ncol=2,
           fontsize=12)
plt.savefig('../figures/SLB21-SuppFig5b.png')
plt.show()

# %% [code]
```
