## AdditionalFile2 for "iLoci: Robust evaluation of genome content and organization for provisional and mature genome assemblies": make-SF5c-SF8.ipynb

```
# %% [code]
%matplotlib inline

import glob
import math
import matplotlib
import matplotlib.pyplot as plt
import matplotlib.lines as ml
import numpy
import pandas
import random
import seaborn

matplotlib.rcParams['axes.labelsize'] = 16
matplotlib.rcParams['xtick.labelsize'] = 12
matplotlib.rcParams['ytick.labelsize'] = 12
matplotlib.rcParams['savefig.dpi'] = 300
seaborn.set_context({"figure.figsize": (12, 8)})

# %% [code]
def calc_rho(species, basedir='../data'):
    """
    Calculate \rho, the proportion of iiLocus length to giLocus length,
    for a particular set of genomes.
    """
    files = ['{}/{}/{}.iloci.tsv'.format(basedir, s, s) for s in species]
    data = pandas.concat([pandas.read_csv(f,sep='\t') for f in files])
    giloci = data.loc[data.LocusClass.isin(['siLocus', 'ciLocus', 'niLocus'])]
    iiloci = data.loc[data.LocusClass.isin(['iiLocus'])]
    quant = list(giloci['Length'].quantile([0.05, 0.95]))
    giloci_q = giloci.loc[(giloci.Length >= quant[0]) & (giloci.Length <= quant[1])]
    quant = list(iiloci['Length'].quantile([0.05, 0.95]))
    iiloci_q = iiloci.loc[(iiloci.Length >= quant[0]) & (iiloci.Length <= quant[1])]
    rho = iiloci_q['Length'].mean() / giloci_q['Length'].mean()
    return rho

# %% [code]
speciesnames = ['Daphnia pulex', 'Anopheles gambiae', 'Drosophila melanogaster',
                'Apis mellifera', 'Tribolium castaneum', 'Tetranychus urticae', ]
specieslist = ['Dpul', 'Agam', 'Dmel', 'Amel', 'Tcas', 'Turt']
palette = seaborn.color_palette("hls", len(specieslist))
rho = calc_rho(specieslist, basedir='../data')

data = pandas.read_csv('../tables/phisigma-Dpul.tsv',sep='\t')
plots = list()
for i, species in enumerate(specieslist):
    specdata = data.loc[(data.Species == species)]
    plot = plt.scatter(specdata['Phi'], specdata['Sigma'], color=palette[i])
    plots.append(plot)
    x = [k / 1000 for k in range(1000)]
    y = [(k/(1+(1-k)*rho)) for k in x]
    _ = plt.plot(x, y, color='#cccccc')
_ = plt.xlim(-0.02, 1.02)
_ = plt.ylim(-0.02, 1.02)
_ = plt.xlabel('φ (Proportion of giLoci merged into miLoci)')
_ = plt.ylabel('σ (Proportion of sequence occupied by miLoci)')
_ = plt.legend(plots, speciesnames, scatterpoints=1,
           loc='upper left',
           ncol=2,
           fontsize=12)
plt.savefig('../figures/SLB21-SuppFig5c.png')

# %% [code]
matplotlib.rcParams['axes.labelsize'] = 18
matplotlib.rcParams['xtick.labelsize'] = 14
matplotlib.rcParams['ytick.labelsize'] = 14

# %% [code]
def draw_hist(data, series, species, numbins=25, mybins=None, density=True,
              xlabel=None, xlim=None, xticks=None, xticklabs=None,
              ylabel='Density', ylim=None, yticks=None, yticklabs=None,
              palette=None):
    
    alldata = list()
    for spec in species:
        subdata = data.loc[data.Species == spec]
        alldata.append(subdata[series])

    if not mybins:
        mybins = numbins
        if xlim:
            mybins = numpy.linspace(xlim[0], xlim[1], numbins)

    fig, ax = plt.subplots()
    plot = ax.hist(alldata, bins=mybins, color=palette, density=density, label=species)

    if xlabel is not None:
        _ = ax.set_xlabel(xlabel)
    if xticks is not None:
        _ = ax.set_xticks(xticks)
    if xticklabs is not None:
        _ = ax.set_xticklabels(xticklabs)
    if ylabel is not None:
        _ = ax.set_ylabel(ylabel)
    if yticks is not None:
        _ = ax.set_yticks(yticks)
    if yticklabs is not None:
        _ = ax.set_yticklabels(yticklabs)
    if xlim:
        _ = ax.set_xlim(xlim[0], xlim[1])
    if ylim:
        _ = ax.set_ylim(ylim[0], ylim[1])
    _ = ax.legend(loc='upper right', ncol=1, fontsize=12)
    
    # Cleanup
    seaborn.despine(left=True)
    _ = matplotlib.pyplot.title("")
    _ = matplotlib.pyplot.suptitle("")

# %% [code]
def load_data(pattern, loglength=True):
    filelist = glob.glob(pattern)
    data = pandas.concat([pandas.read_csv(f,sep='\t') for f in filelist])
    data['Length'] = data['Length'].astype(int)
    if loglength:
        data['LogLength'] = numpy.log10(data['Length'])
    return data

# %% [code]
def silocus_mrnas(pattern, silocus_ids):
    filelist = glob.glob(pattern)
    data = pandas.concat([pandas.read_csv(f, sep='\t', header=None, names=('iLocusID', 'MrnaAccession')) for f in filelist])
    subdata = data.loc[data.iLocusID.isin(silocus_ids)]
    return list(subdata['MrnaAccession'])

# %% [markdown]
## Exploring iiLocus space:

# %% [code]
iloci = load_data('../data/*/*.iloci.tsv')
idf = iloci.groupby(['Species']).sum()
iiloci = iloci[iloci['LocusClass'] == 'iiLocus']
iidf = iiloci.groupby(['Species']).sum()
iidf['Occupancy'] = 100.*iidf['EffectiveLength']/idf['EffectiveLength']
display(iidf)

# %% [code]
iiloci.groupby(['Species']).median().sort_values(by='Length')

# %% [code]
seaborn.set_context({"figure.figsize": (12, 6)})
specieslist = ['Agam', 'Dpul', 'Dmel']
draw_hist(iiloci, 'LogLength', species=specieslist,
          palette=seaborn.color_palette("hls", len(specieslist)),
          xlabel='iiLocus length (bp)', xlim=(2.5, 5.5),
          xticks=(3,4,5), xticklabs=('1,000', '10,000', '100,000'))
plt.savefig('../figures/SLB21-SuppFig8-ii-lengths.png')

# %% [markdown]
## Exploring siLocus space:

# %% [code]
siloci = iloci[iloci['LocusClass'] == 'siLocus']
siloci.groupby(['Species']).median().sort_values(by='Length')

# %% [code]
seaborn.set_context({"figure.figsize": (12, 6)})
specieslist = ['Agam', 'Dpul', 'Dmel']
draw_hist(siloci, 'LogLength', species=specieslist,
          palette=seaborn.color_palette("hls", len(specieslist)),
          xlabel='siLocus length (bp)', xlim=(2.75, 5.05),
          xticks=(3,4,5), xticklabs=('1,000', '10,000', '100,000'))
plt.savefig('../figures/SLB21-SuppFig8-si-lengths.png')

# %% [markdown]
## What are the contributions of exon counts per gene and exon length?

# %% [code]
premrnas = load_data('../data/*/*.pre-mrnas.tsv')
silocus_ids = siloci['LocusId']
silocus_mrna_ids = silocus_mrnas('../data/*/*.ilocus.mrnas.tsv', silocus_ids)
silocus_premrnas = premrnas.loc[premrnas.Accession.isin(silocus_mrna_ids)].copy()
silocus_premrnas['ExonCountMax'] = [min(x, 20) for x in silocus_premrnas['ExonCount']]

# %% [code]
silocus_premrnas.groupby(['Species']).median().sort_values(by='ExonCount')

# %% [code]
seaborn.set_context({"figure.figsize": (12, 6)})
specieslist = ['Agam', 'Dpul', 'Dmel']
draw_hist(silocus_premrnas, 'ExonCount', species=specieslist,
          palette=seaborn.color_palette("hls", len(specieslist)),
          mybins=[x - 0.5 for x in range(1, 22)], xlabel='siLocus exon count', xlim=(0, 20.5))
plt.savefig('../figures/SLB21-SuppFig8-si-exon-counts.png')
plt.show()

# %% [code]
exons = load_data('../data/*/*.exons.tsv', loglength=True)
exons = exons.loc[exons.MrnaAcc.isin(silocus_mrna_ids)]
exons.head()

# %% [code]
exons.groupby(['Species']).median().sort_values(by=['Length'])

# %% [code]
seaborn.set_context({"figure.figsize": (12, 6)})
specieslist = ['Agam', 'Dpul', 'Dmel']
draw_hist(exons, 'LogLength', species=specieslist,
          palette=seaborn.color_palette("hls", len(specieslist)),
          xlabel='siLocus exon length (bp)', xlim=(0.5, 4.05),
          xticks=(1,2,3,4), xticklabs=('10', '100', '1,000', '10,000'))
plt.savefig('../figures/SLB21-SuppFig8-si-exon-lengths.png')

# %% [markdown]
## Exploring intron space:

# %% [code]
introns = load_data('../data/*/*.introns.tsv', loglength=True)
introns = introns.loc[introns.MrnaAcc.isin(silocus_mrna_ids)]
introns.head()

# %% [code]
print("Intron counts per species:")
introns.groupby(['Species']).count()['MrnaAcc']

# %% [code]
introns.groupby(['Species']).median().sort_values(by=['Length'])

# %% [code]
seaborn.set_context({"figure.figsize": (12, 6)})
specieslist = ['Agam', 'Dpul', 'Dmel']
draw_hist(introns, 'LogLength', species=specieslist,
          palette=seaborn.color_palette("hls", len(specieslist)),
          xlabel='siLocus intron length (bp)', xlim=(0.75, 5.05),
          xticks=(3,4,5), xticklabs=('1,000', '10,000', '100,000'))
plt.savefig('../figures/SLB21-SuppFig8-si-intron-lengths.png')
```
