## AdditionalFile2 for "iLoci: Robust evaluation of genome content and organization for provisional and mature genome assemblies": README.md

## Documentation
Instructions necessary to produce the results discussed in the paper, from
download of primary data to computations and tables/figures, are given in a
sequence of scripts and Jupyter notebooks as recored in file ./xdoitall.
Each individual analysis should run with no problem on a laptop or desktop
computer, although reproducing all the basic analyses requires about 200 GB
of free disk space for temporary storage of raw and intermediate data files
(the large-scale genome compactness analysis summarizing different clades of
species in the [./taxa](./taxa) subdirectory would require another 350-400 GB).


## Bash scripts
Some long UNIX commands in this documentation are split across multiple lines.
This is done for the sake of readability.
The use of the backslash (`\`) character indicates that the following line is
part of the same command.
For example, the command

```bash
rws fidibus --workdir=data \
        --numprocs=2        \
        --refr=Scer,Cele    \
        download prep
```

is identical to the command

```bash
rws fidibus --workdir=data --numprocs=2 --refr=Scer,Cele download prep
```


## Jupyter notebooks
The easiest way to run the Juypter notebooks described here is by creating a
suitable python environment.
The following provides one way of doing this:

```bash
#Installation of prerequisites (shown for Fedora Linux; other systems will have
#equivalent pacckage managing tools):
sudo dnf install libXcomposite libXcursor libXi libXtst libXrandr \
                 alsa-lib mesa-libEGL libXdamage mesa-libGL libXScrnSaver

#Download and run the Anaconda3 installer from https://www.anaconda.com/, e.g.:
#
wget https://repo.anaconda.com/archive/Anaconda3-2021.05-Linux-x86_64.sh
bash ./Anaconda3-2021.05-Linux-x86_64.sh
\rm  ./Anaconda3-2021.05-Linux-x86_64.sh

#In a new shell:
#
conda config --set auto_activate_base False
conda deactivate

#In a new shell:
#
conda --version
conda info --envs

conda create --name iLoci biopython pycurl pyyaml pytest pytest-cov \
	jupyter pandas seaborn nb_conda_kernels ipykernel r-irkernel
conda activate iLoci
```

and now
```bash
jupyter notebook
```
