## AdditionalFile2 for "iLoci: Robust evaluation of genome content and organization for provisional and mature genome assemblies": README_explore-Chlorophyta.md

# Exploration of green algae (Chlorophyta) genomes

## Computing clusters/hiLoci

We use the fidibus cluster function to cluster iLocus protein products from *Volvox carteri*, 8 other green algae, and 4 land plants, based on the previously determined iLoci.
We then use a custom Python script to assign each iLocus a provisional status based on the proteins with which it clusters, as follows:

- *HighlyConserved*: iLocus has a protein product conserved in all 9 chlorophyte species
- *Conserved*: iLocus has a protein product conserved in at least 4 of the 9 chlorophyte species
- *Matched*: iLocus has a protein product that is conserved in at least on other species, including the outgroups
- *Unmatched*: iLocus has a protein product that is not conserved in any other species

```bash
fidibus --cfgdir=genome_configs --workdir=data --numprocs=13 \
        --refrbatch=Chlorophyta+ \
        cluster
\rm fidibus.prot.fa
mv fidibus.prot         Chlorophyta/Chlorophyta.prot.fa
mv fidibus.prot.clstr   Chlorophyta/Chlorophyta.prot.clstr
mv fidibus.hiloci.tsv   Chlorophyta/Chlorophyta.hiLoci.tsv

python3 Chlorophyta/det-Chlorophyta-status.py \
	Chlorophyta/Chlorophyta.hiLoci.tsv > Chlorophyta/Chlorophyta.hiLocus.pre-status.tsv
```

## Post-processing of unmatched iLoci

The clustering criteria place somewhat strict limits on length differences between proteins.
Many iLoci labeled *Unmatched* by the initial step are therefore not truly orphans, but may be annotated incompletely or incorrectly in one or more species, or may have evolved to such an extent that near-full-length alignments are not possible.
Most of these *Unmatched* iLoci will indeed encode *bona fide* protein products that are likely conserved in other species, but may need additional attention before they can be reliably used for comparative genomics analysis.
This post-processing procedure will distinguish these initially unmatched iLoci from iLoci that have no reliable matches in other species, the former being relabeled as *Matched*, and the latter being relabeled as *Orphan*.

```bash
# Build BLAST databases for refined status determination:
#
mkdir Chlorophyta/blastdbs
for species in Vcar Apro Crei Csub Cvar Mpus Mcom Oluc Otau
do
    makeblastdb -in data/${species}/${species}.prot.fa -dbtype prot \
		-parse_seqids -out Chlorophyta/blastdbs/${species}
done

# Refined analysis:
#
cd Chlorophyta/

for species in Vcar Apro Crei Csub Cvar Mpus Mcom Oluc Otau
do
    # Grab proteins from iLoci initially labeled  "Unmatched":
    #
    grep Unmatched Chlorophyta.hiLocus.pre-status.tsv \
        | grep $species \
        | cut -f 5 \
        > blastdbs/${species}.unmatched.txt
    blastdbcmd -db blastdbs/${species} \
               -entry_batch blastdbs/${species}.unmatched.txt \
        > blastdbs/${species}.unmatched.fa

    # Compose a database of proteins from all species except this one:
    #
    blastdb_aliastool -title Not${species} \
                      -out blastdbs/Not${species} \
                      -dblist_file <(ls blastdbs/*.phr | cut -d"." -f1 | grep -v $species) \
                      -dbtype prot

    # Execute the BLAST searches in the background, so as to use multiple processors:
    #
    blastp -query blastdbs/${species}.unmatched.fa \
           -db blastdbs/Not${species} \
           -num_alignments 5 \
           -num_descriptions 5 \
           -evalue 1e-10 \
           -out blastdbs/${species}.unmatched.blastp &
done
wait
cd ..

for species in Vcar Apro Crei Csub Cvar Mpus Mcom Oluc Otau
do
    # Extract protein IDs of newly matched proteins:
    #
    MuSeqBox -i Chlorophyta/blastdbs/${species}.unmatched.blastp \
             -L 100 -d 16 -l 24 \
        > Chlorophyta/blastdbs/${species}.unmatched.msb
    egrep "^${species}:XP|^${species}:NP" Chlorophyta/blastdbs/${species}.unmatched.msb \
        | awk '{ print $1 }' \
        | sort \
        | uniq \
        > Chlorophyta/blastdbs/${species}.blast_matches.txt
done

# Re-classify "Unmatched" iLoci as "Matched" or "Orphan":
#
python3 Chlorophyta/det-Chlorophyta-status-post-blast.py <(cat Chlorophyta/blastdbs/*.blast_matches.txt) \
    Chlorophyta/Chlorophyta.hiLocus.pre-status.tsv \
    > Chlorophyta/Chlorophyta.hiLocus.status.tsv

for species in Vcar Apro Crei Csub Cvar Mpus Mcom Oluc Otau
do
	cat data/${species}/${species}.iloci.tsv >> Chlorophyta.iloci.tsv
done
```

Finally, we use custom Python scripts to compute a breakdown of each species, for each species showing the number of iLoci assigned to each category, as well as the proportion of the genome occupied by iLoci of that category.
See [make-F4a-F4b-SF4a.ipynb](make-F4a-F4b-SF4a.ipynb) for the code used to plot these breakdowns.

```bash
# The folllowing will produce warnings for iLoci encoding protein sequences
# below the cd-hit "throwaway" threshold (length 10, by default); totally
# reasonable ...
#
python3 Chlorophyta/hiLoci-breakdown.py \
    Chlorophyta.iloci.tsv Chlorophyta/Chlorophyta.hiLocus.status.tsv \
    > Chlorophyta/Chlorophyta-breakdown-bp.tsv
python3 Chlorophyta/hiLoci-breakdown.py --counts \
    Chlorophyta.iloci.tsv Chlorophyta/Chlorophyta.hiLocus.status.tsv \
    > Chlorophyta/Chlorophyta-breakdown-counts.tsv
\rm Chlorophyta.iloci.tsv


# What are the highly conserved iLoci (by protein product)?
#
for pid in $(grep HighlyConserved Chlorophyta/Chlorophyta.hiLocus.status.tsv | grep Crei | cut -f 5)
do
    grep $pid data/Crei/Crei.all.prot.fa
done > Chlorophyta/Crei-deflines-HighlyConserved.txt
```

## Analysis of function of highly conserved piLoci

We examined the functional annotations of proteins encoded by highly conserved piLoci in *Chlamydomonas reinhardtii*.

```bash
# What are the highly conserved iLoci (by protein product)?
#
for pid in $(grep HighlyConserved Chlorophyta/Chlorophyta.hiLocus.status.tsv | grep Crei | cut -f 5)
do
    grep $pid data/Crei/Crei.all.prot.fa
done > Chlorophyta/Crei-deflines-HighlyConserved.txt
```
