## AdditionalFile2 for "iLoci: Robust evaluation of genome content and organization for provisional and mature genome assemblies": README_refr-genome-compactness.md

# Evaluation of genome compactness

We assessed the compactness of the references genomes on two related measures,

- φ (phi), the proportion of giLoci merged into miLoci
- σ (sigma), the proportion of the genome sequence occupied by miLoci

Because these measures are uninformative on small scales, (φ, σ) values were computed only for chromosome or scaffold sequences of at least 1 Mb in length.
Extremely long iiLoci (those in the top 5% of length for each species) and extremely short giLoci (those in the bottom 5%) were discarded as outliers prior to computing (φ, σ).

```bash
fidibus-compact.py --workdir=data/ --length=1000000 \
                  --iqnt=0.95 --gqnt=0.05 \
                  Scer Cele Crei Mtru Agam Dmel Xtro Drer Mmus Hsap \
    > phisigma-refr.tsv
```

(script implementing the above and the commands in the next section:
[run-explore-compactness-refr.sh](./run-explore-compactness-refr.sh)).

## Different values of δ

To evaluate the robustness of the (φ, σ) measures with respect to the δ (delta) parameter, we recomputed iLoci at δ=300 and δ=750 for comparison with the default δ=500.

```bash
fidibus --workdir=data-delta300/ \
        --numprocs=4 \
        --delta=300 \
        --refr=Scer,Cele,Crei,Mtru,Agam,Dmel,Xtro,Drer,Mmus,Hsap \
        download prep iloci breakdown stats

fidibus --workdir=data-delta750/ \
        --numprocs=4 \
        --delta=750 \
        --refr=Scer,Cele,Crei,Mtru,Agam,Dmel,Xtro,Drer,Mmus,Hsap \
        download prep iloci breakdown stats
```

Centroid (φ, σ) values were then computed for each value of δ for comparison.

```bash
fidibus-compact.py --workdir=data --centroid=2.25 --length=1000000 \
                  --iqnt=0.95 --gqnt=0.05 \
                  Scer Cele Crei Mtru Agam Dmel Xtro Drer Mmus Hsap \
    > phisigma-refr-centroids-delta500.tsv

fidibus-compact.py --workdir=data-delta300 --centroid=2.25 --length=1000000 \
                  --iqnt=0.95 --gqnt=0.05 \
                  Scer Cele Crei Mtru Agam Dmel Xtro Drer Mmus Hsap \
    > phisigma-refr-centroids-delta300.tsv

fidibus-compact.py --workdir=data-delta750 --centroid=2.25 --length=1000000 \
                  --iqnt=0.95 --gqnt=0.05 \
                  Scer Cele Crei Mtru Agam Dmel Xtro Drer Mmus Hsap \
    > phisigma-refr-centroids-delta750.tsv
```


## Random gene arrangement

To investigate whether gene clustering occurs more frequently than expected by chance, we computed random arrangements of genes and re-computed iLoci and associated statistics for comparison.

```bash
parallel --gnu --jobs=10 bash shuffle.sh {} ::: Scer Cele Crei Mtru Agam Dmel Xtro Drer Mmus Hsap
fidibus-milocus-summary.py --shuffled --workdir=species --outfmt=tex \
                          Scer Cele Crei Mtru Agam Dmel Xtro Drer Mmus Hsap
```

(nicely sped up thanks to O. Tange (2018): GNU Parallel 2018, Mar 2018, ISBN 9781387509881, [DOI https://doi.org/10.5281/zenodo.1146014](https://doi.org/10.5281/zenodo.1146014).)


The (phi, sigma) values of genome compactness were computed for the shuffled data for comparison with the observed data.
A single (phi, sigma) value was reported for each genome, computed as the average (centroid) of all (phi, sigma) values for that species.
For any genome with (phi, sigma) values whose distance from the centroid exceeds 2.25 times the average (phi, sigma) distance from the centroid, these outliers are removed and the centroid is recomputed.

```bash
fidibus-compact.py --shuffled --centroid=2.25 --length=1000000 --iqnt=0.95 --gqnt=0.05 \
                  Scer Cele Crei Mtru Agam Dmel Xtro Drer Mmus Hsap \
    > phisigma-refr-shuffled-centroids-delta500.tsv
```

## Figures

See [notebooks/make-F3a-SF6-SF7.ipynb](./notebooks/make-F3a-SF6-SF7.ipynb) for visualization of these data.
