## AdditionalFile2 for "iLoci: Robust evaluation of genome content and organization for provisional and mature genome assemblies": README_refr-genome-summary.md

# Summary of model organism genomes

We selected a diverse set of 10 model organisms to demonstrate the utility of
iLoci for providing a descriptive overview of genome content and organization.

## Data retrieval and processing

The `fidibus` script was used to perform the following tasks:

- retrieve genome assemblies and annotations from [NCBI RefSeq](http://www.ncbi.nlm.nih.gov/refseq/) (the `download` task)
- pre-process the data (the `prep` task)
- compute iLoci (the `iloci` task)
- collate sequences and annotations for various feature types (the `breakdown` task)
- compile tables of summary statistics over the data (the `stats` task)

You can copy/paste the relevant command from below or alternatively execute
the [run-fidibus-stats.sh](./wfscripts/run-fidibus-stats.sh) and
[run-fidibus-cleanup.sh](./wfscripts/run-fidibus-cleanup.sh) workflow bash
scripts in the [wfscripts](./wfscripts) directory.
Note that the scripts include also the work on the special interest genomes
described in the manuscript, documentation of which is not repeated here.

```bash
fidibus --workdir=data \
        --numprocs=10 \
        --refr=Scer,Cele,Crei,Mtru,Agam,Dmel,Xtro,Drer,Mmus,Hsap \
        download prep iloci breakdown stats
```

This command could take up to a couple of hours to run: go enjoy lunch or
read a paper while you wait!
You will want to adjust the `--numprocs` parameter based on the number of
available processors on your computer, as processing multiple genomes in
parallel will complete the task more quickly.
However, the mammalian genomes each take on the order of 30 minutes, so that
is about the best runtime that can be expected.

## Output data

The `fidibus` command creates a directory called `data` containing
subdirectories, each dedicated to a specific genome.
These subdirectories store intermediate and ancillary data files.
Of particular interest are the following:

- `Xxxx.iloci.gff3`: full annotation file, with iLoci specified and associated with their constituent gene features
- `Xxxx.miloci.gff3`: annotation file containing just iLocus features, with overlapping siLoci and niLoci merged into miLoci
- `Xxxx.yyyy.tsv`: statistics calculated on various data types (iLoci, miLoci, pre-mRNAs, mature mRNAs, exons, introns, coding sequences) and compiled into a tabular format for easy loading into R, Python, or similar data analysis environments

## iLocus summary

Summaries of iLocus composition for these genomes, corresponding to Tables 1-3
in the manuscript, were computed with the following commands (script
[wfscripts/make-Tables1-3.sh](./wfscripts/make-Tables1-3.sh)).

```bash
fidibus-ilocus-summary.py  --workdir=data --outfmt=tsv Scer Cele Crei Mtru Agam Dmel Xtro Drer Mmus Hsap
fidibus-pilocus-summary.py --workdir=data --outfmt=tsv Scer Cele Crei Mtru Agam Dmel Xtro Drer Mmus Hsap
fidibus-milocus-summary.py --workdir=data --outfmt=tsv Scer Cele Crei Mtru Agam Dmel Xtro Drer Mmus Hsap
```
