## AdditionalFile2 for "iLoci: Robust evaluation of genome content and organization for provisional and mature genome assemblies": shuffle.sh

```
#!/bin/bash
#

species=$1

locuspocus --verbose --delta=0 --skipends \
	--outfile=data/${species}/${species}.iloci.0d.gff3 \
	data/${species}/${species}.gff3

scripts/shuffle_iloci.py --seed=54321 \
	--out=data/${species}/${species}.shuffled.genes.gff3 \
	data/${species}/${species}.iloci.0d.gff3

locuspocus --verbose --delta=500 --skipends --cds \
	--namefmt="${species}ILCshuf-%06lu" \
	--outfile=data/${species}/${species}.iloci.shuffled.gff3 \
	data/${species}/${species}.shuffled.genes.gff3


# Analyzing iLoci:
#
xtractore --type=locus \
	--outfile=data/${species}/${species}.iloci.shuffled.fa \
	data/${species}/${species}.iloci.shuffled.gff3 \
	data/${species}/${species}.gdna.fa

fidibus-stats.py --species=${species} --iloci \
	data/${species}/${species}.iloci.shuffled.gff3 \
	data/${species}/${species}.iloci.shuffled.fa \
	data/${species}/${species}.iloci.shuffled.tsv


# Analyzing miLoci:
#
miloci.py < data/${species}/${species}.iloci.shuffled.gff3 \
	> data/${species}/${species}.miloci.shuffled.gff3

xtractore --type=locus \
	--outfile=data/${species}/${species}.miloci.shuffled.fa \
	data/${species}/${species}.miloci.shuffled.gff3 \
	data/${species}/${species}.gdna.fa

fidibus-stats.py --species=${species} --miloci \
	data/${species}/${species}.miloci.shuffled.gff3 \
	data/${species}/${species}.miloci.shuffled.fa \
	data/${species}/${species}.miloci.shuffled.tsv
```
