## AdditionalFile2 for "iLoci: Robust evaluation of genome content and organization for provisional and mature genome assemblies": shuffle_iloci.py

```
#!/usr/bin/env python
#
# Copyright (c) 2015   Daniel Standage <>
# Copyright (c) 2015   Indiana University
# All rights reserved.
#
# This file is released under the BSD 3-clause license.
#
# Redistribution and use in source and binary forms, with or without
# modification, are permitted provided that the following conditions are met:
#
#     * Redistributions of source code must retain the above copyright
#       notice, this list of conditions and the following disclaimer.
#     * Redistributions in binary form must reproduce the above copyright
#       notice, this list of conditions and the following disclaimer in the
#       documentation and/or other materials provided with the distribution.
#     * Neither the name of copyright holder nor the names of its contributors
#       may be used to endorse or promote products derived from this software
#       without specific prior written permission.
#
# THIS SOFTWARE IS PROVIDED BY THE COPYRIGHT HOLDERS AND CONTRIBUTORS "AS IS"
# AND ANY EXPRESS OR IMPLIED WARRANTIES, INCLUDING, BUT NOT LIMITED TO, THE
# IMPLIED WARRANTIES OF MERCHANTABILITY AND FITNESS FOR A PARTICULAR PURPOSE
# ARE DISCLAIMED. IN NO EVENT SHALL THE COPYRIGHT HOLDERS BE LIABLE FOR ANY
# DIRECT, INDIRECT, INCIDENTAL, SPECIAL, EXEMPLARY, OR CONSEQUENTIAL DAMAGES
# (INCLUDING, BUT NOT LIMITED TO, PROCUREMENT OF SUBSTITUTE GOODS OR SERVICES;
# LOSS OF USE, DATA, OR PROFITS; OR BUSINESS INTERRUPTION) HOWEVER CAUSED AND
# ON ANY THEORY OF LIABILITY, WHETHER IN CONTRACT, STRICT LIABILITY, OR TORT
# (INCLUDING NEGLIGENCE OR OTHERWISE) ARISING IN ANY WAY OUT OF THE USE OF THIS
# SOFTWARE, EVEN IF ADVISED OF THE POSSIBILITY OF SUCH DAMAGE.
# -----------------------------------------------------------------------------

from __future__ import print_function
import argparse
import numpy
import re
import sys


class Sequence(object):
    def __init__(self, seqreg_entry):
        seqreg_pattern = '^##sequence-region\s+(\S+)\s+(\d+)\s+(\d+)'
        srmatch = re.search(seqreg_pattern, seqreg_entry)
        assert srmatch, 'Unable to parse sequence-region: ' + seqregn_entry
        self.seqid = srmatch.group(1)
        self.start = int(srmatch.group(2))
        self.end = int(srmatch.group(3))

    def __len__(self):
        return self.end - self.start + 1

    def __repr__(self):
        return '##sequence-region   %s %d %d' % (self.seqid, self.start,
                                                 self.end)


class giLocus(object):
    def __init__(self, feature_entry):
        fields = feature_entry.split('\t')
        assert len(fields) == 9 and \
            fields[2] == 'locus' and \
            ';child_gene=' in fields[8]

        self.seqid = fields[0]
        self.start = int(fields[3])
        self.end = int(fields[4])
        self._entry = feature_entry
        assert self.start <= self.end
        idmatch = re.search('ID=([^;\n]+)', fields[8])
        if idmatch:
            self.locusid = idmatch.group(1)

    def __lt__(self, other):
        return self.locusid < other.locusid

    def __len__(self):
        return self.end - self.start + 1

    def __repr__(self):
        return '%s_%d-%d' % (self.seqid, self.start, self.end)

    def place(self, newstart):
        length = len(self)
        self.offset = newstart - self.start
        self.start = newstart
        self.end = newstart + length - 1


def load_giloci(instream):
    sequences = dict()
    giloci = dict()
    for line in instream:
        line = line.rstrip()
        if line.startswith('##sequence-region'):
            seq = Sequence(line)
            sequences[seq.seqid] = seq
            giloci[seq.seqid] = dict()
        elif '\tlocus\t' in line and ';child_gene=' in line:
            locus = giLocus(line)
            giloci[locus.seqid][locus.locusid] = locus
    return sequences, giloci


def shuffle_giloci(sequence, giloci):
    loci = sorted(list(giloci.values()))
    numpy.random.shuffle(loci)

    other = sum([len(x) for x in loci if x.seqid == sequence.seqid])
    #print('DEBUG seqid=%s seqlen=%d other=%d' % (sequence.seqid, len(sequence), other))
    n_open_slots = len(sequence) - other
    #print('DEBUG n_open_slots=%d' % n_open_slots)
    slots_taken = dict()
    collisions = 0
    for locus in loci:
        new_slot = numpy.random.randint(1, n_open_slots)
        while new_slot in slots_taken:
            collisions += 1
            new_slot = numpy.random.randint(1, n_open_slots + 1)
        slots_taken[new_slot] = locus

    slots = sorted(list(slots_taken))
    assert len(slots) == len(loci)
    cumulative_offset = 0
    for slot in slots:
        locus = slots_taken[slot]
        newstart = slot + cumulative_offset
        cumulative_offset += len(locus)
        locus.place(newstart)

    return collisions


def print_giloci(instream, outstream, giloci, sequences, length_threshold):
    offsets = dict()
    for line in instream:
        line = line.rstrip()
        fields = line.split('\t')
        if len(fields) != 9:
            print(line, file=outstream)
            continue

        seqid = fields[0]
        if len(sequences[seqid]) < length_threshold:
            continue

        feature_type = fields[2]
        if feature_type == 'locus':
            continue

        parentmatch = re.search('Parent=([^;\n]+)', fields[8])
        if not parentmatch:
            print(line, file=outstream)
            continue

        parentid = parentmatch.group(1)
        if parentid in giloci[seqid] or parentid in offsets:
            if parentid in giloci[seqid]:
                locus = giloci[seqid][parentid]
                offset = locus.offset
                fields[8] = re.sub(';Parent=[^;\n]+', '', fields[8])
            else:
                offset = offsets[parentid]
            newstart = int(fields[3]) + locus.offset
            newend = int(fields[4]) + locus.offset
            fields[3] = str(newstart)
            fields[4] = str(newend)
            line = '\t'.join(fields)
            #print('DEBUG line="%s"' % line, file=sys.stderr)
            #assert 'Parent' not in line, line
            idmatch = re.search('ID=([^;\n]+)', fields[8])
            if idmatch:
                feature_id = idmatch.group(1)
                offsets[feature_id] = offset

        print(line, file=outstream)


def parse_args():
    parser = argparse.ArgumentParser()
    parser.add_argument('-o', '--out', type=argparse.FileType('w'),
                        default=sys.stdout, help='Output file; default is '
                        'terminal (stdout)')
    parser.add_argument('-s', '--seed', type=int,
                        help='Seed for random number generator',
                        default=numpy.random.randint(0, sys.maxsize))
    parser.add_argument('-l', '--length', type=int, default=1000000,
                        help='minimum length filter; default=1000000 (1 Mb)')
    parser.add_argument('gff3', type=str, help='LocusPocus GFF3 file')
    return parser.parse_args()


def main():
    args = parse_args()
    print('Random seed: %d' % args.seed, file=sys.stderr)
    numpy.random.seed(args.seed)
    with open(args.gff3, 'r') as instream:
        sequences, giloci = load_giloci(instream)
    for sequence in sequences.values():
        if len(sequence) < args.length:
            continue
        seq_giloci = giloci[sequence.seqid]
        collisions = shuffle_giloci(sequence, seq_giloci)
    with open(args.gff3, 'r') as instream:
        print_giloci(instream, args.out, giloci, sequences, args.length)


if __name__ == '__main__':
    main()
```
