## AdditionalFile2 for "iLoci: Robust evaluation of genome content and organization for provisional and mature genome assemblies": configure-genomes.sh

```
#!/bin/bash
#
set -e
set -u
set -o pipefail


branches=(fungi invertebrate plant protozoa vertebrate_mammalian vertebrate_other)

for branch in ${branches[@]}
do
    mkdir ${branch}_configs
    lftp -e "find;quit" ftp://ftp.ncbi.nlm.nih.gov/genomes/refseq/${branch}/ | egrep "latest_assembly_versions/\w+" > ${branch}_latest.txt
    python make-cfg_files.py $branch
done
```
