## AdditionalFile2 for "iLoci: Robust evaluation of genome content and organization for provisional and mature genome assemblies": get-statistics.sh

```
#!/bin/bash
#
set -u

branches=(fungi invertebrate plant protozoa vertebrate_mammalian vertebrate_other)

for branch in ${branches[@]}
do
    species_list=($(cat ${branch}_parsed.txt))
    fidibus-ilocus-summary.py  --workdir=${branch}_work \
 	--outfmt=tsv ${species_list[@]} > ${branch}-ilocus-summaries.tsv
    fidibus-pilocus-summary.py  --workdir=${branch}_work \
	--outfmt=tsv ${species_list[@]} > ${branch}-pilocus-summaries.tsv
    fidibus-milocus-summary.py  --workdir=${branch}_work \
	--outfmt=tsv ${species_list[@]} > ${branch}-milocus-summaries.tsv
    fidibus-compact.py  --cfgdir=${branch}_configs --workdir=${branch}_work \
	--centroid=2.25 --length=1000000 --iqnt=0.95 --gqnt=0.05 ${species_list[@]} \
	> ${branch}_centroids.tsv
done
```
