## AdditionalFile2 for "iLoci: Robust evaluation of genome content and organization for provisional and mature genome assemblies": make-cfg_files.py

```
#!/usr/bin/env pythyon
#
import re
import sys

# We process lists *_latest.txt (derived from successful calls to the
# NCBI ftp site) one list at a time as provided by the argument to this
# script.

cfglist = [] # ... will contain the succesfully configured species labels

if len(sys.argv) < 2:
    print("Enter one branch name")
elif len(sys.argv) > 2: 
    print("Enter only one branch name")
else:
    branch = sys.argv[1]

p = re.compile("\d{3}.\d_") # Accessions of the form GCF_(digits).(digit) for the most part.
# Exceptions: GCF_(digits).(two digits)
p2 = re.compile("\d{3}.\d{2}_")  # Handles this exception

f = open(branch + '_latest.txt', 'r') 
for line in f:
    yaml = {}
    entry = line.strip().split('/')
    name = entry[1].split('_')
    info = entry[-1]
    match = p.search(info)
    if match is None: # Handling the above exception with p2
        match = p2.search(info)
    try:
        fname = name[0][0] + name[1][:3]
    except IndexError:
        fname = name[0][:4]
    if len(fname) == 4:
        yaml['file_name'] = fname
    else:
        continue
    try:
        yaml['species'] = name[0] + ' ' + name[1]
    except IndexError:
        yaml['species'] = name[0]
    yaml['accession'] = info[:match.start()+5]
    yaml['build'] = info[match.start()+6:]
    L = [yaml['file_name'] + ':\n', "    species: " + yaml['species'] + '\n',
         "    source: refseq \n","    branch: " + branch + '\n',
         "    accession: " + yaml['accession'] + '\n',
         "    build: " + yaml['build'] + '\n']
    cfglist.append(yaml['file_name'] + '\n')
    with open(branch + '_configs/' + yaml['file_name'] + ".yml",'w') as y:
        y.writelines(L) 
f.close()

# Write out the list of successfully configured species into file
# *_configured.txt:
#
with open(branch + '_configured.txt','w') as b:
    b.writelines(sorted(set(cfglist)))
```
