## AdditionalFile2 for "iLoci: Robust evaluation of genome content and organization for provisional and mature genome assemblies": make-SuppTable234_taxa_averages.py

```
#Reads in the three types of summary TSV files from fidibus on branches from NCBI. 

import pandas as pd

branches = ['fungi','invertebrate','vertebrate_mammalian','vertebrate_other',
            'plant','protozoa']
df = pd.read_csv('fungi-ilocus-summaries.tsv', sep = '\t')
ilocus_columns = df.columns
df = pd.read_csv('fungi-pilocus-summaries.tsv', sep = '\t')
pilocus_columns = df.columns
df = pd.read_csv('fungi-milocus-summaries.tsv', sep = '\t')
milocus_columns = df.columns
ilocus_data = []
pilocus_data = []
milocus_data = []

for branch in branches:
    df = pd.read_csv(branch + '-ilocus-summaries.tsv', sep = '\t')
    mean = df['Mb'].mean(skipna = True)
    medians = df.median(axis = 0, skipna = True, numeric_only = True)
    medians = [median for median in medians]
    averages = [branch, mean] + medians[1:]
    ilocus_data.append(averages)
    
    df = pd.read_csv(branch + '-pilocus-summaries.tsv', sep = '\t')
    means = list(df.mean(axis = 0, skipna = True, numeric_only = True))
    medians = list(df.median(axis = 0, skipna = True, numeric_only = True))
    medians[1] = medians[1] * 10e-7
    averages = [branch] + medians[:2] + [means[2], medians[3], means[4]] 
    pilocus_data.append(averages)
    
    df = pd.read_csv(branch + '-milocus-summaries.tsv', sep = '\t')
    # One column contains three quartiles, separated by commas. 
    # Split it into three different columns
    quartiles = df.GeneCountQuartiles.str.split(',', expand = True) 
    quartiles.columns = ['one','med','three'] 
    means = list(df.mean(axis = 0, skipna = True, numeric_only = True))
    medians = list(df.median(axis = 0, skipna = True, numeric_only = True))
    medians[1] = medians[1] * 10e-7
    averages = [branch] + medians[:2] + [means[2], quartiles.med.median(), 
                                         medians[3], means[4]]
    milocus_data.append(averages)

# Place the mean/median in the same format as the input data:
#
ilocus_avgs = pd.DataFrame(ilocus_data, columns = ilocus_columns)
pilocus_avgs = pd.DataFrame(pilocus_data, columns = pilocus_columns)
milocus_avgs = pd.DataFrame(milocus_data, columns = milocus_columns)

pilocus_avgs.to_csv('SLB21-SuppTable2-pilocus_averages.tsv', sep = '\t', index = False)
milocus_avgs.to_csv('SLB21-SuppTable3-milocus_averages.tsv', sep = '\t', index = False)
ilocus_avgs.to_csv('SLB21-SuppTable4-ilocus_averages.tsv', sep = '\t', index = False)
```
