## AdditionalFile2 for "iLoci: Robust evaluation of genome content and organization for provisional and mature genome assemblies": README.md

# Genome summaries for NCBI-retrievable genomes from specific taxanomic branches

We first retrieve the hierarchical directory structure for the taxa of interest.
For example,

```bash
lftp -e "find;quit" ftp://ftp.ncbi.nlm.nih.gov/genomes/refseq/invertebrate/ > invertebrate_temp.txt
```

The output lists the main directories for all species, including subdirectories for all available genome versions.
We pull out only the latest assembly version as representative for each species:

```bash
egrep "latest_assembly_versions/\w+" invertebrate_temp.txt > invertebrate_latest.txt
```

The python script [make-cfg_files.py](https://github.com/BrendelGroup/iLoci_SLB21/blob/master/work/taxa/make-cfg_files.py) takes a file like _branch\_latest.txt_ as input to generate _*.yml_ configuration files for input to _fidibus_.
Species are abbreviated by a 4-letter code (e.g., Apis for "_Apis mellifera_").
If there are multiple directories in the download hierarchy that would lead to the same abbreviation, then former configuration files are over-written.
As we are only interested in samples of species from different branches of the taxonomy, this procedure is adequate.

The script [configure-genomes.sh](https://github.com/BrendelGroup/iLoci_SLB21/blob/master/work/taxa/configure-genomes.sh) was used to generate all the data presented in our paper.
This was run at the beginning of May, 2020.
A re-run would likely produce slightly different results based on newly deposited genome assemblies and annotations.

## Analysis
Fidibus analysis was run on each successfully configured species to produce iLoci summaries.
Cases in which fidibus exited with an error code were ignored.
Such failures are mostly due to non-standard GFF3 annotation files.

Running the script [run-fidibus-all.sh](https://github.com/BrendelGroup/iLoci_SLB21/blob/master/work/taxa/run-fidibus-all.sh) followed by [get-statistics.sh](https://github.com/BrendelGroup/iLoci_SLB21/blob/master/work/taxa/get-statistics.sh) will produce the analysis presented in the paper.
