## AdditionalFile2 for "iLoci: Robust evaluation of genome content and organization for provisional and mature genome assemblies": run-fidibus-all.sh

```
#!/bin/bash
#

#Note: We assume that you have sufficient processors to run all branches
#      separately in the background.

mkdir logfiles
branches=(fungi invertebrate plant protozoa vertebrate_mammalian vertebrate_other)

for branch in ${branches[@]}
do
    ./run-fidibus-branch.sh ${branch} >& logfiles/log_${branch} &
done
wait
```
