## AdditionalFile2 for "iLoci: Robust evaluation of genome content and organization for provisional and mature genome assemblies": run-fidibus-branch.sh

```
#!/bin/bash
#
set -u

branch=$1

species_list=($(cat "$branch"_configured.txt))
mkdir ${branch}_work
for species in ${species_list[@]}
do
        if fidibus --cfgdir=${branch}_configs --workdir=${branch}_work --refr=${species} \
			download prep iloci breakdown stats \
			>& logfiles/${branch}_${species}.log
        then
            echo $species >> ${branch}_parsed.txt
        fi
        find ${branch}_work/${species} -type f -not -name "${species}*.gff3" -and -not -name "${species}*.tsv" -exec rm {} \;
done
omm -2 -3 ${branch}_configured.txt ${branch}_parsed.txt > ${branch}_failed.txt
```
