## AdditionalFile2 for "iLoci: Robust evaluation of genome content and organization for provisional and mature genome assemblies": make-figures.sh

```
mv figures figuresORIG
mkdir figures

jupyter nbconvert --to notebook --execute --inplace ./notebooks/make-F3a-SF6-SF7.ipynb 
(cd taxa && jupyter nbconvert --to notebook --execute --inplace ../notebooks/make-F3b.ipynb && cd ..)
jupyter nbconvert --to notebook --execute --inplace ./notebooks/make-F4a-F4b-SF5a.ipynb
jupyter nbconvert --to notebook --execute --inplace ./notebooks/make-F5-F6.ipynb
jupyter nbconvert --to notebook --execute --inplace ./notebooks/make-SF1.ipynb 
jupyter nbconvert --to notebook --execute --inplace \
	--ExecutePreprocessor.timeout=600 ./notebooks/make-SF2.ipynb 
jupyter nbconvert --to notebook --execute --inplace ./notebooks/make-SF3.ipynb 
jupyter nbconvert --to notebook --execute --inplace ./notebooks/make-SF4.ipynb 
jupyter nbconvert --to notebook --execute --inplace ./notebooks/make-SF5b.ipynb 
jupyter nbconvert --to notebook --execute --inplace ./notebooks/make-SF5c-SF8.ipynb 
jupyter nbconvert --to notebook --execute --inplace ./notebooks/make-SF5d.ipynb 
```
