## AdditionalFile2 for "iLoci: Robust evaluation of genome content and organization for provisional and mature genome assemblies": make-SuppTable5.sh

```
#!/bin/bash
#

echo "Arabidopsis thaliana"	 > tables/SLB21-SuppTable5-genes.txt
echo ""	>> tables/SLB21-SuppTable5-genes.txt
echo "Number of genes annotated in TAIR 6:"	>> tables/SLB21-SuppTable5-genes.txt
egrep "	gene	" comparisons/species/Att6/Att6.gff3 | wc -l	>> tables/SLB21-SuppTable5-genes.txt
echo "... protein_coding:"	>> tables/SLB21-SuppTable5-genes.txt
egrep "	gene	" comparisons/species/Att6/Att6.gff3 | egrep -c "protein_coding"	>> tables/SLB21-SuppTable5-genes.txt
echo "... not protein_coding:"	>> tables/SLB21-SuppTable5-genes.txt
egrep "	gene	" comparisons/species/Att6/Att6.gff3 | egrep -v "protein_coding" | wc -l	>> tables/SLB21-SuppTable5-genes.txt
	>> tables/SLB21-SuppTable5-genes.txt
echo ""	>> tables/SLB21-SuppTable5-genes.txt
echo "Number of genes annotated in TAIR 10:"	>> tables/SLB21-SuppTable5-genes.txt
egrep "	gene	" comparisons/species/Atha/Atha.gff3 | wc -l	>> tables/SLB21-SuppTable5-genes.txt
echo "... protein_coding:"	>> tables/SLB21-SuppTable5-genes.txt
egrep "	gene	" comparisons/species/Atha/Atha.gff3 | egrep -c "protein_coding"	>> tables/SLB21-SuppTable5-genes.txt
echo "... not protein_coding:"	>> tables/SLB21-SuppTable5-genes.txt
egrep "	gene	" comparisons/species/Atha/Atha.gff3 | egrep -v "protein_coding" | wc -l	>> tables/SLB21-SuppTable5-genes.txt
	>> tables/SLB21-SuppTable5-genes.txt
	>> tables/SLB21-SuppTable5-genes.txt
echo ""	>> tables/SLB21-SuppTable5-genes.txt
echo ""	>> tables/SLB21-SuppTable5-genes.txt
echo "Apis mellifera"	>> tables/SLB21-SuppTable5-genes.txt
echo ""	>> tables/SLB21-SuppTable5-genes.txt
echo "Number of genes annotated in Amel4.5:"	>> tables/SLB21-SuppTable5-genes.txt
egrep "	gene	" comparisons/species/Am45/Am45.gff3 | wc -l	>> tables/SLB21-SuppTable5-genes.txt
echo "... protein_coding:"	>> tables/SLB21-SuppTable5-genes.txt
egrep "	gene	" comparisons/species/Am45/Am45.gff3 | egrep -c "protein_coding"	>> tables/SLB21-SuppTable5-genes.txt
echo "... not protein_coding:"	>> tables/SLB21-SuppTable5-genes.txt
egrep "	gene	" comparisons/species/Am45/Am45.gff3 | egrep -v "protein_coding" | wc -l	>> tables/SLB21-SuppTable5-genes.txt
	>> tables/SLB21-SuppTable5-genes.txt
echo ""	>> tables/SLB21-SuppTable5-genes.txt
echo "Number of genes annotated in AmelHAv3.1:"	>> tables/SLB21-SuppTable5-genes.txt
egrep "	gene	" comparisons/species/Amel/Amel.gff3 | wc -l	>> tables/SLB21-SuppTable5-genes.txt
echo "... protein_coding:"	>> tables/SLB21-SuppTable5-genes.txt
egrep "	gene	" comparisons/species/Amel/Amel.gff3 | egrep -c "protein_coding"	>> tables/SLB21-SuppTable5-genes.txt
echo "... not protein_coding:"	>> tables/SLB21-SuppTable5-genes.txt
egrep "	gene	" comparisons/species/Amel/Amel.gff3 | egrep -v "protein_coding" | wc -l	>> tables/SLB21-SuppTable5-genes.txt
echo ""	>> tables/SLB21-SuppTable5-genes.txt
echo ""	>> tables/SLB21-SuppTable5-genes.txt
echo ""	>> tables/SLB21-SuppTable5-genes.txt

fidibus-ilocus-summary.py  --workdir=comparisons/species --outfmt=tex \
        Att6 Atha Am45 Amel  > tables/SLB21-SuppTable5-iloci.tex
```
