## AdditionalFile2 for "iLoci: Robust evaluation of genome content and organization for provisional and mature genome assemblies": make-Tables1-3.sh

```
#!/bin/bash
#

# Write summary iLocus information into tab-delimited files in the tables
# directory:
#
mkdir tables

# ... generating *.tsv files for the record:
#
fidibus-ilocus-summary.py  --workdir=data --outfmt=tsv \
	Scer Cele Crei Mtru Agam Dmel Xtro Drer Mmus Hsap Vcar Pdom Dpul Mvit  > tables/ilocus-delta500-summary.tsv
fidibus-pilocus-summary.py --workdir=data --outfmt=tsv \
	Scer Cele Crei Mtru Agam Dmel Xtro Drer Mmus Hsap Vcar Pdom Dpul Mvit  > tables/pilocus-delta500-summary.tsv
fidibus-milocus-summary.py --workdir=data --outfmt=tsv \
	Scer Cele Crei Mtru Agam Dmel Xtro Drer Mmus Hsap Vcar Pdom Dpul Mvit  > tables/milocus-delta500-summary.tsv

# ... generating *.tex files for the manuscript:
#
fidibus-ilocus-summary.py  --workdir=data --outfmt=tex \
	Scer Cele Crei Mtru Agam Dmel Xtro Drer Mmus Hsap Vcar Pdom Dpul Mvit  > tables/SLB21-Table1-iloci.tex
fidibus-pilocus-summary.py --workdir=data --outfmt=tex \
	Scer Cele Crei Mtru Agam Dmel Xtro Drer Mmus Hsap Vcar Pdom Dpul Mvit  > tables/SLB21-Table2-piloci.tex
fidibus-milocus-summary.py --workdir=data --outfmt=tex \
	Scer Cele Crei Mtru Agam Dmel Xtro Drer Mmus Hsap Vcar Pdom Dpul Mvit  > tables/SLB21-Table3-miloci.tex
```
