## AdditionalFile2 for "iLoci: Robust evaluation of genome content and organization for provisional and mature genome assemblies": run-explore-compactness-othg.sh

```
#!/bin/bash
#


# Calculate the (phi,sigma) compactness variables for Vcar and related genomes:
#
fidibus-compact.py --workdir=data --length=1000000 \
		--iqnt=0.95 --gqnt=0.05 \
		Vcar Crei Cvar Csub Mcom Oluc \
    > tables/phisigma-Vcar.tsv


# Calculate the (phi,sigma) compactness variables for Pdom and related genomes:
#
fidibus-compact.py --workdir=data --length=1000000 \
		--iqnt=0.95 --gqnt=0.05 \
		Amel Agam Aech Dmel Pdom Nvit \
    > tables/phisigma-Pdom-min1Mb.tsv
fidibus-compact.py --workdir=data --length=2000000 \
		--iqnt=0.95 --gqnt=0.05 \
		Amel Agam Aech Dmel Pdom Nvit \
    > tables/phisigma-Pdom-min2Mb.tsv
fidibus-compact.py --workdir=data --length=3000000 \
		--iqnt=0.95 --gqnt=0.05 \
		Amel Agam Aech Dmel Pdom Nvit \
    > tables/phisigma-Pdom-min3Mb.tsv


# Calculate the (phi,sigma) compactness variables for Dpul and related genomes:
#
fidibus-compact.py --workdir=data --length=2000000 \
		--iqnt=0.95 --gqnt=0.05 \
		Amel Agam Dmel Dpul Tcas Turt \
    > tables/phisigma-Dpul.tsv


# Calculate the (phi,sigma) compactness variables for Mvit and related genomes:
#
fidibus-compact.py --workdir=data --length=1000000 \
		--iqnt=0.95 --gqnt=0.05 \
		Acen Clan Lcan Hsap Mmus Mvit \
    > tables/phisigma-Mvit.tsv
```
