## AdditionalFile2 for "iLoci: Robust evaluation of genome content and organization for provisional and mature genome assemblies": run-explore-compactness-refr.sh

```
#!/bin/bash
#

# Calculate the (phi,sigma) compactness variables for the model genomes:
#
fidibus-compact.py --workdir=data --length=1000000 \
                  --iqnt=0.95 --gqnt=0.05 \
                  Scer Cele Crei Mtru Agam Dmel Xtro Drer Mmus Hsap \
    > tables/phisigma-refr.tsv


# Explore the dependence of iLoci breakdown results on the extension parameter delta
# by re-running the prior analyses (with default value delta=500) with delta=300:
#
mkdir data-delta300
for species in  Scer Cele Crei Mtru Agam Dmel Xtro Drer Mmus Hsap
do
	mkdir data-delta300/$species
	ln -s ../../data/$species/$species.gdna.fa     data-delta300/$species/$species.gdna.fa
	ln -s ../../data/$species/$species.gff3        data-delta300/$species/$species.gff3
	ln -s ../../data/$species/$species.all.prot.fa data-delta300/$species/$species.all.prot.fa
done

fidibus --workdir=data-delta300 \
        --numprocs=10 \
        --delta=300 \
        --refr=Scer,Cele,Crei,Mtru,Agam,Dmel,Xtro,Drer,Mmus,Hsap \
        iloci breakdown stats

# ... and now delta=750:
#
mkdir data-delta750
for species in  Scer Cele Crei Mtru Agam Dmel Xtro Drer Mmus Hsap
do
	mkdir data-delta750/$species
	ln -s ../../data/$species/$species.gdna.fa     data-delta750/$species/$species.gdna.fa
	ln -s ../../data/$species/$species.gff3        data-delta750/$species/$species.gff3
	ln -s ../../data/$species/$species.all.prot.fa data-delta750/$species/$species.all.prot.fa
done

fidibus --workdir=data-delta750 \
        --numprocs=10 \
        --delta=750 \
        --refr=Scer,Cele,Crei,Mtru,Agam,Dmel,Xtro,Drer,Mmus,Hsap \
        iloci breakdown stats


# Derive the centroids of (phi,sigma) value pairs for plotting;
#
fidibus-compact.py --workdir=data --centroid=2.25 --length=1000000 \
                  --iqnt=0.95 --gqnt=0.05 \
                  Scer Cele Crei Mtru Agam Dmel Xtro Drer Mmus Hsap \
    > tables/phisigma-refr-centroids-delta500.tsv

fidibus-compact.py --workdir=data-delta300 --centroid=2.25 --length=1000000 \
                  --iqnt=0.95 --gqnt=0.05 \
                  Scer Cele Crei Mtru Agam Dmel Xtro Drer Mmus Hsap \
    > tables/phisigma-refr-centroids-delta300.tsv

fidibus-compact.py --workdir=data-delta750 --centroid=2.25 --length=1000000 \
                  --iqnt=0.95 --gqnt=0.05 \
                  Scer Cele Crei Mtru Agam Dmel Xtro Drer Mmus Hsap \
    > tables/phisigma-refr-centroids-delta750.tsv


# Compare observed compactness with calculations on randomized genomes:
#
parallel --gnu --jobs=10 bash scripts/shuffle.sh {} ::: \
	Scer Cele Crei Mtru Agam Dmel Xtro Drer Mmus Hsap

fidibus-milocus-summary.py --workdir=data --shuffled --outfmt=tex \
	Scer Cele Crei Mtru Agam Dmel Xtro Drer Mmus Hsap \
	> tables/SLB21-SuppTable1-miloci.tex

fidibus-compact.py --workdir=data --shuffled \
	--centroid=2.25 --length=1000000 --iqnt=0.95 --gqnt=0.05 \
	Scer Cele Crei Mtru Agam Dmel Xtro Drer Mmus Hsap \
	> tables/phisigma-refr-shuffled-centroids-delta500.tsv
```
