## AdditionalFile2 for "iLoci: Robust evaluation of genome content and organization for provisional and mature genome assemblies": run-explore-gene-orientation.sh

```
#!/bin/bash
#

# Explore the dependence of iLoci breakdown results on the extension parameter delta
# by re-running the prior analyses (with default value delta=500) with delta=0:
#
mkdir data-delta0
for species in  Scer Cele Crei Mtru Agam Dmel Xtro Drer Mmus Hsap
do
	mkdir data-delta0/$species
	ln -s ../../data/$species/$species.gdna.fa     data-delta0/$species/$species.gdna.fa
	ln -s ../../data/$species/$species.gff3        data-delta0/$species/$species.gff3
	ln -s ../../data/$species/$species.all.prot.fa data-delta0/$species/$species.all.prot.fa
done

fidibus --workdir=data-delta0 \
        --numprocs=10 \
        --delta=0 \
        --refr=Scer,Cele,Crei,Mtru,Agam,Dmel,Xtro,Drer,Mmus,Hsap \
        iloci breakdown stats

# Compare observed results with calculations on randomized genomes:
#
for species in  Scer Cele Crei Mtru Agam Dmel Xtro Drer Mmus Hsap
do
	locuspocus --verbose --delta=0 --skipends --cds \
        	--namefmt="${species}ILCshuf-%06lu" \
        	--outfile=data-delta0/${species}/${species}.iloci.shuffled.gff3 \
        	data/${species}/${species}.shuffled.genes.gff3

	fidibus-stats.py --species=${species} --iloci \
        	data/${species}/${species}.iloci.shuffled.gff3 \
        	data/${species}/${species}.iloci.shuffled.fa \
        	data-delta0/${species}/${species}.iloci.shuffled.tsv
done
```
