## AdditionalFile2 for "iLoci: Robust evaluation of genome content and organization for provisional and mature genome assemblies": run-fidibus-cleanup.sh

```
#!/bin/bash
#

# We retain only the output files from run-fidibus-stats.sh that are necessary
# for our subsequent analyses:
#
for species in  Scer Cele Crei Mtru Agam Dmel Xtro Drer Mmus Hsap \
	Vcar Apro Crei Csub Cvar Mcom Mpus Oluc Otau \
	Atha Bdis Osat \
	Pdom Aech Agam Amel Bter Cflo Dmel Hsal Nvit Pcan \
	Mvit Tcas Turt
do
  fidibus \
	--workdir=data \
	--keep .gdna.fa .gff3 .prot.fa .protids.txt \
	--refr=$species \
	cleanup
done

fidibus --workdir=data \
	--keep .gdna.fa .gff3 .prot.fa .protids.txt \
        --local \
        --label=Dpul \
        --gdna=downloads/Daphnia_pulex.fasta.gz \
        --gff3=downloads/FrozenGeneCatalog20110204.gff3.gz \
        --prot=downloads/FrozenGeneCatalog20110204.proteins.fasta.gz \
        cleanup

\rm data/*/*.fna.gz data/*/*.gff.gz data/*/*.faa.gz

# Let's also put the error warnings into a subdirectory for safekeeping:
#
mkdir logfiles
\mv err* logfiles
```
