## AdditionalFile2 for "iLoci: Robust evaluation of genome content and organization for provisional and mature genome assemblies": run-fidibus-stats.sh

```
#!/bin/bash
#
NUMPROCS=10

# The fidibus commands are first illustrated for the model organisms and
# then run on the remaining data sets analyzed in the paper.

# 1) Run the download/prep steps individually per species in the background,
# capturing any warning messages concerning the GFF3 input:
#
for species in  Scer Cele Crei Mtru Agam Dmel Xtro Drer Mmus Hsap
do
  fidibus  --workdir=data  --refr=$species  download prep  >& err_$species &
done
wait

# 2) The following steps are run using the multiprocessing implementation in
# fidibus:
#
fidibus   --workdir=data  --numprocs=${NUMPROCS} \
	--refr=Scer,Cele,Crei,Mtru,Agam,Dmel,Xtro,Drer,Mmus,Hsap \
	iloci breakdown stats   >& err_all1


# Now we work on the remaining data sets:
#
# (i) Volvox carteri and other Chlorophyta:
#
for species in  Vcar Apro Crei Csub Cvar Mcom Mpus Oluc Otau
do
  fidibus  --workdir=data  --refr=$species  download prep  >& err_$species &
done
wait

fidibus   --workdir=data  --numprocs=${NUMPROCS} \
	--refr=Vcar,Apro,Crei,Csub,Cvar,Mcom,Mpus,Oluc,Otau \
	iloci breakdown stats   >& err_all2


# (ii) More plants:
#
for species in  Atha Bdis Osat
do
  fidibus  --workdir=data  --refr=$species  download prep  >& err_$species &
done
wait

fidibus   --workdir=data  --numprocs=${NUMPROCS} \
	--refr=Atha,Bdis,Osat \
	iloci breakdown stats   >& err_all3


# (iii) Polistes dominula and other Hymenoptera:
#
for species in  Pdom Aech Agam Amel Bter Cflo Dmel Hsal Nvit Pcan 
do
  fidibus  --workdir=data  --refr=$species  download prep  >& err_$species &
done
wait

fidibus   --workdir=data  --numprocs=${NUMPROCS} \
	--refr=Pdom,Aech,Agam,Amel,Bter,Cflo,Dmel,Hsal,Nvit,Pcan \
	iloci breakdown stats   >& err_all4


# (iv) More:
#
fidibus --cfgdir=genome_configs --workdir=data --refr=Dpul \
        download prep iloci breakdown stats	>& err_Dpul

for species in  Acen Clan Lcan Mvit Tcas Turt
do
  fidibus  --workdir=data  --refr=$species  download prep  >& err_$species &
done
wait

fidibus   --workdir=data  --numprocs=${NUMPROCS} \
	--refr=Acen,Clan,Lcan,Mvit,Tcas,Turt \
	iloci breakdown stats   >& err_all5
```
